# Modulation of macrophage inflammatory function through selective inhibition of the epigenetic reader protein SP140

**DOI:** 10.1101/2020.08.10.239475

**Authors:** Mohammed Ghiboub, Jan Koster, Peter D. Craggs, Andrew Y.F. Li Yim, Anthony Shillings, Sue Hutchinson, Ryan P. Bingham, Kelly Gatfield, Ishtu L. Hageman, Gang Yao, Heather P. O’Keefe, Aaron Coffin, Amish Patel, Lisa A. Sloan, Darren J. Mitchell, Laurent Lunven, Robert J. Watson, Christopher E. Blunt, Lee A. Harrison, Gordon Bruton, Umesh Kumar, Natalie Hamer, John R. Spaull, Danny A. Zwijnenburg, Olaf Welting, Theodorus B.M. Hakvoort, Johan van Limbergen, Peter Henneman, Rab K. Prinjha, Menno PJ. de Winther, Nicola R. Harker, David F. Tough, Wouter J. de Jonge

**Affiliations:** Tytgat Institute for Liver and Intestinal Research, Amsterdam Gastroenterology & Metabolism, Amsterdam University Medical Centers, University of Amsterdam, the Netherlands; Department of Oncogenomics, Amsterdam University Medical Centers, University of Amsterdam and Cancer Center Amsterdam, Amsterdam, Netherlands; Immuno-Epigenetics, Adaptive Immunity Research Unit, GlaxoSmithKline, Medicines Research Centre, Stevenage, United Kingdom; Medicine Design, Medicinal Science and Technology, GlaxoSmithKline, Stevenage, United Kingdom; Department of Pediatrics, Division of Pediatric Gastroenterology & Nutrition, Emma Children’s Hospital, Amsterdam University Medical Centers, University of Amsterdam, the Netherlands; Department of Clinical Genetics, Genome Diagnostics Laboratory, Amsterdam Reproduction & Development, Amsterdam University Medical Centers, University of Amsterdam, the Netherlands; GlaxoSmithKline, Cambridge, Massachusetts, USA; Constellation Pharmaceuticals, Cambridge, Massachusetts, USA; WuXi AppTec, Cambridge, Massachusetts, USA; Department of Medical Biochemistry, Amsterdam University Medical Centers, University of Amsterdam, Amsterdam, Netherlands; Institute for Cardiovascular Prevention (IPEK), Munich, Germany; Adaptive Immunity Research Unit, GlaxoSmithKline, Medicines Research Centre, Stevenage, United Kingdom; Department of Surgery, University of Bonn, Bonn, Germany

## Abstract

Speckled 140 KDa (SP140) is a nuclear body protein, mainly expressed in immune cells, which contains multiple domains suggestive of an epigenetic reader function; namely a bromodomain, a PHD domain and a SAND domain. Single nucleotide polymorphisms and epigenetic modifications in the *SP140* locus have been linked to autoimmune and inflammatory diseases including Crohn’s disease (CD). However, little is known about the cellular function of SP140; this is due in part to the fact that, unlike for other many other epigenetic proteins, no small molecule inhibitors have been available to investigate the biological role of SP140. We report the discovery of the first small molecule SP140 inhibitor (GSK761) and utilize this to elucidate SP140 function in innate immune cells. We show that SP140 is highly expressed in CD68^+^ CD mucosal macrophages and in *in vitro-*generated inflammatory macrophages. SP140 inhibition through GSK761 reduced monocyte differentiation into inflammatory macrophages and lipopolysaccharide (LPS)-induced inflammatory activation, whilst inducing the generation of CD206^+^ regulatory macrophages that mark anti-TNF remission induction in CD patients. ChIP-seq analyses revealed that SP140 preferentially occupies transcriptional start sites (TSS) in inflammatory macrophages, with enrichment at gene loci encoding pro-inflammatory cytokines/chemokines and inflammatory pathways. GSK761 specifically reduced SP140 binding and thereby expression of SP140-dependent downstream inflammatory genes. Notably, in CD14^+^ macrophages isolated from CD intestinal-mucosa, GSK761 inhibited the spontaneous expression of cytokines, including *TNF*. Together, this study identifies SP140 as a druggable epigenetic reader and therapeutic target for CD.

## Introduction

Although various therapeutic strategies provide benefit in Crohn’s disease (CD), the unmet need remains high with only approximately 30% of patients maintaining long term remission^1,2,3,4^. Intestinal macrophages, which arise mainly from blood-derived monocytes, play a central role in maintaining gut homeostasis in health, but are also implicated as key contributors to CD^5,6^. These opposing activities are linked to the capacity of macrophages to take on different functional properties in response to environmental stimuli. Epigenetic processes play an important role in the regulation of macrophage differentiation and gene expression, mediated by DNA methylation and histone post-translational modification^7,8^. As such, epigenetic mechanisms mediated through the activity of specific histone acetylases and deacetylases activity are implicated in CD pathogenesis^9,10,11^.

Speckled 140 KDa (SP140) is a nuclear body protein^12,13^ predominantly expressed in immune cells^14,15^. Genome-wide association studies identified an association between single nucleotide polymorphisms in *SP140* and CD^16^ while rare *SP140*-associated variants have been identified in pediatric CD patients^17^. Further, we identified two differentially hyper-methylated positions at the *SP140* locus in blood cells of CD patients^18^, suggesting aberrant regulation of its expression. Although little is known about how SP140 may be functionally linked to CD, a recent study reported that *SP140* knockdown in *in vitro*-generated inflammatory macrophages results a decrease in levels of pro-inflammatory cytokines^10^. Lower expression of *SP140* in intestinal biopsies was correlated with an improved response to anti-TNF therapy in CD patients^10^. Conversely, in biopsies from those who showed resistance to anti-TNF therapy, SP140 was highly expressed^10^. Together, these previous findings suggest a role for SP140 in CD pathogenesis. SP140 protein harbors both a bromodomain (Brd) and a PHD finger^10,15^. Since Brd and PHD domains are capable of binding to histones, this suggests that SP140 might function as an epigenetic reader. Readers are recruited to chromatin possessing specific epigenetic ‘marks’ and regulate gene expression through mechanisms that modulate DNA accessibility to transcription factors (TFs) and the transcriptional machinery^19^. Brds are tractable to small molecule antagonists, with selective inhibitors of several Brd-containing proteins (BCPs) reported^20^. Such compounds have helped elucidate the function of their BCP targets; for instance, multiple inhibitors targeting BRD2, BRD3 and BRD4 have been used to demonstrate the role of these proteins in selectively regulating gene expression in the context of cancer and inflammation^9,21,22^. To date, no inhibitors targeting SP140 have been reported.

Here, we describe the discovery of the first specific SP140 inhibitor and utilize this compound to investigate the functional relevance of SP140 in CD pathogenesis, focusing on its role in macrophages. We show that SP140 functions as an epigenetic reader in controlling the generation of an inflammatory macrophage phenotype, and in regulating the expression of pro-inflammatory and CD-associated genes. We show that GSK761 inhibits SP140 binding to an array of inflammatory genes and demonstrate its efficacy in CD colon macrophages *ex vivo*, presenting evidence for SP140 as a target for regulating inflammation in CD. In addition, this study suggests that SP140 inhibition may also serve to support anti-TNF therapy in non-responder CD patients.

## Results

### Elevated SP140 expression in inflammatory diseases and mucosal macrophages of CD patients

*SP140* gene expression in a variety of cells and tissues was analyzed using an in-house GSK microarray profiler. *SP140* was predominantly expressed in blood, CD8^+^ and CD4^+^ T cells, monocytes and in secondary immune organs such as lymph node and spleen (Fig. 1a). We then investigated SP140 expression in white blood cells (WBC) and intestinal tissue of IBD patients versus controls. While *SP140* gene expression in WBC was comparable in IBD and healthy controls (Fig. 1b), SP140 gene and protein expression were found to be increased in inflamed colonic mucosa of both CD and UC patients (Fig. 1c,d and S1). Notably, an increase in CD68^+^SP140^+^ but not CD68^+^SP140^-^ macrophages was observed in CD inflamed colonic mucosa compared to normal control mucosa (Fig. 1e,f). Elevated SP140 expression was also found in other inflammatory conditions including in WBC of systemic lupus erythematosus and rheumatoid arthritis patients (Fig. 1b), and in inflamed tissues of appendicitis, sarcoidosis, psoriatic arthritis, rheumatoid arthritis, Hashimoto’s thyroiditis and Sjogren’s syndrome patients (Fig. S1). High expression of SP140 was also observed in chronic lymphocytic leukemia (Fig. 1b).

**Figure 1:**
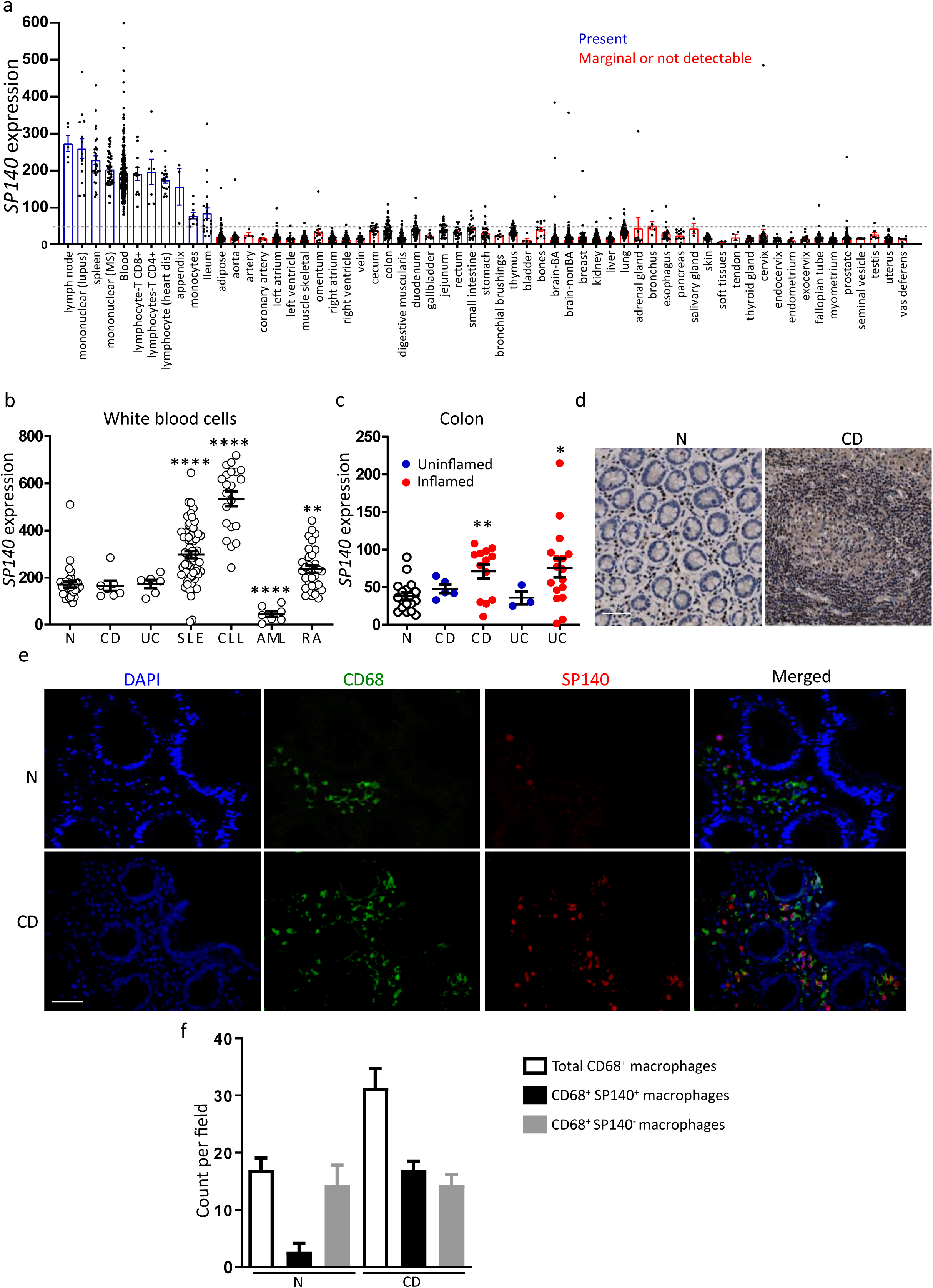
SP140 is highly associated with inflammatory diseases and mucosal CD68^+^ macrophages of CD patients. **(a)** *SP140* gene expression in human tissues and cell types. **(b)** *SP140* gene expression in white blood cells of normal healthy controls (N), Crohn’s disease (CD), ulcerative colitis (CD), systemic lupus erythematosus (SLE), chronic lymphocytic leukemia (CCL), acute myeloid leukemia (AML) and rheumatoid arthritis (RA) patients. **(c)** *SP140* gene expression in human colon tissue obtained from N, inflamed and non-inflamed CD and UC colonic tissues. Data was collected from in-house GSK microarray profiler. **(d)** Immunohistochemistry of SP140 protein in colon tissue obtained from N and inflamed CD tissue, scale bar: 50 µm. **(e)** Immunofluorescence staining of DAPI (blue), CD68 (green) and SP140 (red) in N or inflamed CD colon tissue, scale bar: 100 µm. **(f)** Mucosal cell count per visual field of total CD68^+^ macrophages, SP140^+^ CD68^+^ macrophages and SP140^-^ CD68^+^ macrophages in N and inflamed CD tissue (n=3). Statistical significance is indicated as follows: \**P* < 0.05, \*\**P* < 0.01, \*\*\*\**P* < 0.0001.

Next, we analyzed data set from a publicly available single-cell RNA-sequencing experiment^23^ of inflamed and uninflamed ileal tissue (biopsies) obtained from CD patients. Unsupervised clustering analysis identified 22 cluster blocks (Fig. 2a). Based on the expression of *CD14, CD16* (*FCGR3A), CD64 (FCGR1A), CD68, CD163* and *CD206 (MRC1)*, we identified cluster 6 as containing monocytes, macrophages and dendritic cells (Fig. 2b,c), representing mononuclear phagocytes. Inflamed ileal biopsies contained more cells in cluster 6 as compared to uninflamed biopsies (Fig. S2). Interestingly, within cluster 6 the number of *SP140*-expressing cells was higher in inflamed ileal biopsies as well (Fig. 2d). In addition, the percentage of SP140 expressing cells in cluster 6 was higher in inflamed compared with uninflamed tissue, which coincides with an increased number of *CD68*-, *CD64*-*CD14-* and *CD16*-expressing cells (Fig. 2e).

**Figure 2.**
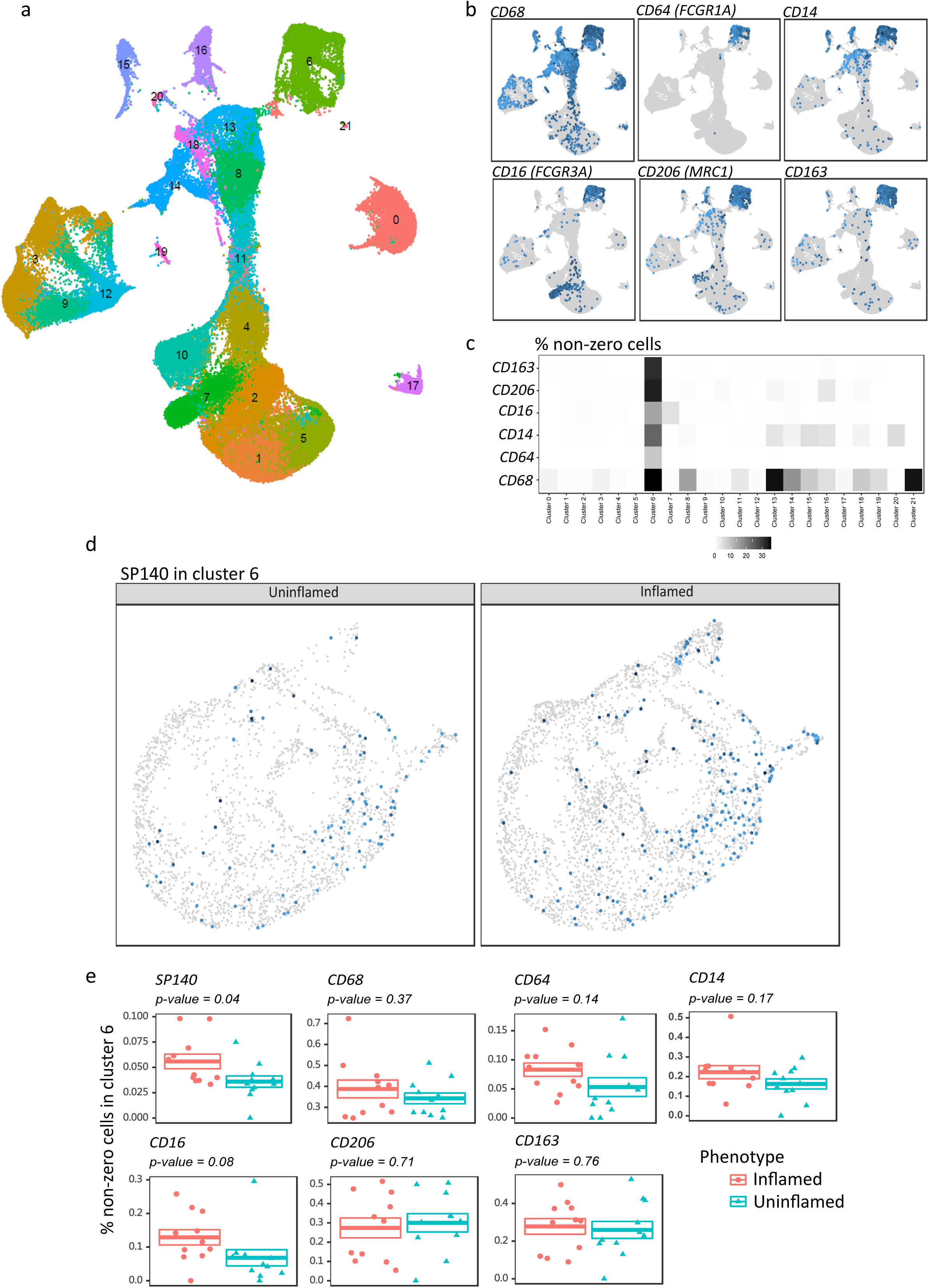
The percentage of *SP140* expressing macrophages in cluster 6 is higher in inflamed tissue compared with uninflamed CD-ileal biopsies. Publicly available single-cell RNA-sequencing was used to investigate *SP140* expression in ileal macrophages in inflamed (n=11) and uninflamed (n=11) biopsies of CD patients. **(a)** Clustering analyses of all cells were performed using the top 2000 most variable genes and the top 15 principal components yielding 22. **(b)** Visual illustration of the actual counts of cells expressing different monocytes/macrophages markers. Darker blue represents more reads per cell. **(d)** The percentage (represented by the hue) of cells that are non-zero for the indicated macrophage marker genes **(c)**. UMAP of cluster 6 only with blue dots representing *SP140* expression per cell. **(e)** Comparative analyses of the percent non-zero cells when comparing inflamed with uninflamed, where the percentage was calculated relative to cluster 6 only.

### SP140 mediates inflammatory M1 macrophage function

Given the high SP140 expression in CD mucosal macrophages, we investigated whether SP140 is associated with inflammatory activation. Human CD14^+^ monocytes or THP-1 cells were differentiated into macrophages and then polarized into M1 and M2 phenotypes using IFN-γ and IL-4 respectively or left without treatment (M0) (Fig. 3a). The polarization was verified by measuring gene expression of *CD64* and *CCL5* (markers of M1 macrophages) and *CD206* and *CCL22* (markers of M2 macrophages) (Fig. S3a,b). *SP140* mRNA was expressed at significantly higher levels in M1 compared to M0 or M2 macrophages derived from primary monocytes (Fig. 3b) or from THP-1 cells (Fig. S3c). Protein staining showed increased numbers of SP140 protein-containing speckles in the nucleus of M1 compared to M2 macrophages derived from both human monocytes (Fig. 3c) and THP-1 cells (Fig. S3d). LPS induced an increase in *SP140* gene expression in M0 and M1 macrophages, along with enhanced *CCL5* and decreased *CCL22* gene expression (Fig. S3e). The expression levels of 13 other BCPs *(SP140L, SP100, SP110, BRD2, BRD3, BRD4, BRD9, BRDT, BAZ2A, BAZ2B, PCAF, EP300 and CREBBP)* showed no increase in M1 compared to M0 macrophages (Fig. S4a), indicating that the association between *SP140* and inflammatory macrophages was not a common phenomenon amongst BCPs.

**Figure 3:**
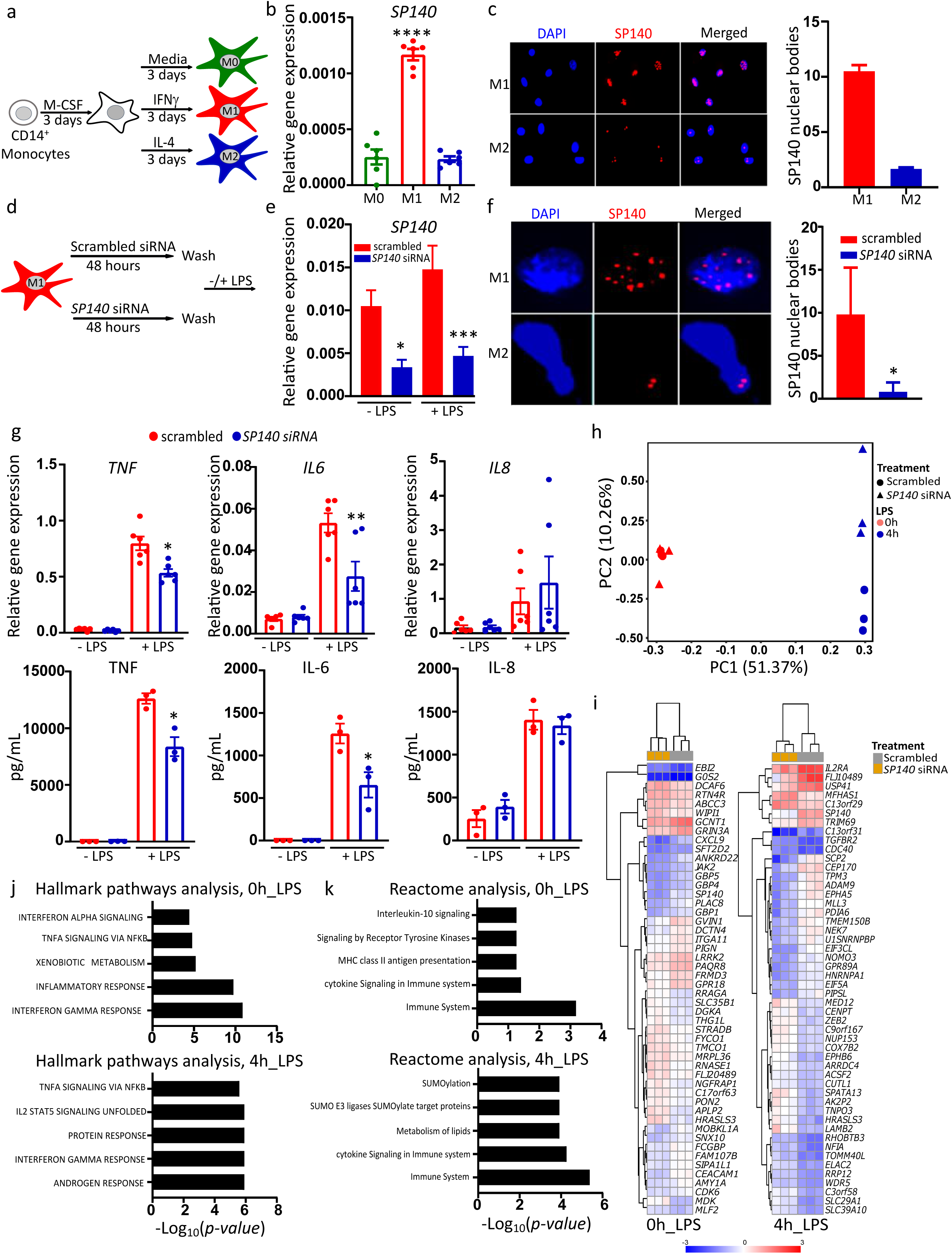
*SP140* knock-down reduces the activity of the inflammatory macrophages. **(a)** Polarization protocol of human CD14^+^ monocytes to M0, M1 and M2 macrophage phenotypes. Freshly isolated CD14^+^ monocytes were differentiated for 3 days using 20 ng/mL M-CSF. The cells were then washed with PBS and polarized for 3 days to M0, M1 and M2 macrophages using media only, 100 ng/mL IFN-γ or 40ng/mL IL-4, respectively. **(b)** Relative gene expression (qPCR) of *SP140* in M0, M1 and M2 macrophages, n=6. **(c)** Immunofluorescence staining of SP140 speckles in M1 and M2 macrophages imaged by microscopy (left) or nuclear bodies count (right). **(d)** *SP140* silencing protocol; M1 macrophages were incubated with either scrambled control siRNA or *SP140* siRNA for 48h. The cells were then washed with PBS and stimulated with 100 ng/mL LPS for 4h (for qPCR) and 24h (for Elisa) or kept without LPS-stimulation. (**e**) The efficiency of *SP140* silencing was assessed by measuring relative gene expression (qPCR) of *SP140* (n=6) and **(f)** immunofluorescence staining of SP140 speckles nuclear bodies (left) and nuclear bodies count (right). **(g)** Relative gene expression (qPCR) (top) n=6, and protein levels (ELISA) (bottom) n=3, of TNF, IL-6 and IL-8. **(h)** PCA of microarray dataset; PC1 represents most variance associated with the data (LPS-stimulation) and PC2 represents second most variance (siRNA), n=3. **(i)** Heatmap of top 50 DEGs at 0h LPS or 4h of 100 ng/mL LPS (non-annotated genes were not included). **(j)** Hallmark pathways analysis at 0h (top) and 4h of 100 ng/mL LPS (bottom). **(k)** Reactome pathway analysis carried out using ShinyGO v0.60 in unstimulated M1 macrophages (bottom) or 4h of 100 ng/mL LPS-stimulated M1 macrophages (bottom). In all assays, statistical significance is indicated as follows: \**P* < 0.05, \*\**P* < 0.01, \*\*\**P* < 0.001, \*\*\*\**P* < 0.0001.

To investigate SP140 function, we initially used siRNA-mediated knock-down to reduce *SP140* expression in M1 macrophages, achieving approximately 75% reduction in mRNA (Fig. 3e) and a clear decrease in the number of SP140 protein-containing speckles (Fig. 3f). Knockdown seemed specific to *SP140*, as expression of related family members *SP110* (Fig. S4b) and *SP100* was not affected (Fig. S4b,c). After knockdown, cells were simulated with LPS or kept unstimulated (Fig. 3d). *SP140* silencing led to a decrease in LPS-induced IL-6 and TNF mRNA and protein levels (Fig. 3g). After transcriptional profiling, principal component analysis (PCA) revealed that the variance was most associated with *SP140* siRNA treatment in LPS-stimulated M1 macrophages (Fig. 3h). Downregulation of some key CD-associated genes, notably *CXCL9, CEACAM1* and *JAK2*, was apparent amongst *SP140*-siRNA-treated unstimulated macrophages, and *IL2RA, TRIM69* and *ADAM9* amongst *SP140*-siRNA LPS-stimulated macrophages (Fig. 3i). *SP140* silencing affected common inflammatory pathways, such as TNF-signaling via NFKB, IFN-γ response, inflammatory response (Fig. 3j) and cytokine-signaling (Fig. 3k).

### The development of a selective inhibitor of SP140 (GSK761)

To identify selective SP140 binding compounds, we utilized encoded library technology to screen the GSK proprietary collection of DNA-linked small molecule libraries^29, 30^. Affinity selection utilizing a recombinant protein construct spanning the PHD and Brd domains of SP140 was carried out, leading to identification of an enriched building block combination in DNA Encoded Library 68 (DEL68) (Fig. 4a). Representative compounds of the identified three-cycle benzimidazole chemical series were synthesized, which yielded the small molecule, GSK761 (Fig. 4b). To quantify the interaction between GSK761 and SP140 (aa 687-867), the dissociation constant (*K*_d_) was determined using a fluorescence polarization (FP) binding approach. A fluorophore-conjugated version of GSK761, GSK064 (Fig. 4c), was prepared and subsequently used to determine a *K*_d_ value of 41.07 ± 1.42 nM for the interaction with SP140 (aa 687-867) (Fig. 4d). To further validate this interaction, competition studies were carried out using GSK761 in a FP-binding assay configured using GSK064 and SP140 (aa 687-867). Competitive displacement of GSK064 from SP140 (aa 687-867) by GSK761 was observed, subsequently leading to the determination of an IC_50_ value of 77.79 ± 8.27 nM (Fig. 4e). Prior to utilizing GSK761 for cell-based binding studies to endogenous SP140, the cell penetration capacity of the compound was assessed using mass spectrometry and the methodology described by ^28^. Concentration measurements showed that GSK761 had a pΔC_total_ value of 1.45 ± 0.21 (data not shown), indicating that GSK761 permeates human cells and accumulates intracellularly by an order of magnitude, when compared to GSK761 free in solution.

**Figure 4:**
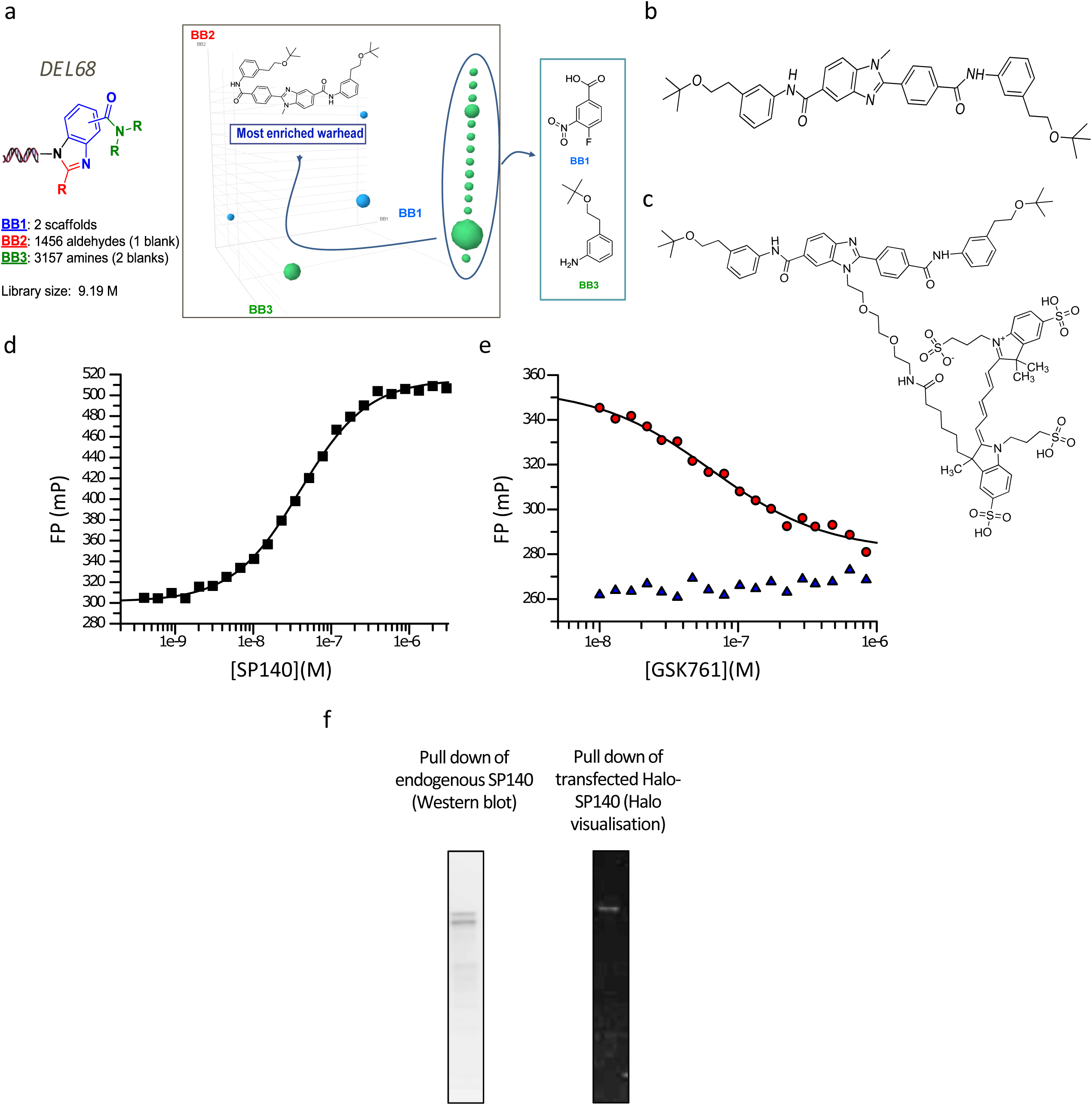
The Development of the first small molecule inhibitor of SP140 (GSK761) and investigating its affinity and selectivity. **(a)** Three-cycle benzimidazole library (left) and cube view of the SP140 selection output (right). Each individual dot in the cube represents a DEL68 molecule, while the size of the dot corresponds to the total copy number (2-24) identified for that library molecule. The dots are colored by the BB3s that compose the library molecules. Library members with a single copy were removed to simplify visualization revealing a prominent line defined by a specific BB1&BB3 combination. **(d)** Biochemical characterization of the interaction between **(b)** GSK761 and recombinant SP140 in a FP binding assay using **(c)** GSK064, generated by fluorescent labelling of GSK761. The mean binding affinity for this interaction was a *K*_d_ = 41.07 ± 1.42 nM (n = 5). **(e)** A FP binding assay was configured using recombinant SP140 and GSK064, which was used to determine the potency of GSK761.Displacement of GSK064 from SP140 by GSK761 (circles) was achieved and determined to have a mean IC_50_ of 77.79 ± 8.27 nM (n = 3). No effect on GSK064 motion was observed in the presence of varying concentrations of GSK761 (triangles). The data presented in **(d)** and **(e)** are representative data form a single experimental replicate, affinities and potencies are mean values determined from multiple test occasions. **(f)** An endogenous SP140 (HuT78 nuclear extracts) and Halo-transfected SP140 were pulled-down using biotinylated GSK761 and visualized by Western blotting and gel electrophoresis.

To confirm the binding of GSK761 to full-length SP140, immobilized GSK761 was used to probe for SP140 in nuclear extracts from HuT78 cells and HeLa cells transfected with Halo-tagged SP140. Both endogenous and Halo-tagged SP140 were pulled down with biotinylated GSK761 and visualized via Western Blotting (Fig. 4f). Endogenous SP140 is observed as a doublet containing the four largest isoforms (predicted 86-96KDa)^10^ while Halo-tagged SP140 exhibits a single band. The specificity of GSK761 for SP140 was profiled using the BROMOscan® Assay, in which DNA tagged BCPs were incubated with an increasing concentration of GSK761 or DMSO. Binding is assessed by measuring the amount of bromodomain captured in GSK761 vs DMSO samples by using an ultra-sensitive qPCR. There was evidence of low affinity interaction between GSK761 and several tested BCPs, with binding detected at concentrations >21000 nM. However, no binding was detected at concentrations ≤ 21000 nM indicating a high degree of specificity of GSK761 for SP140 (Supplemental table 2).

### GSK761 reduces the inflammatory activation of macrophages and expression of pro-inflammatory cytokines

SP140 expression was selectively increased in M1 compared to M0 macrophages, prompting us to test whether SP140 is required for the polarization to inflammatory macrophage phenotype. We first tested whether GSK761 demonstrated any toxicity in macrophages to determine the appropriate dose range. At ≤0.12 µM, GSK761 showed no cytotoxicity (Fig. S5a,b). To investigate the effect of GSK761 on macrophage polarization to inflammatory phenotype, M0 macrophages were treated with either DMSO or GSK761 for 3 days in presence of IFN-γ or IL-4 (during the polarization to M1 and M2 macrophages, respectively). GSK761 enhanced mRNA expression of the anti-inflammatory marker *CD206* in both M1 and M2 macrophages and decreased the pro-inflammatory marker *CD64* in M1 macrophages (Fig. 5a). FACS analysis showed that GSK761-treatment prior to IFN-γ (M1) polarization reduced CD64^+^ cells and CD64 protein expression (Fig. 5b) and increased CD206^+^ cells and CD206 protein expression (Fig. 5c). Adding GSK761 to the cells during IFN-γ (M1) polarization lowered *TNF* (M1 polarization marker) gene expression in these cells compared to the control (Fig. S5c). Altogether, these data suggest that SP140 inhibition during M1 polarization biased differentiation towards a regulatory M2 phenotype.

**Figure 5:**
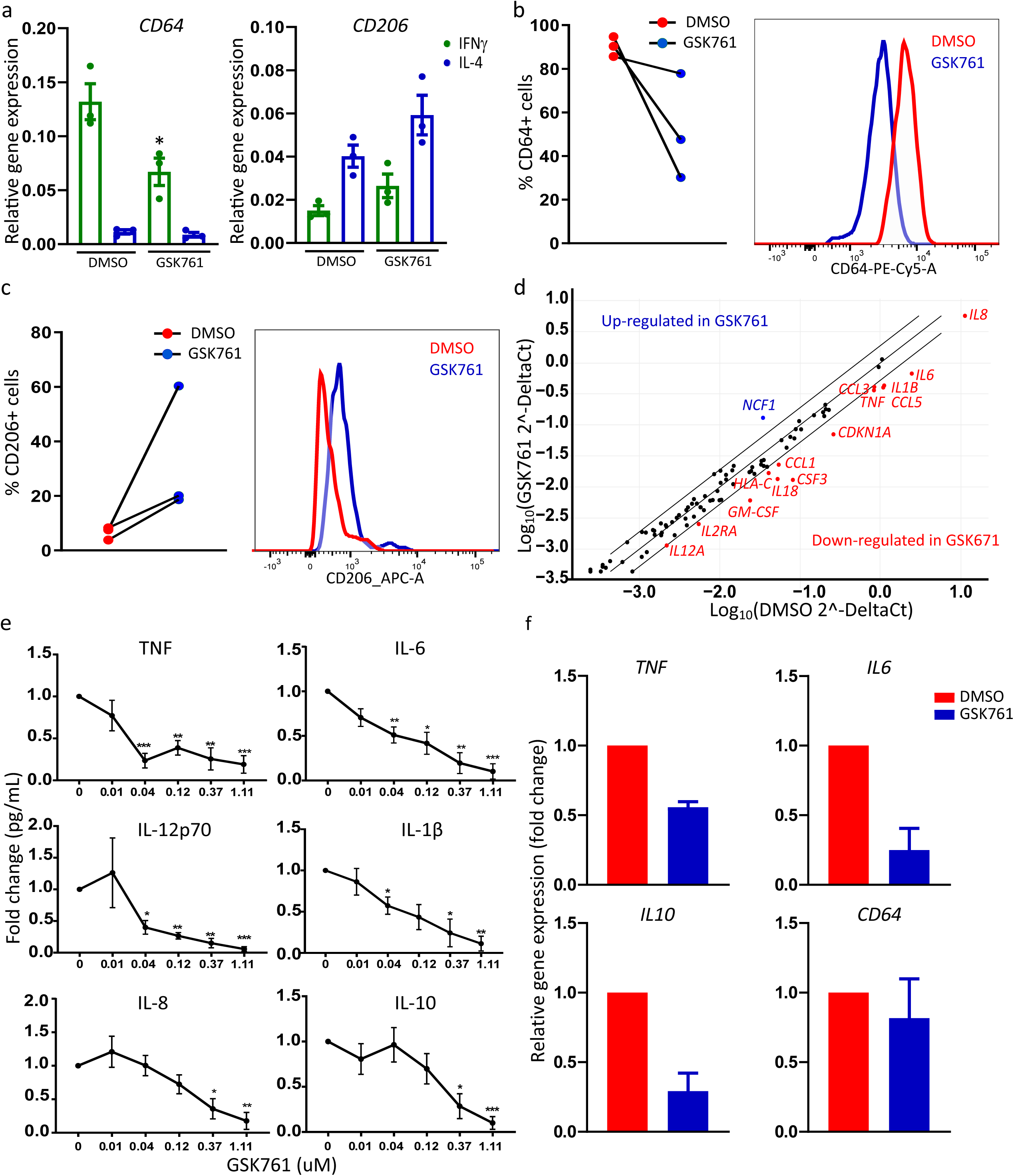
GSK761 biases differentiation towards an CD206 expressing macrophages and reduces pro-inflammatory cytokine expression in CD14^+^ macrophages isolated from CD anti-TNF refractory patients’ colonic-mucosa. Human primary CD14^+^ monocytes were differentiated with 20 ng/mL M-CSF for 3 days. The cells where then washed with PBS and treated with 0.1% DMSO or 0.04 µM GSK761 for 1h prior to 3 days polarization to M1 and M2 macrophages phenotypes with 100 ng/mL INF-γ or 40 ng/mL IL-4, respectively (GSK761 and DMSO were not washed and kept in the culture during the 3 days of polarization). **(a)** Gene expression of the inflammatory marker CD64 (M1 marker) and anti-inflammatory marker CD206 (M2 marker) was measured by qPCR in IFN-γ(M1) and IL-4(M2) polarized macrophages and **(b**,**c)** FACS analysis was performed in IFN-γ(M1) polarized macrophages to assess: **(b)** the frequency of CD64^+^ macrophages (left) and CD64 protein expression intensity (right) and **(c)** the frequency of CD206^+^ macrophages (left) and CD206 protein expression intensity, n=3. For FACS data, the count was normalized to mode. **(d)** M1 polarized macrophages were pretreated for 1h with 1% DMSO or with 0.04 µM GSK761. The cells were then stimulated for 4h with 100 ng/mL LPS and Customized RT^2^ Profiler PCR Arrays was performed, the scatter plot illustrates the differentially expressed genes (2-fold change), n=4 to 5. **(e)** M1 polarized macrophages were pretreated for 1h with 1% DMSO or with an increasing concentration of GSK761 (0.01, 0.04, 0.12, 0.37. 1.11 µM). The cells were then stimulated for 24h with 100 ng/mL LPS. Protein levels of TNF, IL-6, IL-1β, IL-10, IL-8 and IL-12p70 were measured in the supernatant using ProInflammatory 7-Plex MSD kit, n=4 to 7. The Y axis indicates the fold change in cytokine protein expression relative to DMSO control. **(f)** CD14^+^ cells were isolated from inflamed CD mucosa. The cells were then incubated *ex vivo* for 4h with either 0.1% DMSO or 0.04 µM GSK761. Relative gene expression of *TNF, IL6, IL10* and *CD64* were measured using qPCR, n=2. The Y axis indicates the fold change in mRNA level relative to DMSO control. Statistical significance is indicated as follows: \**P* < 0.05, \*\**P* < 0.01, \*\*\**P* < 0.001, \*\*\*\**P* < 0.0001.

We next assessed the effect of SP140 inhibition on the response of M1 polarized macrophages to inflammatory stimuli. LPS-stimulated M1 macrophages pretreated with GSK761 showed a strong reduction in secretion of IL-6, TNF, IL-1β and IL-12 (Fig. 5e) at concentrations where cytotoxicity was not observed (0.04 and 0.12 µM). When employing RNA transcriptional profiling using a customized qPCR array, SP140 inhibition was found to reduce the expression of many other pro-inflammatory cytokines and chemokines including *GM-CSF, CCL3, CCL5* and *CCL1* (Fig. 5d).

The marked anti-inflammatory effects of GSK761 on macrophages *in vitro* suggests that targeting SP140 may be an effective approach for IBD. Unfortunately, due to poor *in vivo* pharmokinetics (data not shown), GSK761 was not suitable to evaluate the effects of SP140 inhibition in *in vivo* animal models of colitis. To evaluate the impact of SP140 inhibition in human CD, CD14^+^ mucosal macrophages were isolated from CD anti-TNF refractory patients’ colonic-mucosa and then cultured *in vitro* with either DMSO or GSK761 for 4h. Spontaneous gene expression of *TNF, IL6* and *IL10* was strongly decreased in GSK761-treated macrophages, demonstrating that GSK761 inhibited their immune reactivity (Fig. 5f). Previous studies have demonstrated that successful anti-TNF therapy is associated with an increase in CD206^+^ regulatory macrophages^31^ and a decrease in pro-inflammatory cytokines^10^ and SP140 expression^10^ in CD mucosa. Thus, our data suggests that SP140 inhibition could be also beneficial in supporting the anti-TNF therapy.

### SP140 preferentially interacts with transcription start sites (TSS) and enhancer regions

SP140 possesses multiple chromatin binding domains, namely Brd, PHD and SAND (<u>S</u>P100, <u>A</u>IRE-1, <u>N</u>ucP41/75, <u>D</u>EAF-1) domains, which may allow it to function as an epigenetic reader ^10, 32^. Indeed, we found that recombinant SP140 Brd-PHD protein bound to histone H3 peptides with sub-µM affinity (Fig. S6a). The strongest binding was observed for an unmodified peptide corresponding to the N-terminal 21 aa, while methylation at lysine residue 4 (K4) led to substantially (∼30-fold) reduced affinity (Fig. S6b). K9 acetylation also resulted in lower affinity binding (4-5-fold), while SP140 binding was unaffected by either acetylation or methylation of K14 (Fig. S6c). Similar binding affinity was observed for unmodified H3_1-21_ vs H3_1-18_, while reduced (4-5-fold) but still significant SP140 binding was measured for H3_1-9_ (Fig. S6c). Taken together, these data suggest that the SP140 Brd-PHD module functions as a reader for un-modified histone H3, with binding focused around the N-terminal 9 aa.

To evaluate whether native SP140 is also capable of binding to histones, we used immobilized histone H3 peptides with varying modifications to precipitate proteins from nuclear extracts of anti-CD3/CD28-stimulated HuT78 T cells and probed via Western blotting for the presence of SP140. SP140 was efficiently pulled down by un-modified H3_1-21_ (Fig. 6a). Notably, H3_1-21_ peptides bearing certain methylation and acetylation marks (K4me3, K9me3, K9ac and K14ac) were also capable of capturing native SP140 (Fig. 6a), despite the lower measured affinity of some of these for recombinant SP140 Brd-PHD (see above). To evaluate whether SP140 interacts with histone H3 in a cellular setting, we utilized a NanoBRET system (Fig. S6d). NanoBRET is a proximity-based assay that can detect protein interactions by measuring energy transfer from a bioluminescent protein donor NanoLuc® (NL) to a fluorescent protein acceptor HaloTag® (HT). This energy transfer was observed in the nuclei of HEK293 cells transfected with SP140-NL and Histone3.3-HT DNA, indicating close proximity of the two proteins.

**Figure 6:**
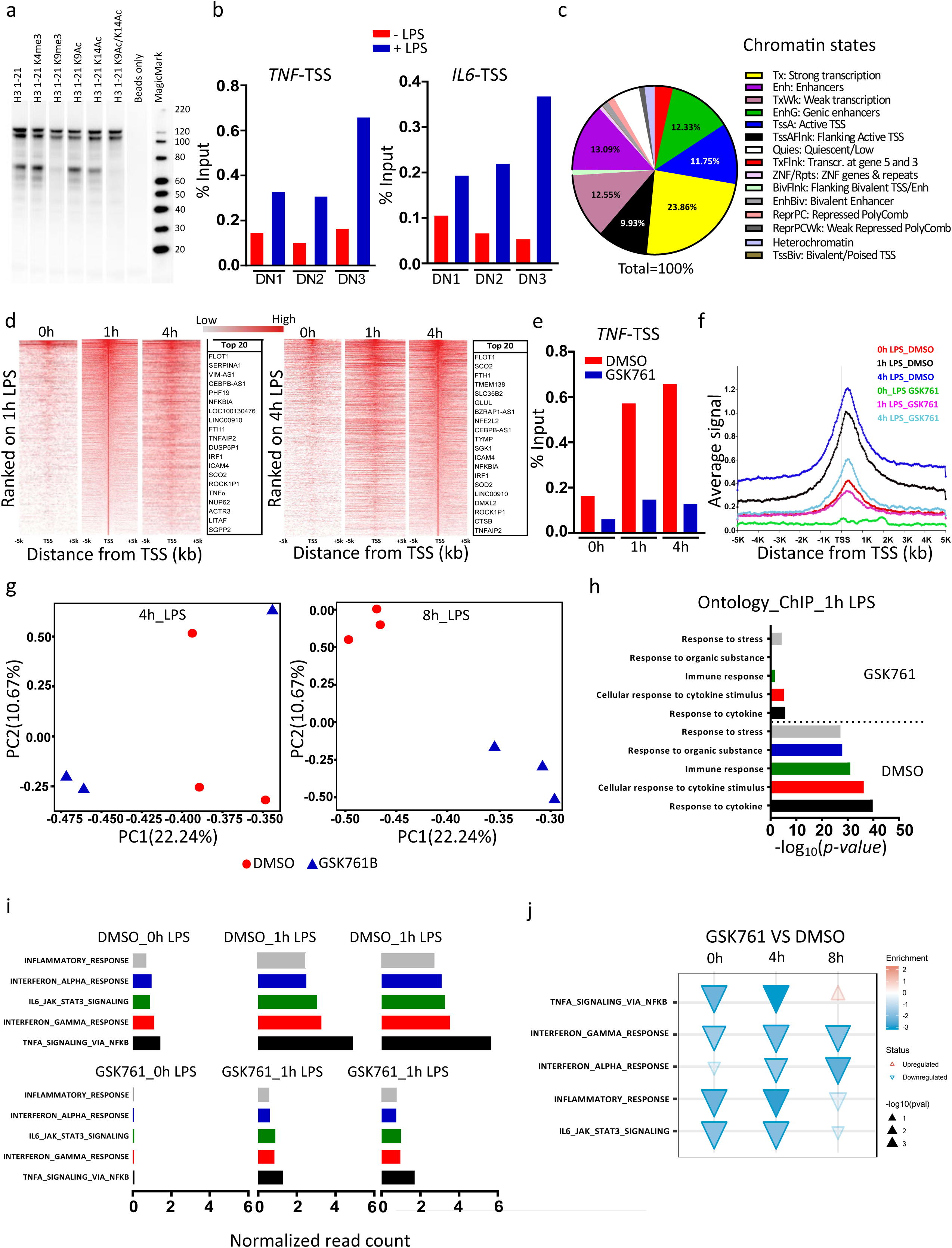
LPS-stimulation enhances SP140 protein recruitment to chromatin; GSK761 reduces this recruitment and dampens inflammatory pathways in inflammatory macrophages. **(a)** Nuclear extracts from αCD3/αCD28 stimulated HuT78 cells were incubated with unmodified or modified (acetylated and methylated) histone H3 peptides. SP140 was then pulled–down and its interaction with H3 peptides was visualized. **(b)** ChIP-qPCR of SP140 occupancy at TSS of *TNF* and *IL6* in M1 macrophages stimulated with 100 ng/mL LPS for 4h or without LPS-stimulation, n=3 donors (DN).**(c)** Epigenome roadmap scan illustrating proportions of SP140 genome-wide occupancy after SP140 ChIP-seq experiment. **(d)** Heatmap (1000 top SP140-bound genes) of SP140 ChIP-seq reads ranked on 1h LPS-stimulated (left) or on 4h LPS-stimulated (right) M1 macrophages rank-ordered from high to low occupancy centered on TSS. Top 20 genes with high SP140 occupancy were listed. **(e)** ChIP-qPCR of SP140 occupancy at TSS of *TNF* in M1 macrophages pretreated with 0.1% DMSO or 0.04 µM GSK761 for 1h and then stimulated with 100 ng/mL LPS for 1 or 4h or kept unstimulated (0h LPS). **(f)** Metagene created from normalized genome-wide average reads for SP140 centered on TSS. **(g)** PCA of RNA-seq comparing 0.1% DMSO to 0.04 µM GSK761 treated M1 macrophages after 4h of LPS-stimulation (left) or after 8h of LPS-stimulation (right). **(h)** SP140 ChIP-seq gene ontology analysis of the most enriched molecular function and biological process after 1h of LPS-stimulation, comparing 0.1% DMSO with 0.04 µM GSK761 treated M1 macrophages. **(i**,**j)** Hallmarks pathway enrichment analysis at **(i)** 0, 1 and 4h of 100 ng/mL LPS-stimulation for ChIP-seq and at **(j)** 0, 4 and 8h of 100 ng/mL LPS-stimulation for RNA-seq. **(j)** The direction and color of the arrow indicates the direction and size of the enrichment score, the size of the arrow is proportional to the –log_10_(*p*-value), and non-transparent arrows represent significantly affected pathways.

To assess whether SP140 associates with chromatin in macrophages and how this might be regulated in inflammation, we initially conducted ChIP-qPCR experiments in unstimulated or LPS-stimulated M1 macrophages. Binding of SP140 to the TSS of *TNF* and *IL6* genes was observed in unstimulated cells, and this was increased following LPS-stimulation (Fig. 6b). We then evaluated chromatin occupancy of SP140 on a genome-wide level using ChIP-seq and tested the effects of GSK761 on both SP140 binding and gene expression (RNA-seq) in the context of LPS-stimulation. In DMSO-treated, LPS-stimulated macrophages, epigenome roadmap scan analysis revealed that the majority of SP140 occupancy was at strong transcription associated-regions, enhancers and TSS regions (Fig. 6c). However, this occupancy was decreased when the cells were pretreated with GSK761 at 1h (Fig. S6f) and 4h (data not shown) post LPS-stimulation. Heatmap rank ordering of SP140 occupancy in unstimulated and LPS-stimulated M1 macrophages showed a strong SP140 enrichment at the TSS (Fig. 6d). The top 20 enriched genes belonged mostly to those involved in the innate immune response, including *TNF, ICAM4, IRF1, LITAF, TNFAIP2* and *NFKBIA* (Fig. 6d). However, the most SP140-enriched gene *FLOT1* (Fig. 6d) has been reported to be strongly involved in tumorigenesis^33^. We found SP140 also to occupy active enhancers, as marked by H3K27Ac in human macrophages by overlapping the publicly available H3K27Ac ChIP-seq data (GSE54972) with our SP140 ChIP-seq dataset at 1h of LPS-stimulation (Fig. S6e). ChIP-qPCR showed a strong reduction of SP140 occupancy at the *TNF*-TSS in GSK761-treated M1 macrophages, which was reduced to that of unstimulated cells (Fig. 6e), correlating with the previously observed reduced expression of TNF in GSK761-treated macrophages.

Following LPS-stimulation, we observed a marked increase in binding at the TSS of the top 1000 genes occupied by SP140, reaching its maximum at 4h (Fig. 6f). GSK761-treatment prior to LPS-stimulation strongly reduced SP140 occupancy at the TSS (Fig. 6f) associated with an altered LPS-induced gene expression (Fig. 6g). PCA of RNA-seq data revealed a large separation in global gene expression between DMSO- and GSK761-treated macrophages at both time points post of LPS-stimulation (4h and 8h) (Fig. 6g).

Gene Ontology and Reactome enrichment analysis of SP140-bound genes showed enrichment of genes which participate in the processes of cytokine response and immune response that were inhibited through GSK761 pre-exposure (Fig. 6h and S6g). To elucidate the regulatory pathways enriched by SP140 and affected by GSK761, we carried out hallmark pathway analysis for differentially SP140 bound-genes (DBGs) (Fig. 6i) and DEGs (Fig. 6j) upon DMSO- or GSK761-treatment at each time point of LPS-stimulation. In DMSO-treated M1 macrophages, we found that SP140 binding is predominantly enriched at many pathways typically defined as inflammatory such as TNF-signaling via NFKB, and inflammatory response (Fig. 6i). However, this enrichment was no longer seen in GSK761-treated macrophages (Fig. 6i). Similar pathways were observed for DEGs and were downregulated in GSK761-treated macrophages, especially at 4h (Fig. 6j). In addition, we observed an up-regulation of MYC targets and oxidative phosphorylation gene sets in GSK761-treated macrophages, whereas these normally are down-regulated because of the induction of aerobic glycolysis upon LPS-stimulation of macrophages (Fig. S7a)^34^. These data suggest SP140 as a critical regulator of genes involved in the inflammatory response and that GSK761 can inhibit this response through displacing SP140 binding to TSS and enhancer regions.

### SP140 preferentially controls the expression of specific gene sets involved in the innate immune response

We next investigated the functional significance of inhibiting SP140 binding for gene expression. Heatmap (Top 100) and volcano plots of DEGs after 4h LPS-stimulation demonstrated a strong effect of GSK761 on gene sets that are involved in innate immune response such as the downregulation of *TNFSF9, IL6, F3, CXCL1, CCL5, IL1β, TRAF1, IL23A, IL18* and *GEM* and the upregulation of *PLAU* and *CXCR4* (Fig. 7a,b). We next explored the global DBGs in 1h LPS-stimulated M1 macrophages using R2 TSS-plot (Fig. 7c) and R2 TSS-peak calling (Fig. 7d). Interestingly, the top DBGs belonged to TNF-signaling via NFKB such *as TNFAIP2, TNF, LTA, TRAF1, IRF1* and *NFKBIA* (Fig. 7c,d). GSK761 clearly reduced SP140 binding to those genes (Fig.7,d). Similarly, gene expression of *TNFAIP2, TAF1, TNF, LTA*, and *IRF1* was reduced (Fig. 7e). By conducting a focused TNF-signaling enrichment analysis (Fig. S7b) and R2 TSS-plot analysis (Fig. S7c), we illustrated a new set of DBGs including TFs (*TNFAIP3, CEBPB, ATF4, MAP3K8, MAP2K3* and *JUNB*), cytokines (*IL1β, IL23A*) and adhesion molecules (*ICAM1*). Most of these genes were DEGs (Fig. S7d).

**Figure 7:**
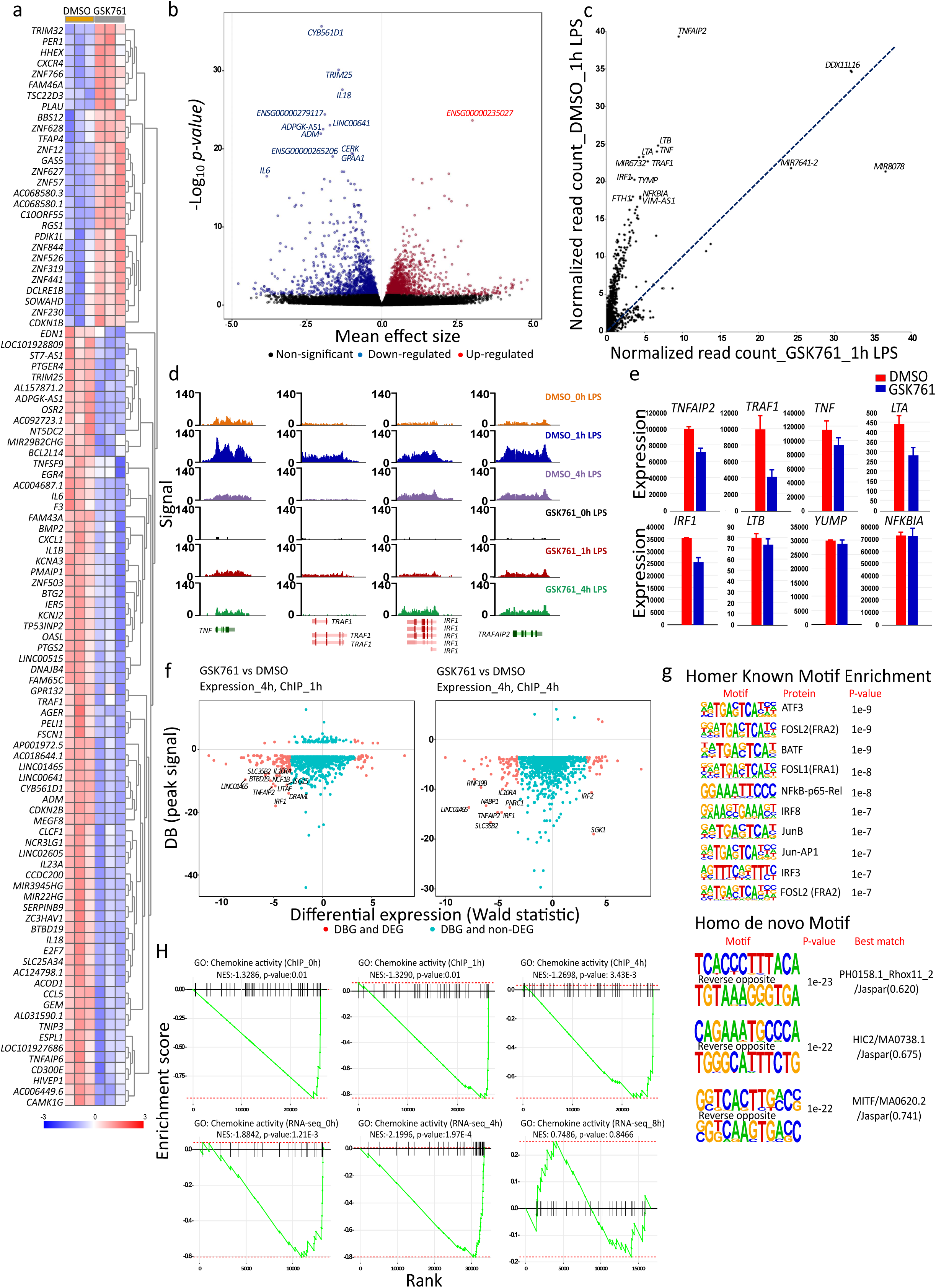
SP140 preferentially controls the expression of specific gene sets involved in the innate immune response. **(a)** Heatmap of the top 100 DEGs and **(b)** volcano plots of the all genes comparing 0.1% DMSO-with 0.04 µM GSK761-treated M1 macrophages after 4h of 100 ng/mL LPS-stimulation. **(c)** R2 TSS-plot comparing a global DBGs after 1h of 100 ng/mL LPS-stimulation of 0.1% DMSO- and 0.04 µM GSK761-treated M1 macrophages. **(d)** SP140 ChIP-seq genome browser view of some of the most affected DBGs; *TNF, TRAF1, IRF1* and *TRAFAIP2*. Y axis represents signal score of recovered sequences in 0.1% DMSO and 0.04 µM GSK761 treated macrophages after 0, 1 and 4h of 100 ng/mL LPS-stimulation. **(e)** RNA-sequencing derived gene expression of some of the most DBGs. (**f)** Comparative analyses of the top 1000 DBGs (signal) with their gene expression (Wald statistic). Gene expression at 4h and ChIP at 1h (left) and gene expression at 4h and ChIP at 4h (right), Y axis represents the SP140 differential binding signal (BD). **(g)** Homer Known Motif Enrichment using TF motifs and their respective p-value scoring (top) and Homer de *novo* Motif results with best match TFs (bottom) in 0.1% DMSO treated M1 macrophages after 1h of 100 ng/mL LPS-stimulation. **(h)** An enrichment analysis targeted chemokine activity for SP140 ChIP-seq (top) and RNA-seq (bottom) comparing 0.1% DMSO-with 0.04 µM GSK761 treated-M1 macrophages at 0, 1 and 4h of 100 ng/mL LPS-stimulation.

We next verified the impact of global reduced SP140 binding on gene expression by integrating the DBGs and the DEGs. Interrogating the top 1000 DBGs (at 1h or 4h LPS) for their gene expression (at 4h or 8h LPS), implicated genes that are involved in TNF-signaling pathway (*NFKBIA, SOD2, LITAF, SGK1, PNRC1, IRF1* and *RNF19B*) and cytokine-mediated signaling pathway (For example: *IL10RA, NFKBIA, IRF1, SOD2, CAMK2D, IRF2* and *ISG15*) which showed the strongest concordant differential binding and expression (Fig. 7f and S8).

In addition to SP140 binding to chromatin, we speculated that SP140 may also interact with other TFs proteins to regulate gene expression. To this end, we performed homer known motif enrichment analysis (HKMEA) at DMSO_1h LPS (Fig. 7g). We also conducted a homo de novo motif discovery to define new DNA sequence motifs that are recognized specifically by SP140 (Fig. 7g). Interestingly, HKMEA suggests that SP140 may bind the same DNA sequence motifs recognized by certain TFs defined as key proteins in TNF-signaling via NFKB (ATF3, FOSL2, FOSL1, NFKB-p65-rel, JUNB and JUN-AP1) and of cytokine-mediated signaling pathway (BATF, NFKB-p65-rel, JUNB, IRF8 and IRF3) (Fig. 7g).

LPS-stimulation strongly recruits SP140 to multiple human leukocyte antigen (HLA) genes, including *HLA-A, HLA-B, HLA-C, HLA-F, HLA-DPA* and *HLA-DPB* (Fig. S7e), binding which was strongly reduced by GSK761. This indicates a role of SP140 in regulating antigen presentation-associated genes. Notably, the Brd-containing protein 2 (*BRD2*) gene was occupied by SP140, and GSK761 reduced this binding and lowered BRD2 gene expression (Fig. S7e and table 5). BRD2 has been reported to play a key role in inflammatory response in murine macrophages and in inducing insulin resistance^35^.

Finally, we noticed that SP140 occupancy on TSS of chemokine activity genes was dramatically reduced by GSK761, inducing a reduced global chemokine activity at mRNA level (Fig. 7h). R2 TSS-plots and R2 TSS-peak calling indicate a strong SP140 occupancy at several inflammatory macrophage-associated chemokines (*CCL2, CCL3, CCL4, CCL5, CCL8, CXCL1, CXCL3, CXCL8 CXCL9, CXCL10 and CXCL11)*, chemokine ligands *(CCL4L1, CCL3L1, CCL3L3, CCL4L2*), chemokine receptors (*CCR7*), TFs (*STAT1, JAK2, PIK3R5, RELA*) and protein kinases (*PRKCD, FGR and HCR*) (Fig. S9a,b,c). This binding was reduced by GSK761, affecting expression of many genes involved in chemokine-signaling (Fig. S9d). However, SP140 binding was selective as it did not bind to the TSS of a range of other chemokines such as *CXCL6, CXCL5, CCL11*, and *CCL7* (Fig. S9c).

## Discussion

SP140 has been implicated in CD and other autoimmune diseases through genetic and epigenetic association studies^16,36^. Here, we describe the first small molecule inhibitor of SP140 protein, which has been used to investigate its function. The novel SP140-binding small molecule GSK761 was shown to compete with the N-terminal tail of histone H3 for interactions with the SP140 Brd-PHD module. In human macrophages, SP140 associated with the regulatory regions of immune-related/inflammatory genes in a stimulus-dependent manner. GSK761 inhibited SP140 binding to the TSS of many inflammatory genes, correlating with compound-induced decreased expression of these genes and reduced macrophage inflammatory function. The predominant SP140 expression in immune cells and in particular inflammatory macrophages, further identify SP140 as an epigenetic reader protein that contributes to inflammatory gene expression.

Inflammatory macrophages generated *in vitro* or in CD patients were shown to express high levels of SP140, and *SP140* silencing in macrophages reduced expression of a range of inflammatory genes. These anti-inflammatory effects were extended using GSK761, which was able to inhibit spontaneous cytokine expression from CD14^+^ macrophages isolated from CD anti-TNF refractory patients’ colonic mucosa. LPS-stimulation of macrophages was shown to cause SP140 recruitment to the TSS of a specific set of inflammatory genes, which was prevented by GSK761, dampening the induced expression of those genes. The strongest SP140 occupancy was at the TSS of a range of genes involved in TNF-signaling. TNF-signaling plays an integral role in CD as evidenced by the efficacy of chimeric anti-TNF mAbs-therapy^37^. Notably, anti-TNF therapy response rests on TNF neutralisation, but specifically in CD also on its potency to bind membrane bound TNF on CD-tissue infiltrated macrophages^38^, which causes differentiation of macrophages to a more regulatory CD206^+^ M2-like phenotype^31,39^. This is for example evidenced by the clinical observation using etanercept (does not bind membrane bound TNF), which is not effective in CD, whilst infliximab is^39^. A reduced expression of SP140 in CD-biopsies was shown to be correlated with a better anti-TNF response^10^. In this study, we found that GSK761 increased CD206 expressing macrophages in *in vitro*. Furthermore, GSK761 reduced pro-inflammatory cytokine expression in CD-mucosal macrophages of anti-TNF resistant patients. Thus, we anticipate that SP140 inhibition would be useful in enhancing anti-TNF remission-induction in CD patients. In this respect, SP140 was also shown by ChIP-seq analysis to bind and regulate expression of some key CD-associated genes such as *IL23A*^40^, *TYK2*^41^, *JAK2*^41^, *NOD2*^42^ and *CARD9*^43^.

In IBD, epigenetic mechanisms control macrophage inflammatory activation and polarization^44^. These macrophages are central components of the inflamed mucosa and contribute to disease pathology by producing inflammatory cytokines, which promote the differentiation and activation of Th1 and Th17 cells^45^. In this study, SP140 is shown to be critical for both the polarization of inflammatory macrophages and in the response of polarised macrophages to inflammatory stimuli. In particular, the TFs STAT1 and STAT2 are crucial for inflammatory macrophages polarization^5,46,47^. We found high levels of SP140 binding associated with the *STAT1 and STAT2* genes; GSK761 disrupted SP140 binding to these genes and reduced their expression, providing a probable mechanistic basis for the inhibitory effects of the compound on M1 polarisation. Notably, GSK761 inhibited macrophage-induced cytokines and co-stimulatory molecules required for inflammatory T cell activation including *IL23A*^40^, *IL12*^48^, *IL18*^49^, *IL1β*^49^, *CD40*^50^ and *CD80*^51^. Therefore, in addition to reducing the direct inflammatory function of macrophages, inhibition of SP140 may inhibit downstream activation/polarisation of responding T cells.

While the mechanism by which SP140 is targeted to gene regulatory regions remains to be determined, motif analysis indicated a strong overlap between the DNA sequences enriched for SP140 binding and the binding motifs of some key TFs involved in regulating the inflammatory response as well as the differentiation and polarization to inflammatory macrophages, such as NFkB-p50^52^, IRF3^53^ and IRF8^54^. Thus, it is possible that SP140 is recruited to chromatin via binding to stimulation-induced TFs – either before or after the TFs bind to DNA. Conversely, SP140 may associate with other proteins recruited to sites bound by these TFs. By conducting ChIP-seq at multiple time points (1 and 4h), we observed differential binding kinetics of SP140 to different sets of genes; for example, SP140 is bound to the *CCL5* TSS by 1h after LPS-stimulation but the TSS of *CCL2* only after 4h. The factors controlling the differential binding kinetics are currently unknown but could include differences in the initial chromatin environments of these genes in addition to distinct kinetics of TF activation and binding.

In a previous study, SP140 was found to be enriched at the *HOXA* genes in macrophages, and SP140 binding at these locations was proposed to play a role in repression of these lineage-inappropriate genes^10^. However, SP140 binding to this region was minimal in our study in both unstimulated and LPS-stimulated macrophages. The reason for this discrepancy is unclear, but the use of different SP140 antibodies in the two studies is one possibility. The antibody used in our ChIP experiments was found to be the most SP140-specific amongst 5 antibodies that we tested. Importantly, the displacement of SP140 binding by GSK761 provides strong support for the specificity of the SP140 binding identified in our ChIP-seq study.

Analysis of the gene sets identified in our SP140 ChIP-seq study suggested a role for SP140 in immune defence against microbes (in accordance with^55^ and^56^) and some other diseases, such as graft-versus-host-disease, cancers and rheumatoid arthritis. The intestinal microbiota is thought to play a central role in the pathogenesis of CD, and subsequent investigations could consider the role of SP140 in this respect as a previous study linked *SP140* SNPs to microbial dysbiosis in human intestinal microbiota^57^. In this study, ChIP-seq analysis was mainly focused on investigating SP140 binding to coding genes. However, by including non-coding RNA molecules, we observed very strong SP140 binding to a limited set of microRNAs (miRNAs) mainly involved in tumorigenesis, such as MIR3687^58^, MIR3648^59^ and MIR663A^60^. Accordingly, previous studies have demonstrated that certain BCPs (such as BRD4) are capable of binding and modulating transcription of some miRNAs involved in tumorogenesis^61,62^.

Contrary to the coding genes, GSK761 was unable to affect SP140 binding to miRNAs. This suggests that SP140 binding in these locations is independent of the Brd-PHD region with which GSK761 interacts; binding of SP140 to DNA via the SAND domain is one possibility. While not the focus of this study, other potential links between SP140 and cancer were observed, including high expression of SP140 in chronic lymphocytic leukemia cells (although reduced expression in acute myelogenous leukemia), and high binding of SP140 to the *FLOT1* gene, which has been implicated in tumorigenesis^33^

We conclude that SP140 is an epigenetic reader protein that regulates gene expression in macrophages in response to inflammatory stimuli, including those present in the CD intestinal mucosa of patients failing therapy. Targeting SP140 with a selective inhibitor was shown to displace SP140 from chromatin and decrease inflammatory gene expression in macrophages. This study suggests that SP140 inhibition could represent a promising approach for CD, which might include in addition to remission induction, the potential for combination with anti-TNF therapy.

## Competing interests

PDC, AYFLY, AS, SH, RPB, KG, GY, HPK, AC, AP, LAS, DJM, LL, RJW, CEB, LAH, GB, UK, NH, JRP, RKP, NRH and DFT were employed by GSK at the time of conducting this study. MG, JK, ILH, DAZ, OW, TBMH, JVL, PH, MPJW and WJDJ were employed by Amsterdam University Medical Centers at the time of conducting this study. GSK has a patent EP2643462B1 related to the therapeutic use of SP140 inhibitors.

## Acknowledgment

We thank Charlotte J. Barrett, previous student trainee at GSK for the assistance with a compound synthetic work.

## Funding

This project was funded by European Union’s Horizon 2020 research and innovation program under Grant Agreement No. ITN-2014-EID-641665. Further grant support is acknowledged from Dutch Ministry of Economical Affairs, LSH TKI, grants nr TKI-LSH T2017, and European Crohn’s and Colitis Organization (ECCO) Pioneer Grant, 2018.

## Author Contributions

Conduction of the study, laboratory and writing of the manuscript: MG; study design: MG, WJdJ, NRH and DFT; bioinformatics analyses; MG, JK, AYFLY and DAZ; supervision: WJdJ, NRH, DFT, MPJV and PH; reviewing and editing: WJdJ, NRH, DFT, MPJV, JVL, TBMH, PH and RKP; compound development: PDC, AS, SH, RPB, KG, LAS, DJM, GY, HPO, AC, AP, LL, RJW, CEB, LAH and UK; laboratory analysis: OW, NH, JRS and ILH; technical support: GB. All authors have read and agreed to the published version of the manuscript.

## Materials and methods

The human biological samples were sourced ethically, and their research use was in accord with the terms of the informed consents under an IRB/EC approved protocol. Written informed consent was obtained from each donor, as approved by the UK East of England - Cambridgeshire and Hertfordshire Research Ethics Committee.

### In-house GSK microarray profiler

Human gene expression data, along with accompanying sample descriptions, was purchased from GeneLogic (GeneLogic Division, Ocimum Biosolutions, Inc.) in 2006, and later organized by sample type. Gene expression data for each sample had been determined using mRNA amplification protocols as recommended by Affymetrix (Affymetrix, Inc.) and subsequently hybridized to the Affymetrix U133_plus2 chip. Purchased data was subject to reported quality control measures including ratios for *ACTB* and *GAPDH*, as well as maximal scale factors as reported by Affymetrix MAS 5.0. Expression data was normalized using MAS5.0 with a target intensity of 150.

### Immunohistochemistry

An anti-SP140 antibody produced in rabbit (HPA006162; Sigma Prestige Antibody) was used to visualize SP140 protein in a Cambridge Bioscience 69571061 Normal Human tissue microarray, a Cambridge Bioscience 4013301 Human Autoimmune Array, a Cambridge Bioscience 4013101 Human Colitis Array and on in-house rheumatoid arthritis synovial samples. Anti-SP140 antibody was detected with a Leica polymer secondary antibody. Sections were de-waxed using proprietary ER1, low pH 6 and ER2, high pH 8 buffers for 20 minutes at 98°C. Sites of antibody binding were visualized with peroxidase and DAB. Staining was performed on a Leica Bondman immunostaining instrument, using protocol IHC F.

### Tissue immunofluorescence of SP140 and CD68

Paraffin embedded tissue sections from inflamed and non-inflamed colon were obtained from patients with CD during surgical procedures, were first deparaffinized and washed with TBS. For antigen retrieval, slides were treated at 96°C for 10 minutes in 0.01 M sodium citrate buffer pH 6.0. The tissue sections were blocked and permeabilized with PBT (PBS, 0.1% Triton X-100 (Biorad), 1% w/v BSA (Sigma-Aldrich)) and incubated with primary antibodies for 2 hours (h) at room temperature (RT), to detect pan-macrophage marker CD68 (dilution 1:200) (M0876, Dako) and SP140 (dilution 1:200) (ab171141; Abcam). After washing, slides were stained with the following secondary antibodies: polyclonal goat anti-Mouse Alexa Fluor488 to detect CD68 protein (Green color) (A-11029, Invitrogen) or polyclonal goat anti-Rabbit Alexa Fluor546 to detect SP140 protein (Orange color) (A-11035, Invitrogen). Slides were mounted in SlowfadeGold reagent containing DAPI (4′,6-diamidino-2-phenylindole, Thermo-Fisher) and imaged using a Leica DM6000B microscope equipped with LAS-X software (Leica Microsystems).

### SP140 expression in ileal macrophages

Publicly available single-cell RNA-sequencing data from 11 involved (inflamed) and 11 paired uninvolved (uninflamed) ileal biopsies was downloaded from the sequence read archive accession number SRP216273^23^. Raw reads were aligned against GRCh38 using Cellranger (v3.1.0). The resulting unique molecular identifier (UMI) count matrices were then imported into the R statistical environment (v3.6.3) whereupon the samples were analyzed in an integrative fashion using Seurat (v3.1.5)^24,25^. Clustering analyses were performed using the top 2000 most variable genes and the top 15 principal components yielding 22 clusters whereupon they were visualized through UMAP (arXiv:1802.03426). We identified the monocytes and macrophages containing cluster through the expression of *CD68, FCER1A (CD64), CD163, MRC1 (CD206), CD14* and *FCGR3A (CD16A)*, after which *SP140* was visualized in the cluster representing monocytes/macrophages. Comparative cell count analyses were performed using the Wald test as implemented in DESeq2 (v1.24.0). Comparative percentage non-zero cells were performed using t-tests.

### Isolation, differentiation and polarization of primary human monocytes and THP-1 cells

Peripheral blood mononuclear cells (PBMCs) were obtained from whole blood of healthy donors (from Sanquin Institute Amsterdam or from GSK Stevenage Blood Donation Unit) by Ficoll density gradient (Invitrogen). CD14^+^ monocytes were positively selected from PBMCs using CD14 Microbeads according to the manufacturer’s instructions (Miltenyi Biotec). CD14^+^ cells were differentiated with 20 ng/mL of macrophage colony-stimulating factor (M-CSF) (R&D systems) for 3 days followed by 3 days of polarization into naïve macrophages (M0) (media only)^26^, classically activated (inflammatory) M1 macrophages (100 ng/mL IFN-γ; R&D systems)^26^ or alternative activated (regulatory) M2 macrophages (40 ng/mL IL4;R&D systems)^26^. Human monocytic cell line (THP-1) cells were differentiated with 100 nM phorbol myristate acetate (Sigma-Aldrich) for 3 days and then the same polarization protocol was performed. Cells were incubated in Isocove’s Modified Dulbecco’s Medium (Lonza) supplemented with 10 % fetal bovine serum (FBS) (Lonza), 2 mM l-glutamine (Lonza), 100 U/mL penicillin (Lonza) and 100 U/mL streptomycin (Lonza), at 37 °C, 5 % CO2.

### siRNA-mediated SP140 knockdown

Human M1 macrophages were generated *in vitro* from human primary CD14^+^ monocytes as described above. M1 macrophages were transfected with siGENOME human smartpool *SP140* siRNA or non-targeting scrambled siRNA for 48h with DharmaFECT™ transfection reagents according to manufacturer’s protocol (Dharmacon). The cells were left unstimulated or stimulated with 100 ng/mL LPS (E. coli 0111:B4; Sigma) for 4h (for qPCR) or 24h (for Elisa). The supernatant was harvested for cytokine measurement and the cells were lysed (ISOLATE II RNA Lysis Buffer RLY-Bioline) for RNA extraction.

### RNA isolation and Reverse Transcriptase, Polymerase Chain Reaction (PCR) and Real-Time Quantitative PCR

Total RNA was extracted from macrophages using RNeasy Mini Kit (Qiagen) following the manufacturer’s instructions. The concentration of RNA was determined using spectrophotometry (Nano-Drop ND-1000). Complementary DNA (cDNA) was synthesized with qScript cDNA SuperMix (Quanta Biosciences). PCR amplification of *SP140, SP100, SP110, SP140L, BRDT, BRD2, BRD3, BRD4, BRD9, EP300, BAZ2A, BAZ2B, PCAF, CREBBP, TNF, IL6, IL10, IL8, CCL5, CCL22, CD206* and *CD64* was performed by Fast Start DNA Master^plus^ SYBR Green I kit on the Light Cycler 480 (Roche, Applied Science). Relative RNA expression levels were normalized to the geometric mean of two reference genes *RPL37A* and *ACTB*. Primer sequences are listed in Supplementary table 1.

### Immunofluorescence cell staining

Primary human monocyte derived macrophages were polarized to M1 phenotype on coverslips. The cells were fixed with 4% paraformaldehyde and then permeabilized in 0.2% Triton (Biorad)/PBS. Blocking buffer 2% BSA (Sigma-Aldrich) was added for 30 minutes. Rabbit polyclonal anti-SP140 antibody (dilution 1:200) (ab171141; Abcam) or mouse polyclonal anti-SP100 antibody (dilution 1:200) (ab167605; Abcam) were added for 2h) followed by 2h of secondary antibody, Polyclonal goat anti-Rabbit, Alexa Fluor546 (A-11035, Invitrogen) (dilution of 1:1000) or goat anti-mouse, Alexa Fluor546 (A-21123, Life Technologies) (dilution of 1:500), respectively. DAPI (Thermo-Fisher) was used for nuclear detection.

### Microarray

RNA from human M1 macrophages transfected with *SP140* siRNA or scrambled siRNA was extracted using a RNeasy mini-kit (Qiagen). 150 ng total RNA was labelled using the cRNA labelling kit for Illumina BeadArrays (Ambion) and hybridized with Ref8v3 BeadArrays (Illumina). Arrays were scanned on a BeadArray 500GX scanner and data were normalized using quantile normalization with background subtraction (GenomeStudio software; Illumina). Genes with negative values were removed from analysis. Differentially expressed genes (DEGs) had a P-value <0.05 (analysis of variance). The data was analyzed in R (v3.6.3) and R2 Genomics Analysis and Visualization Platform-UMC (r2.amc.nl). Gene ontology overrepresentation analyses were performed in ShinyGO v0.60^27^.

### Discovery and synthesis of the compounds (GSK761, GSK064, GSK675 and GSK306)

The text describing the discovery and the synthesis of different compounds is added as supplementary information.

### Fluorescence Polarization (FP) Binding Affinity Studies

6HisFLAGTEVSP140 (687-867) was serially diluted in the presence of 3 nM GSK064 in Assay Buffer (50 mM HEPES, pH 7.5, 150mM NaCl, 0.05 % Pluronic F-127, 1 mM DTT) in a total assay volume of 10 µl. Following incubation for 60 minutes, FP was measured on a Perkin Elmer Envision multi-mode plate reader, by exciting the Alexa647 fluorophore of GSK064 at a wavelength of 620 nm and then measuring emission at 688 nm in both parallel and perpendicular planes. The FP measurement, expressed as milliP (mP), was then calculated using the following equation;

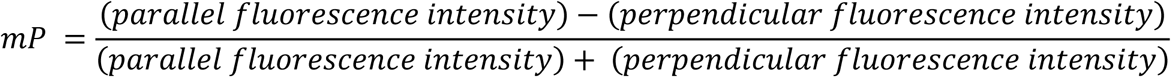

These data were then used to calculate an apparent *K*_d_ value by application of the following Langmuir-Hill equation; *AB* = (*A*_0_ + *B*)/(*K*_d_ + *B*) where AB is the concentration of the bound complex, A_0_ is the total amount of one binding molecule added, B is the free concentration of the second binding molecule and *K*_*d*_ is the dissociation complex. IC_50_ determination, GSK761 was serially diluted in DMSO (1% final assay concentration) and tested in the presence of 50 nM SP140 and 3 nM GSK064 in the same assay buffer and volume as above. Reactions were incubated for 60 minutes, and IC_50_ values calculated using a four-parameter logistic equation; 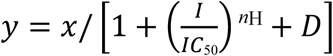 where *y* is the mP signal in the presence of the inhibitor at concentration I, *x* is the mP signal without the inhibitor, IC_50_ is the concentration of the inhibitor that gives 50% inhibition, *nH* is the Hill coefficient and *D* is the assay background.

### Pull-down of endogenous SP140 and Halo-transfected SP140 with GSK761

HuT78 cells were stimulated with 100 ng/mL anti-CD3 antibody (in-house reagent) and 3 mg/mL anti-CD28 antibody (in-house reagent) for 96h. Nuclear extract from HuT78 cells (endogenous SP140) was prepared following manufacturer’s protocol (Active Motif nuclear extract and co-IP kit). Protein concentration was determined by Bradford Assay (Pierce). Nuclear extracts were incubated with GSK675 (biotinylated GSK761) prebound to streptavidin Dynabeads (Thermo Fisher). Dynabeads were incubated with 2X SDS reducing buffer (Invitrogen) and the eluted proteins resolved on a 4-12% Bis-Tris SDS-PAGE (Invitrogen) with MagicMark (Invitrogen) and SeeBlue 2 (Invitrogen) protein ladders. The gel was subjected to Western blotting (Invitrogen) and incubated with Rabbit anti-SP140 antibody (HPA-006162; Sigma) followed by anti-rabbit IgG HRP (A4914; Sigma). The Western blot was developed with Super Signal West femto kit (Thermo Fisher) and imaged on Carestream chemilluminescence imager (Kodak). A chimeric gene encoding Halo-SP140 was synthesized using overlapping oligonucleotides and cloned in an expression vector pCDNA3.1. HEK293 cells were transfected with expression vector using Lipofectamine and incubated for 24h. The transfected cells were labelled with HaloTag TMR ligand following manufacturer’s protocol (HaloTag® TMR Ligand Promega). Nuclear extract from transfected HEK293 cells (Halo-SP140) was prepared and SP140 immunoprecipitation was carried out as described above. Eluted proteins were resolved on a 4-12% Bis-Tris SDS-PAGE gel (Invitrogen) with SeeBlue 2 (Invitrogen) protein ladder. Halo-SP140 was visualised using Versa Doc scanner and 520LP UV Transilluminator.

### BROMOscan® Bromodomain Profiling

BROMOscan® bromodomain profiling was provided by Eurofins DiscoverX Corp (Fremont, CA, USA, http://www.discoverx.com). Determination of the K_*d*_ between test compounds and DNA tagged bromodomains was achieved through binding competition against a proprietary reference immobilized ligand.

### Cell penetration assessment assay of GSK761

Intracellular GSK761 compound concentration was measured by RapidFire Mass Spectrometry utilizing the methodology described by^28^.

### Cell viability and cytotoxicity assays

M1 macrophages were generated *in vitro* from human primary CD14^+^ monocytes as described above. M1 macrophages were plated into opaque-walled 96-well plate at 10 × 10^5^ cells per well and incubated with a concentration gradient of GSK761 (0.04–1.11 μM) for 1h (0.1% DMSO was used as control). The cells were left unstimulated or stimulated with 100 ng/mL LPS for 24h. Cell viability was assessed using CellTiter-Glo® Luminescent Cell Viability Assay kit (Promega) according to the manufacturer’s protocol. This assay quantifies ATP, an indicator of metabolically active cells. An equal volume of freshly prepared CellTiter-Glo® reagent was added to each well the plate was shaken for 10 minutes at RT and luminescent signals were recorded using a plate reader (SpectraMax M5). The index of cellular viability was calculated as the fold change of luminescence with respect to untreated control cells.

Cell viability was also determined using Muse® Count & Viability Kit (Luminex corp) according to the manufacturer’s protocol and measured on the Guava® Muse® Cell Analyzer (Luminex corp).

### Histone H3 peptide displacement

Various H3 peptides (Anaspec library (https://www.anaspec.com/)) were serially diluted in DMSO (1% final assay concentration) and tested in the presence of 10 nM GSK306 (FAM-labelled version of GSK761) and 80 nM 6HisFLAGTEVSP140 (687-867) (2x apparent *K*_*d*_ for this ligand) in 50 mM HEPES (Sigma-Aldrich), pH 7.5, 50 mM NaCl (Sigma-Aldrich), 1 mM CHAPS (Sigma-Aldrich), 1 mM DTT (Sigma-Aldrich) in a total volume of 10 µl. Reactions were incubated for 30 minutes and FP was measured on a Perkin Elmer Envision multi-mode plate reader (PerkinElmer), by exciting the FAM fluorophore of GSK306 at a wavelength of 485 nm and then measuring emission at 535 nm in both parallel and perpendicular planes. The FP measurement, expressed as milliP (mP), was then calculated as described previously, and normalised to free- and bound-ligand controls to determine % response. IC_50_ values were calculated using the described four-parameter logistic equation.

### SP140/Histone Nano-bioluminescence resonance energy transfer™ (NanoBRET™) Assay Testing

The NanoBRET™ System is a proximity-based assay that can detect protein interactions by measuring energy transfer from a bioluminescent protein donor NanoLuc® (NL) to a fluorescent protein acceptor HaloTag® (HT). Briefly, SP140-NL and Hitone3.3-HT DNA was transfected into HEK293 cells using the following ratios: 1:1, 1:10 and 1:100, respectively. Signal window was determined by relative NL/HT-fused protein expression. Minus HT controls were used as the baseline in this experiment for each condition. The data is presented as NanoBRET response (mBU) which is dependent on the presence of the HT Ligand, and by microscopic imaging of NL/HT-fused protein signal. For more information about NanoBRET assay protocol: (https://www.promega.com/-/media/files/resources/protocols/technical-manuals/101/nanobret-proteinprotein-interaction-system-protocol.pdf?la=en)

### Flow cytometry (FACS)

Primary human CD14^+^ monocytes were positively selected from PBMCs using CD14 Microbeads according to the manufacturer’s instructions (Miltenyi Biotec) and differentiated for 3 days with 20 ng/mL M-CSF. The monocytes were then washed with PBS and treated with 0.1% DMSO or 0.04 µM GSK761. After 1h, 100 ng/mL IFN-γ (R&D systems) was added to the cells to generate M1 macrophages. After 3 days of incubation, the cells were harvested and subsequently permeabilized using 1% Saponin (Sigma-Aldrich) for 10 minutes on ice. Cells were then stained using the conjugated antibodies; PerCP-Cy5 mouse anti-human CD64 (dilution 1:50) (305023, Biolegend) and PE mouse anti-human CD206 (dilution 1:20) (2205525, Sony Biotechnology). The analysis was performed by flow cytometry (FACS) using the LSRFortessa and FACSCalibur (both BD Biosciences). FlowJo (BD LSRFortessa™ cell analyzer) was used for data analysis.

### Cytokine analysis

Cytokine expression in the supernatant of DMSO- or GSK761-treated M1 macrophages (LPS-stimulated or unstimulated) were measured using electro-chemiluminescence assays (Meso Scale Discovery [MSD])-Human ProInflammatory 7-Plex Tissue Culture Kit (IFN-γ, IL-1β, IL-6, IL-8, IL-10, IL-12p70, TNF) according to manufacturer’s protocols and analyzed on an MSD 1250 Sector Imager 2400 (Mesoscale). Supernatant IL-6, IL-8 and TNF from *SP140* siRNA or scrambled-treated M1 macrophages (LPS-stimulated or unstimulated) were determined by Sandwich Enzyme-Linked Immunosorbent Assay (ELISA; R&D systems) according to manufacturer’s protocol.

### Customized qPCR-Array

Human M1 macrophages were generated *in vitro* from human primary CD14^+^ monocytes as described above (from 5 independent donors). M1 macrophages were treated either with 0.1% DMSO or 0.04 µM GSK761 for 1h. The cells where then washed with PBS and stimulated with LPS for 4h. Total RNA was isolated using RNeasy Mini Kit (Qiagen) and treated with DNaseI (Qiagen) according to the manufacturer’s instructions. RNA was reverse transcribed using the First Strand Synthesis Kit (Qiagen) and loaded onto a customized RT^2^ profiler array for selected 89 genes according to the manufacturer’s instructions (Qiagen) and run on QuantStudio 7 Flex (software v1.0). Qiagen’s online GeneGlobe Data Analysis Center (https://geneglobe.qiagen.com/us/analyze/) was used to determine the DEGs. The data was presented as scatter plot. All data were normalized to the geometric mean of two reference genes (*RPL37A* and *ACTB)*. The list of genes included in this experiment were selected from DEG in *SP140* silenced M1 macrophages in ^10^ and from our MSD, qPCR and microarray datasets of *SP140* silenced M1 macrophages.

### Genome wide expression profiling (RNA-sequencing (RNA-seq))

M1 macrophages were generated *in vitro* from human primary CD14^+^ monocytes as described above (from 3 independent donors). M1 macrophages were incubated for 1h with either 0.1% DMSO or 0.04 µM GSK761. M1 macrophages were then kept unstimulated or stimulated with 100 ng/mL LPS (*E. coli* 0111:B4; Sigma) for 4h or 8h. Total RNA was isolated from macrophages using the RNAeasy mini kit (Qiagen) and transcribed into cDNA by qScript cDNA SuperMix (Quanta Biosciences) according to manufacturer’s instructions. Sequencing of the cDNA was performed on the Illumina HiSeq4000 to a depth of 35M reads at the Amsterdam UMC Core Facility Genomics. Quality control of the reads was performed with FastQC (v0.11.8) and summarization through MultiQC (v1.0). Raw reads were aligned to the human genome (GRCh38) using STAR (v2.7.0) and annotated using the Ensembl v95 annotation. Post-alignment processing was performed through SAMtools (v1.9), after which reads were counted using the featureCounts application in the Subread package (v1.6.3). Differential expression (DE) analysis was performed using DESeq2 (v1.24.0) in the R statistical environment (v3.6.3), in R2 Genomics Analysis and Visualization Platform-UMC (r2.amc.nl) and ShinyGO v0.60^27^.

### Chromatin immunoprecipitation (ChIP)

M1 macrophages were generated *in vitro* from human primary CD14^+^ monocytes as described above (10^7^ cells were used for each condition). M1 macrophages were either incubated for 1h with 0.1% DMSO or with 0.04 µM GSK761 and left unstimulated or simulated with 100 ng/mL LPS for 1h or 4h. The cells were cross-linked with 1% formaldehyde for 10 minutes at RT and quenched with 2.5 M glycine (Diagenode) for 5 minutes at RT. The ChIP assay was performed using the iDeal ChIP kit for Transcription Factors (Diagenode) and sonication was performed using the Picoruptor^™^ (Diagenode) according to the manufacturer’s protocols. Chromatin shearing was verified by migration on a 1% agarose gel (E-Gel, Thermo-Fisher) and visualised using E-Gel imager (Thermo-Fisher). Immunoprecipitation was performed with a polyclonal SP140 antibody (H00011262-M07, Abnova). DNA was purified using IPure kit (Diagenode) according to the manufacturer’s protocol. For validation, quantitative real-time ChIP-qPCR was performed on DNA isolated from input (unprecipitated) chromatin and SP140 ChIP DNA with primer pairs specific for the TSS of *TNF* and *IL6* genes. For detailed PCR primer sequences, please see supplemental table 1. Quantitative PCR was performed using SYBR Green (Applied Biosystems) and StepOnePLus (Applied Biosystems). Results were quantitated using the delta–delta CT (ΔΔCT) method. Libraries of input DNA and ChIP DNA were prepared from gel-purified >300–base pair DNA.

### Global profiling of chromatin binding sites

The DNA was used to generate sequencing libraries according to the manufacturer’s procedure (Life Technologies). The DNA was end polished and dA tailed, and adaptors with barcodes were ligated. The fragments were amplified (eight cycles) and quantified with a Bioanalyzer (Agilent). Libraries were sequenced using the HiSeq PE cluster kit v4 (Illumina) with the HiSeq 2500 platform (Illumina), resulting in 125-bp reads. FastQ files were adapter clipped and quality trimmed using BBDukF (java -cp BBTools.jar jgi.BBDukF -Xmx1g in=fastqbef.fastq.gz out=fastq.fastq minlen=25 qtrim=rl trimq=10 ktrim=r k=25 mink=11 hdist=1 ref=adapters.fa). Subsequently the reads were mapped to NCBI37/HG19 using Bowtie (bowtie2 -p 8 -x hg19 fastq.fastq > sample.sam) and sorted and converted into bam files using samtools. Peaks were called using MACS2, with the reads being extended to 200bp (callpeak --tempdir /data/tmp -g hs -B -t sample.bam -c control.bam -f BAM -n result/ --nomodel --extsize 149) and duplicates removed. BDG files were binned in 25bp regions and loaded in the R2 platform (r2.amc.nl) for subsequent analyses and visualization. For motif analyses, Summits obtained from the MACS2 analyses were extended by 250 bases and subsequently analyzed on a repeat masked hg19 genome with HOMER to identify over represented sequences (both known as well as de novo). The data was uploaded and analyzed in R2 Genomics Analysis and Visualization Platform-UMC (r2.amc.nl).

### SP140 inhibition in isolated human mucosal macrophages

Colon tissues were obtained during surgery procedures from patients with CD. The mucosa was stripped and dissociated in GentleMACS tubes in digestion medium (DM; complete medium (RPMI 1640 with PGA/L-glutamine/10% FCS) with 1 mg/mL Collagenase D (Roche), 1 mg/mL soybean Trypsin inhibitor (Sigma), 50 µg/mL DNase I (Roche)), and then mechanically dissociate on GentleMACS using program B01. The mucosa was then incubated in DM for 1h at 37°C while shaking. During digestion the dissociation was repeated two times. The tissue suspension was passed through a cell strainer (200-300 um) and centrifuged at 1500 rpm for 10 minutes. The dissociated cells were resuspended in cold MACS Buffer and macrophages were isolated using CD14 MicroBeads according to the manufacturer’s instructions (MiltenyiBiotec). The macrophages were incubated with either 0.1% DMSO or 0.04 µM GSK761 for 4h. RNA was extracted as described above and gene expression of *IL6, TNF, IL10* and *CD64* was measured by qPCR. All data were normalized to the reference gene *ACTB*.

### Statistics analysis

Statistical analysis was performed with GraphPad Prism v8.0.2.263 (GraphPad Software Inc.). For group analysis, data were subjected to one-way ANOVA or Student’s t-test. The two-tailed level of significance was set at p ≤ 0.05 (*), 0.01 (**), 0.001 (***) or 0.0001 (****) for group differences. Data is shown as mean ± SEM. The figures were prepared using Inkscape 0.92.4.

### Data Availability

Processed microarray data is publicly available and can be explored at r2.amc.nl under the data set name ‘Exp SP140 siRNA in M1 macrophages - deJonge - 12 - custom - ilmnht12v4’. The summary statistics for microarray data can be found in supplemental tables 3 and 4. Raw data of RNA-seq and ChIP-seq are publicly available and have been deposited in European Genome-phenome Archive (https://ega-archive.org/studies/EGAS00001004460). The counts and the summary statistics for RNA-seq data can be found in supplemental table 5, 6 and 7. The normalized read count for ChIP-seq data can be found in supplemental tables 8, 9, 10, 11, 12 and 13.

## Supplementary figure legends

**Supplementary figure 1:**
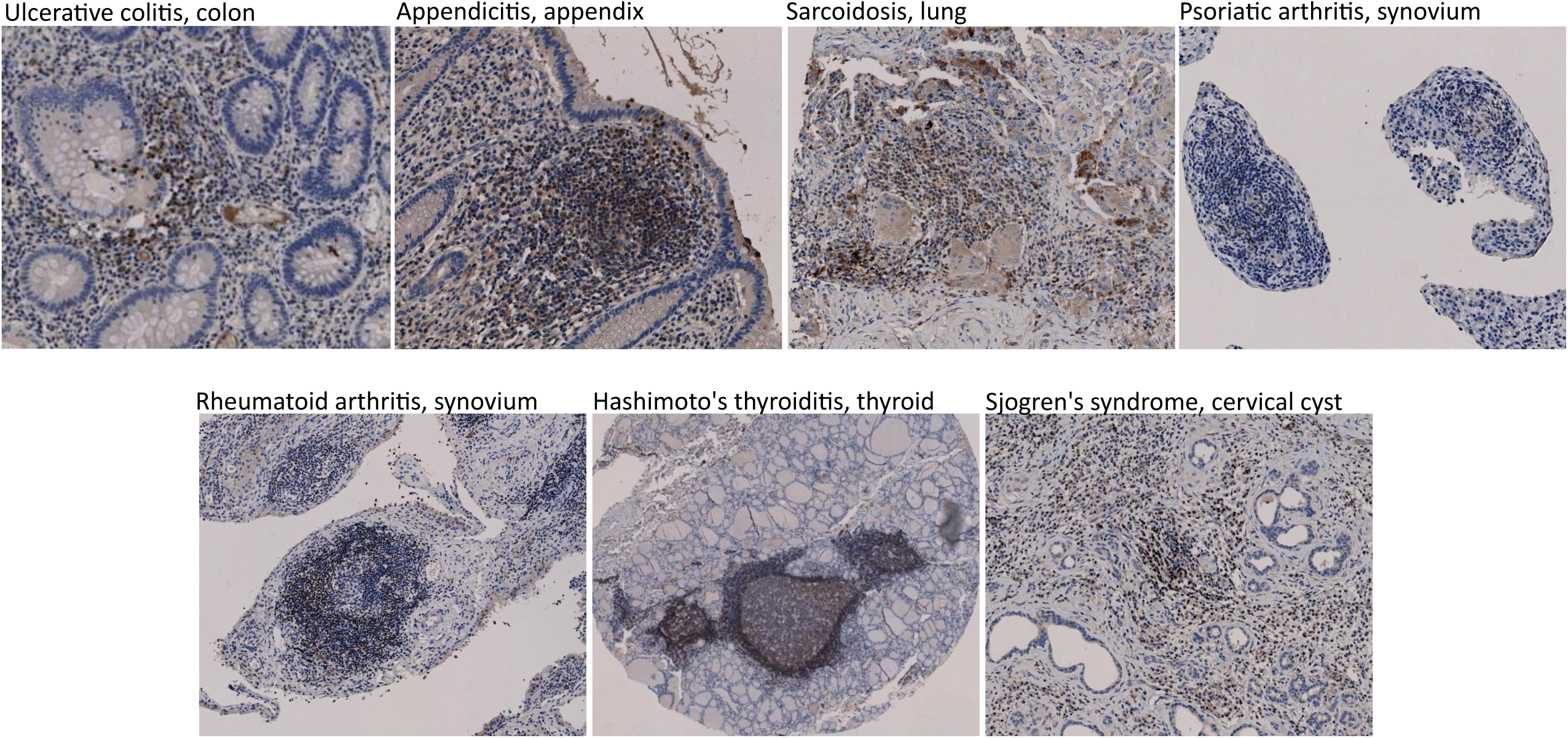
SP140 expression is associated with inflammatory diseases. Immunohistochemistry of SP140 in ulcerative colitis (colon), appendicitis (appendix), sarcoidosis (lung), psoriatic arthritis (synovium), rheumatoid arthritis (synovium), Hashimoto’s thyroiditis (thyroid) and Sjogren’s syndrome (cervical cyst). SP140 is illustrated by peroxide staining.

**Supplementary figure 2:**
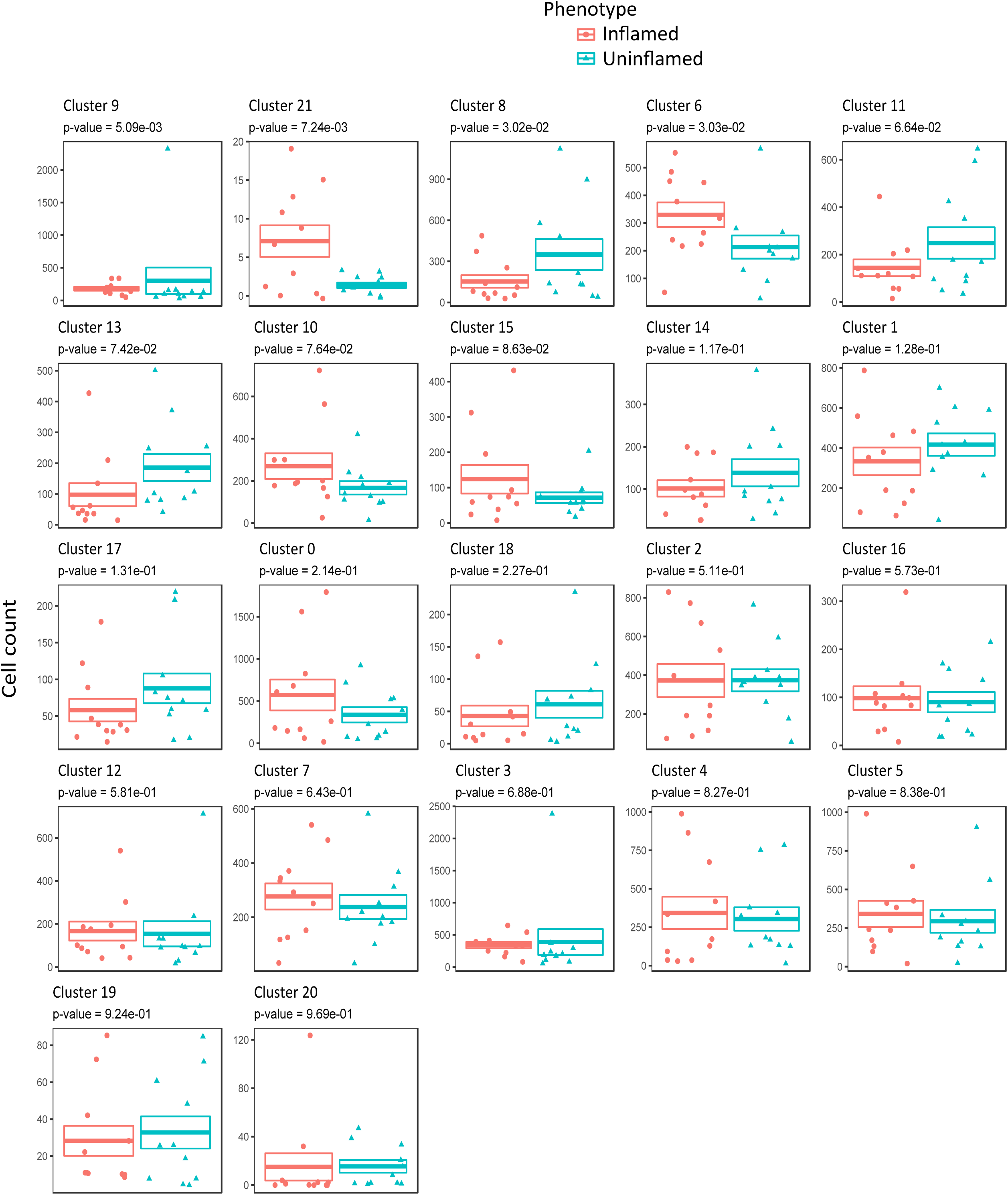
CD-ileal inflamed tissue has consistently more cells than uninflamed tissue in cluster 6. Publicly available single-cell RNA-sequencing was used to investigate *SP140* expression in ileal macrophages in inflamed (n=11) and uninflamed (n=11) biopsies of CD patients. Comparative analyses of the cell counts per cluster when comparing inflamed with uninflamed. Graphs are ranked by p-values, which were calculated through the Wald test as implemented in DESeq2.

**Supplementary figure 3:**
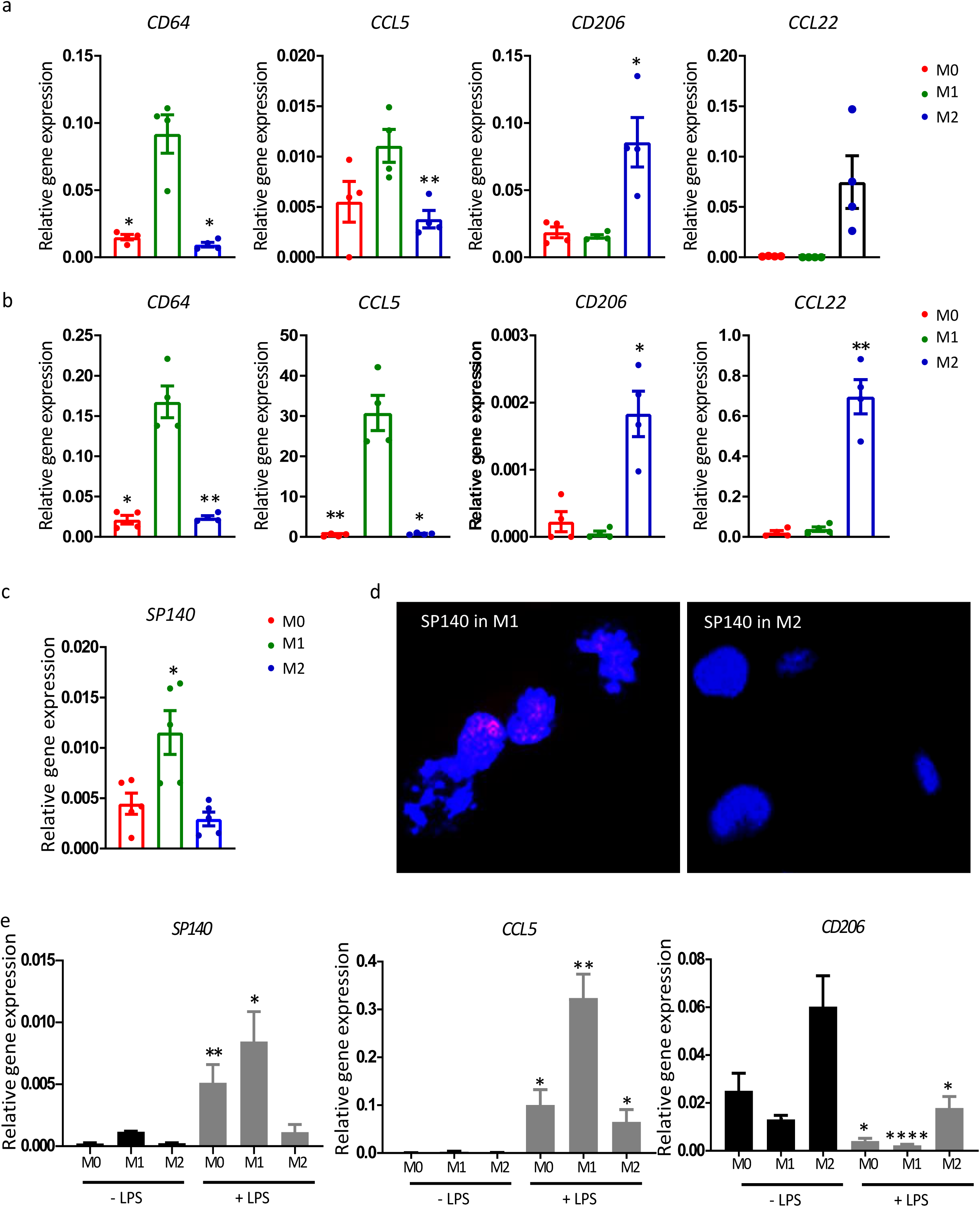
Inflammatory stimulus induces SP140 expression. (**a**,**b)** Relative gene expression of *CD64, CCL5, CD206* and *CCL22* in M0, M1 and M2 macrophages derived from **(a)** human primary CD14^+^ monocytes or **(b)** from THP-1 cells, n=4. **(c)** Relative gene expression of *SP140* in M0, M1 and M2 macrophages derived from THP-1 cells, n=5. **(d)** Immunofluorescence staining of SP140 in M1 and M2 macrophages derived from THP-1 cells. DAPI staining nucleus in blue and SP140 speckles in red. **(e)** Relative gene expression of *SP140, CCL5* and *CD206* after 24h of 100 ng/mL LPS-stimulated or unstimulated M0, M1 and M2 macrophages derived from human primary CD14^+^ monocytes, n=4. Relative gene expression was measured by qPCR. Statistical significance is indicated as follows: \**P* < 0.05, \*\**P* < 0.01, \*\*\**P* < 0.001, \*\*\*\**P* < 0.0001.

**Supplementary figure 4:**
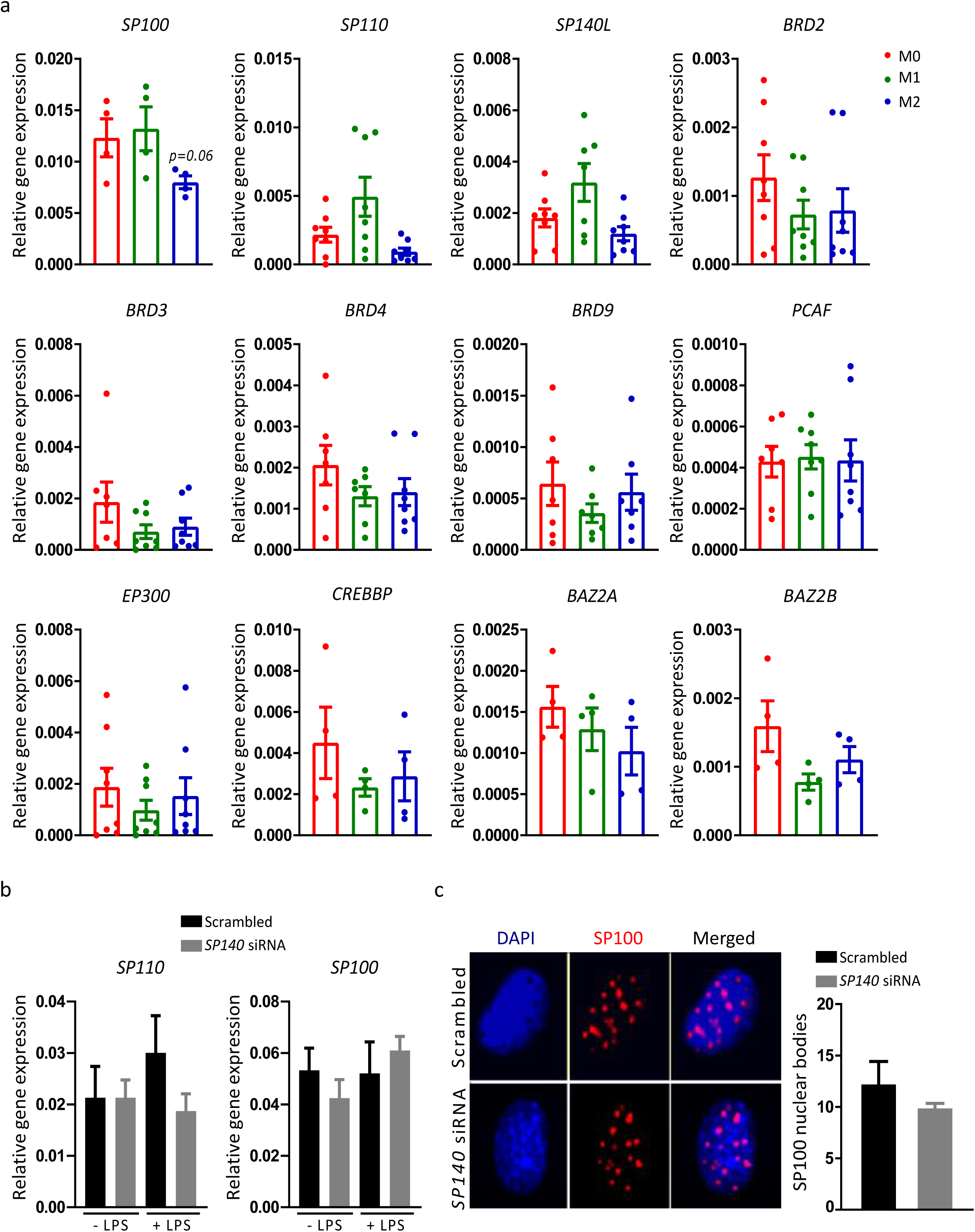
The association between *SP140* and inflammatory macrophages is not a common phenomenon amongst BCPs. **(a)** Relative gene expression (qPCR) of 12 BCPs; *SP100, SP110, SP140L, BRD2, BRD3, BRD4, BRD9, PCAF, EP300, CRBBP, BAZ2A* and *BAZ2B* in M0, M1 and M2 macrophages derived from human primary CD14^+^ monocytes, n= 4 to 8. **(b)** Relative gene expression of *SP110* and *SP100* in M1 macrophages, transfected with siRNA against *SP140* (grey bars) or a scrambled control siRNA (black bars), and then stimulated or not with LPS (100 ng/mL for 4h). **(c)** Immunofluorescence staining of SP140 in M1 macrophages, transfected with siRNA against *SP140* or a scrambled control siRNA, and then stimulated with LPS (100 ng/mL for 24h). SP140 speckles (red dots) were imaged and SP140 nuclear bodies were counted. DAPI (blue) was used to stain the nucleus.

**Supplementary figure 5:**
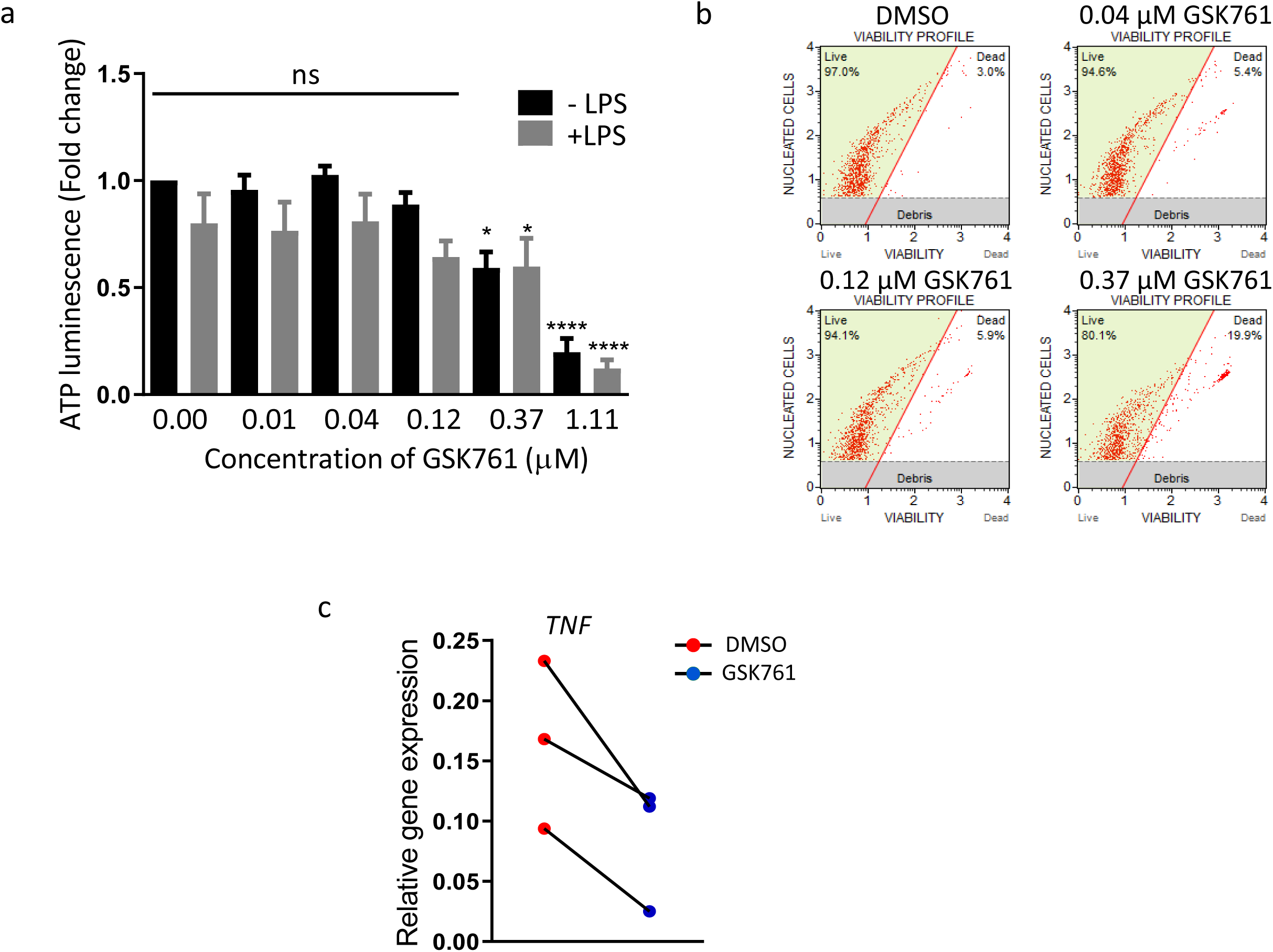
GSK761 lowers the expression of the M1 macrophage polarization marker *TNF*. **(a)** ATP luminescence measurement within M1 macrophages treated with an increasing concentration of GSK761 (0.01, 0.04, 0.12, 0.37. 1.11 µM) or 1% DMSO in presence or absence of 100 ng/mL of LPS for 24h, n=6, statistical significance is indicated as follows: \**P* < 0.05, \*\**P* < 0.01. The Y axis indicates the fold change in detected ATP luminescence (relative to unstimulated DMSO control). **(b)** M1 macrophages were pretreated with 0.1% DMSO or with GSK761 (0.04 µM, 0.12 µM or 0.37 µM), and then stimulated with 100 ng/mL for 24h. The cells were stained with Muse® Count & Viability Reagent, and then analyzed on the Muse® Cell Analyzer. The data shown as viability vs nucleated cells and presented as % viability. **(c)** Primary human CD14^+^ monocytes were differentiated with 20 ng/mL M-CSF for 3 days. The cells were then washed with PBS and treated with 0.1% DMSO or 0.04 µM GSK761 for 1h prior to 3 days polarization to M1 phenotype with 100 ng/mL INF-γ (DMSO and GSK761 were not washed and kept in the culture during the 3 days of polarization). *TNF* gene expression was measured by qPCR, n=3.

**Supplementary figure 6:**
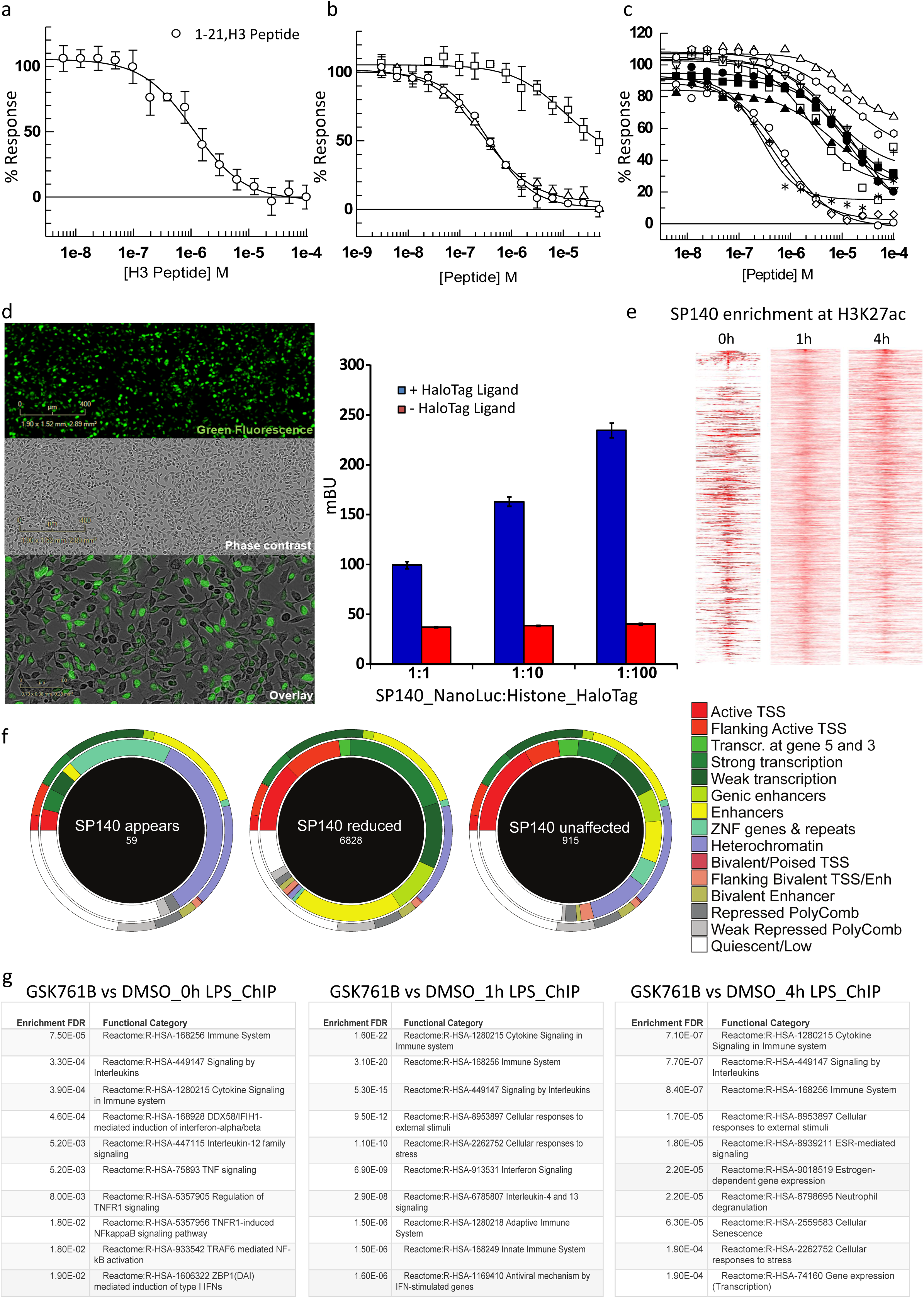
SP140 protein binds to histone H3. **(a)** IC50 (0.36 +/- 0.03 µM) for the displacement of GSK761 from SP140 by H3 peptide (1-21, ARTKQTARKSTGGKAPRKQLA). **(b)** IC50 for the displacement of GSK761 by H3 peptide (circles) compared to IC50s for H3 peptide tri-methylated at position K4 (1-21, H3K4(3Me)) (Squares) and H3 peptide tri-methylated at position K14 (1-21, H3K14(3Me)) (10 +/- 1 and 0.28 + 0.03 µM, respectively). **(c)** IC50s for the displacement of a range of modified H3 and truncated peptides. Open circles; Biotinylated H3 peptide (1-21, IC50 = 0.51 µM), open squares; Biotinylated H3K9(Ac) (1-21, IC50 = 2.22 µM), open triangles up; Biotinylated H3K4(3Me)K9(Ac) (1-21, IC50 = 11.32 µM), open triangles down; Biotinylated H3K4(3Me) (1-21, IC50 = 17.66 µM), open diamonds; biotinylated H3K14(Ac) (1-21, IC50= 0.43 µM), open octagon; H3K4(3Me) (1-10, IC50 = 12.47 µM), plus symbol; H3K4(3Me) (1-21, IC50 = 6.10 µM), star symbol; biotinylated H3(+YCK) (1-18 +YCK, IC50 = 0.27 µM), solid circle; H3K4(Ac)K14(3Me) (1-21, IC50 = 13.10 µM), solid square; H3K4(3Me)K14(Ac) (1-21 IC50 = 9.99 µM and solid triangle up; biotinylated H3 (1-9 IC50 = 2.83 µM). **(b)** Peptides tested and data not shown (due to no fit); Biotinylated H3K14(Ac) (9-20), Biotinylated H3 (15-40) and Biotinylated H3 (5-24). **(d)** SP140-NL and Histone3.3-HT DNA was transfected into HEK293 cells. NL/HT-fused protein green signal was imaged by microscopy (left) and NanoBRET response (mBU) was measured (right). **(e)** SP140 enrichment at H3K27ac after 0h, 1h and 4h of 100 ng/mL LPS-stimulation of M1 macrophages. **(f)** Roadmap comparing the DMSO versus GSK761 on the peaks that were called in both instances (In 1h LPS-stimulated M1 macrophages). If the peaks in both conditions are overlapping, then we call them unaffected (a peak is found under both conditions) (right). If there was a peak in DMSO, but in the presence of GSK761, SP140 binding is reduced or absent, then this is described as ‘SP140 reduced’ (middle). Finally, if there was no peak called in DMSO samples, but a peak appears in the GSK761 condition, then SP140 binding is described as ‘SP140 appears’ (left). For the circles: All has been compared to the epigenome roadmap data for 1 particular tissue (primary monocytes from peripheral blood). This provides an annotation for every 200bp region in the genome and states its association in 15 different categories. The narrow outer circle reflects the proportion of the annotated genome in that particular tissue (primary monocytes from peripheral blood). The thicker inner circle reflects the proportions of the annotated genome where SP140 binding is seen (basepair with the highest signal in a peak) for the peaks as termed above. The numbers inside the circles reflect the number of SP140 enriched genes. **(g)** Reactome pathway analysis of SP140 DBGs in unstimulated, 1h LPS- and 4h LPS-stimulated M1 macrophages comparing 0.1% DMSO and 0.04 µM GSK761.

**Supplementary figure 7:**
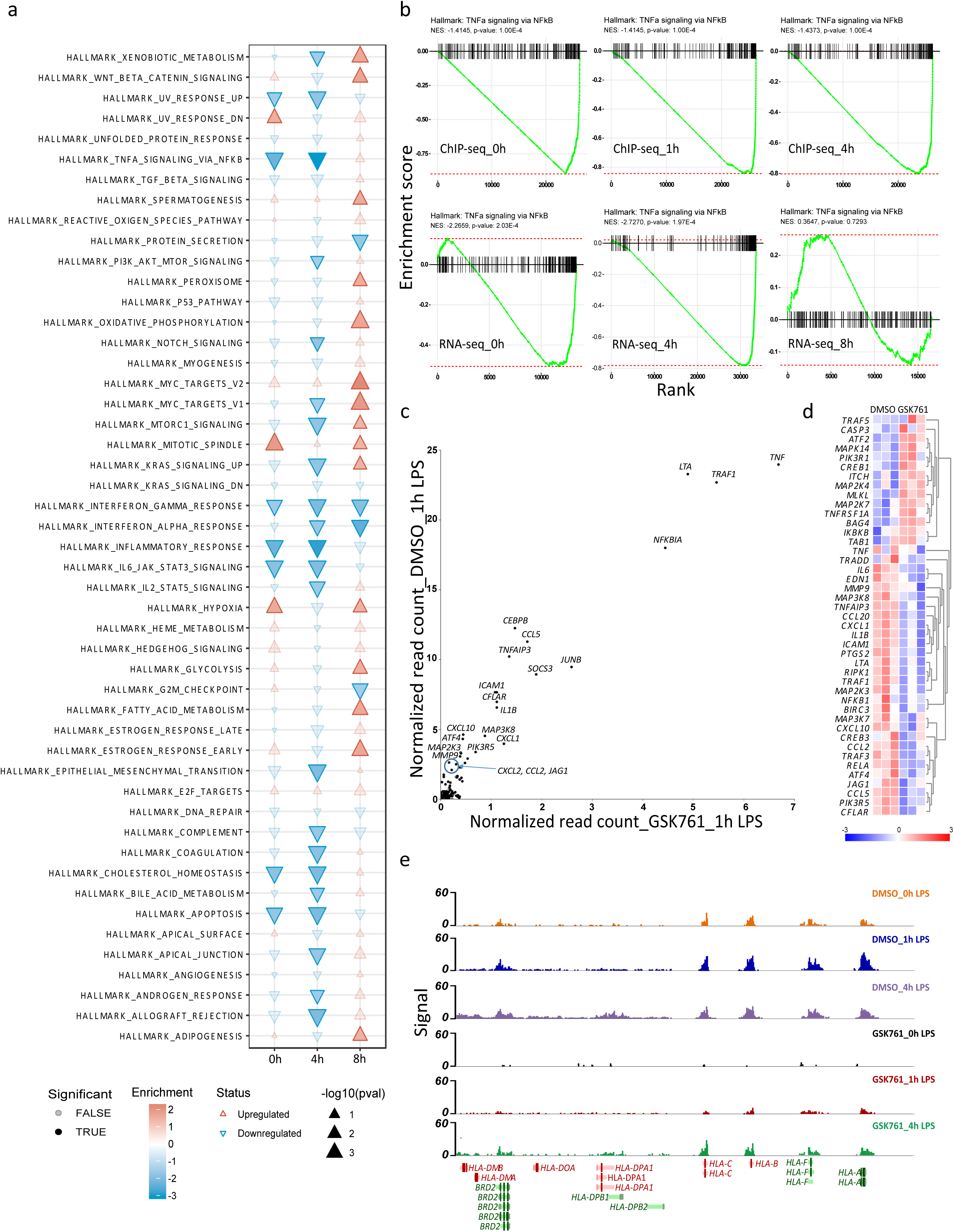
GSK761 prevents SP140 enrichment at gene set involved in TNF-signaling and antigen presentation. **(a)** A complete data set of hallmark pathway analysis for global DEGs in LPS-unstimulated or stimulated (4 or 8h) M1 macrophages (pretreated with 0.1% DMSO or 0.04 µM GSK761). The direction and color of the arrow indicates the direction and size of the enrichment score, the size of the arrow is proportional to the –log_10_(*p*-value), and non-transparent arrows represents significantly affected pathways,n=3. **(b)** An enrichment pathway analysis targeting TNF-signaling for SP140 ChIP-seq (top) and RNA-seq (bottom) comparing 0.1% DMSO-with 0.04 µM GSK761 treated-M1 macrophages at 0, 1 and 4h of 100 ng/mL LPS-stimulation. **(c)** R2 TSS-plot targeting TNF-signaling pathway genes, comparing 0.1% DMSO with 0.04 µM GSK761 treated M1 macrophages after 1h of 100 ng/mL LPS-stimulation. The indicated genes in the graph present the most DBGs. **(d)** Heatmap of most DEGs that are involved in TNF-signaling, comparing 0.1% DMSO with 0.04 µM GSK761 treated M1 macrophages after 4h of 100 ng/mL LPS-stimulation.**(e)** SP140 ChIP-seq genome browser view of *HLA-DMB, HLA-DMA, HLA-DQA, HLA-DPA1, HLA-DPB1, HLA-DPB2, HLA-C, HLA-B, HLA-F, HLA-A* and *BRD2*. Y axis represents signal score of recovered sequences in 0.1% DMSO and 0.04 µM GSK761 treated macrophages after 0, 1 and 4h of 100 ng/mL LPS-stimulation.

**Supplementary figure 8:**
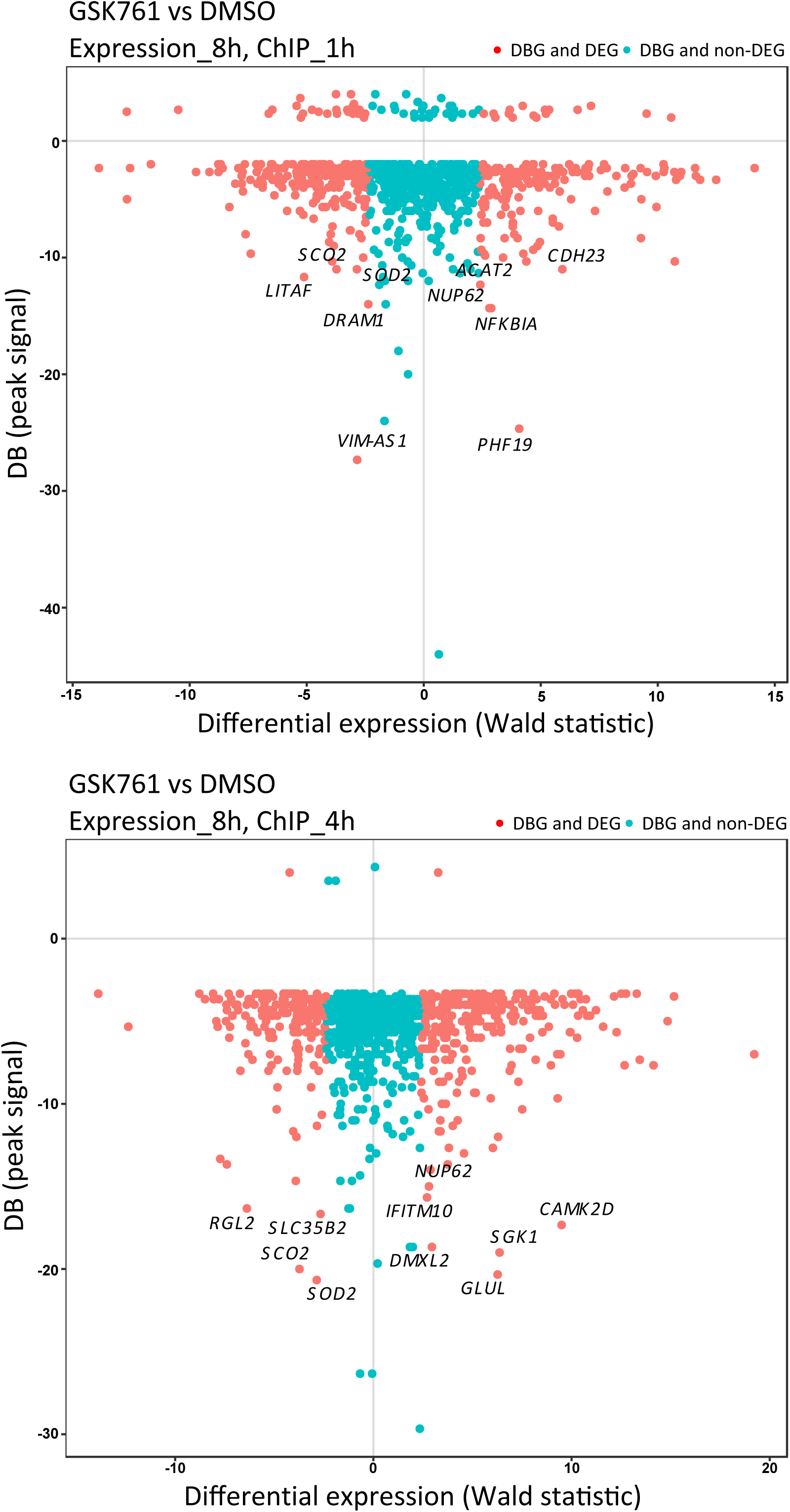
Genes that are involved in innate immune response showed the strongest concordant differential SP140 binding and gene expression. Comparative analyses of the top 1000 DBGs (signal) with their gene expression (Wald statistic). Gene expression at 8h LPS and ChIP at 1h LPS (top) and gene expression at 8h LPS and ChIP at 4h LPS (bottom). Y axis represents the SP140 differential binding signal (BD).

**Supplementary figure 9:**
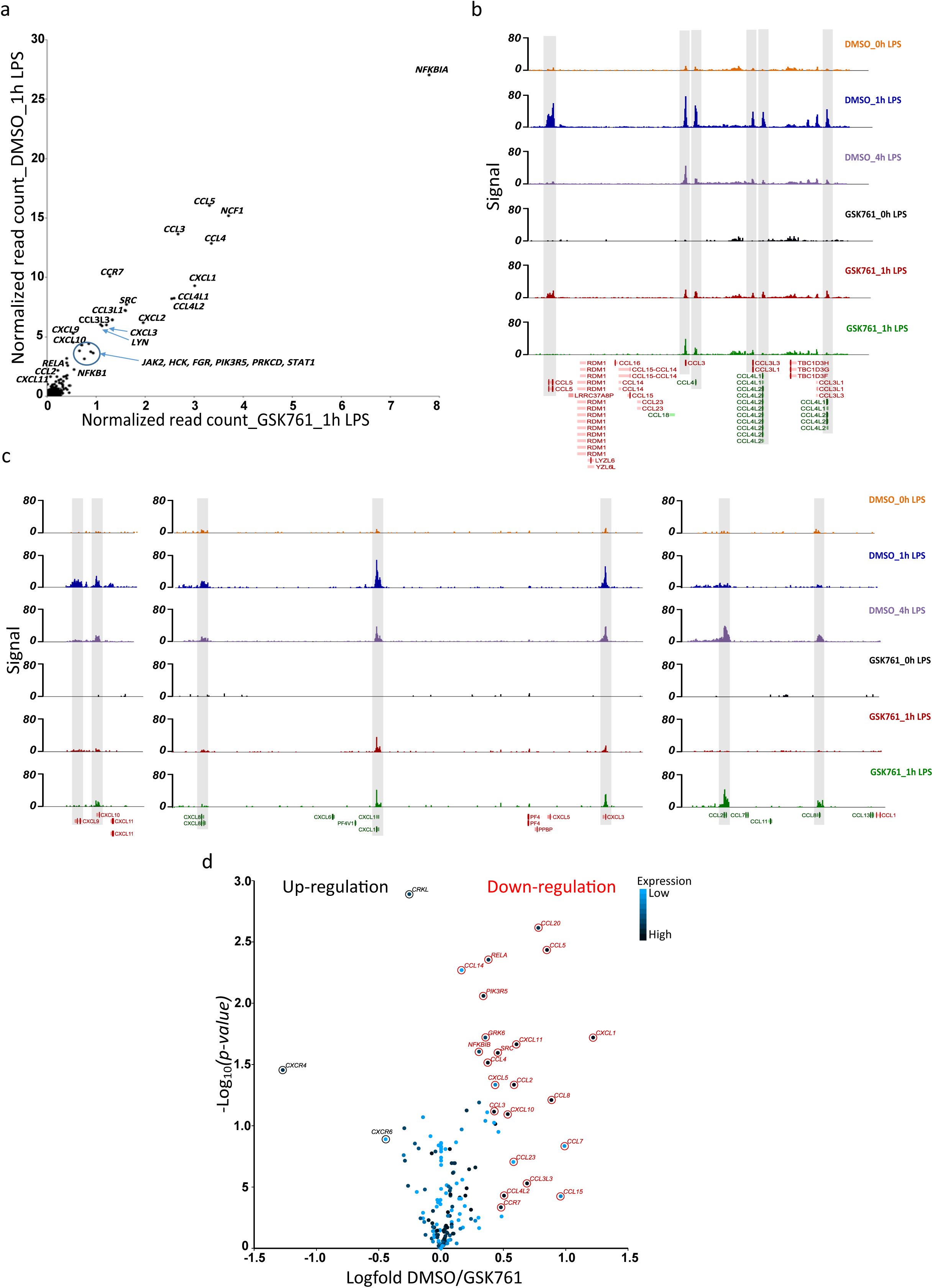
GSK761 reduces SP140 enrichment at several chemokine genes. **(a)** R2 TSS-plot of chemokine-signaling genes, comparing 0.1% DMSO with 0.04 µM GSK761-treated M1 macrophages after 1h of 100 ng/mL LPS-stimulation. The indicated genes in the graph represent the most DBGs involved in chemokine activity/response. **(b, c**) SP140 ChIP-seq genome browser view of some of most SP140 differentially bound chemokines at different time points of LPS-stimulation in M1 macrophages.SP140-bound chemokines were highlighted. **(d)** Volcano plot of DEGs involved in chemokine signaling/genes after 1h of 100 ng/mL LPS-stimulation.

## Supplementary table legends

**Supplemental table 1.**
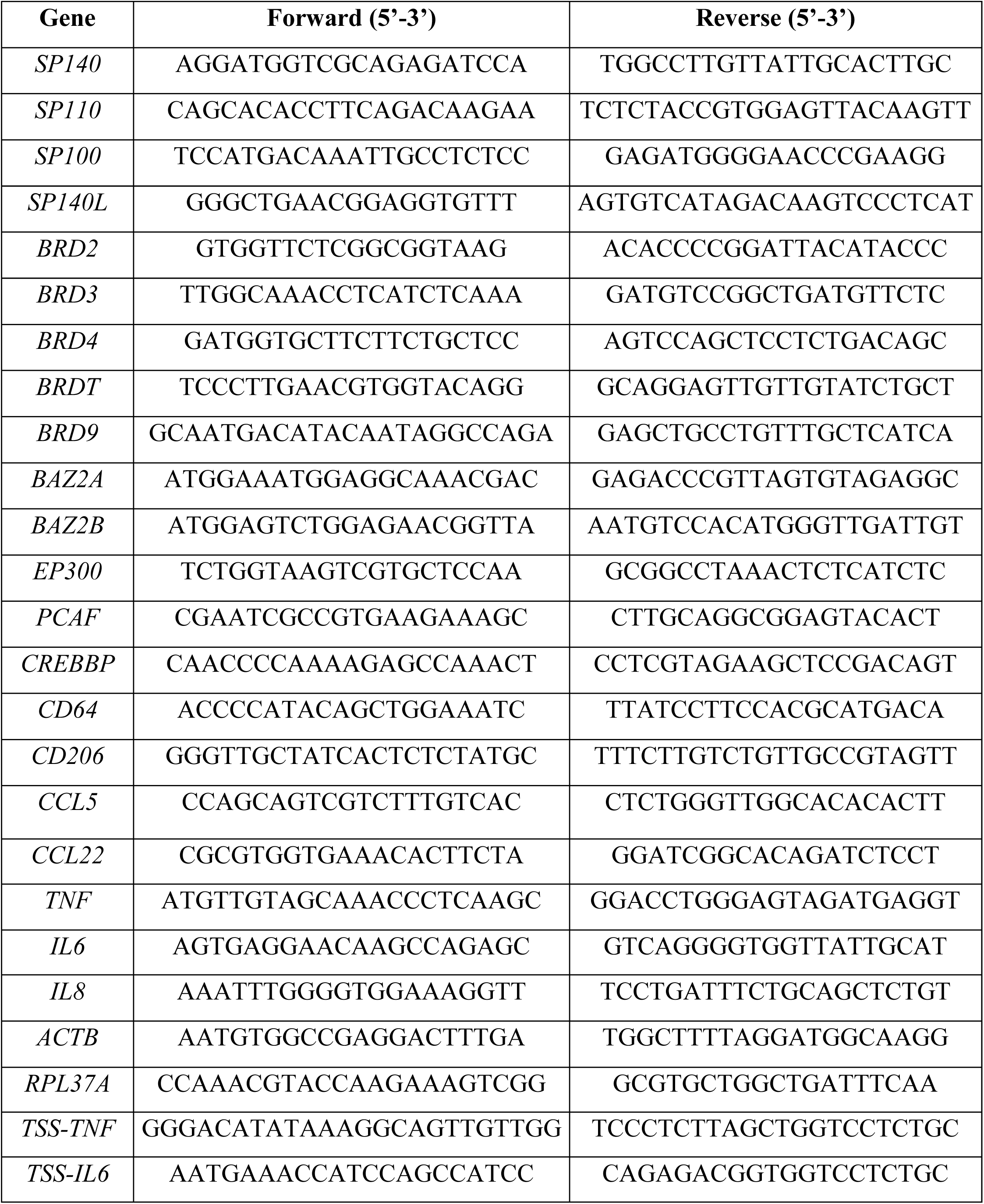
Primer sequences used in the quantitative PCR analysis of the genes of interest.

**Supplemental table 2.**
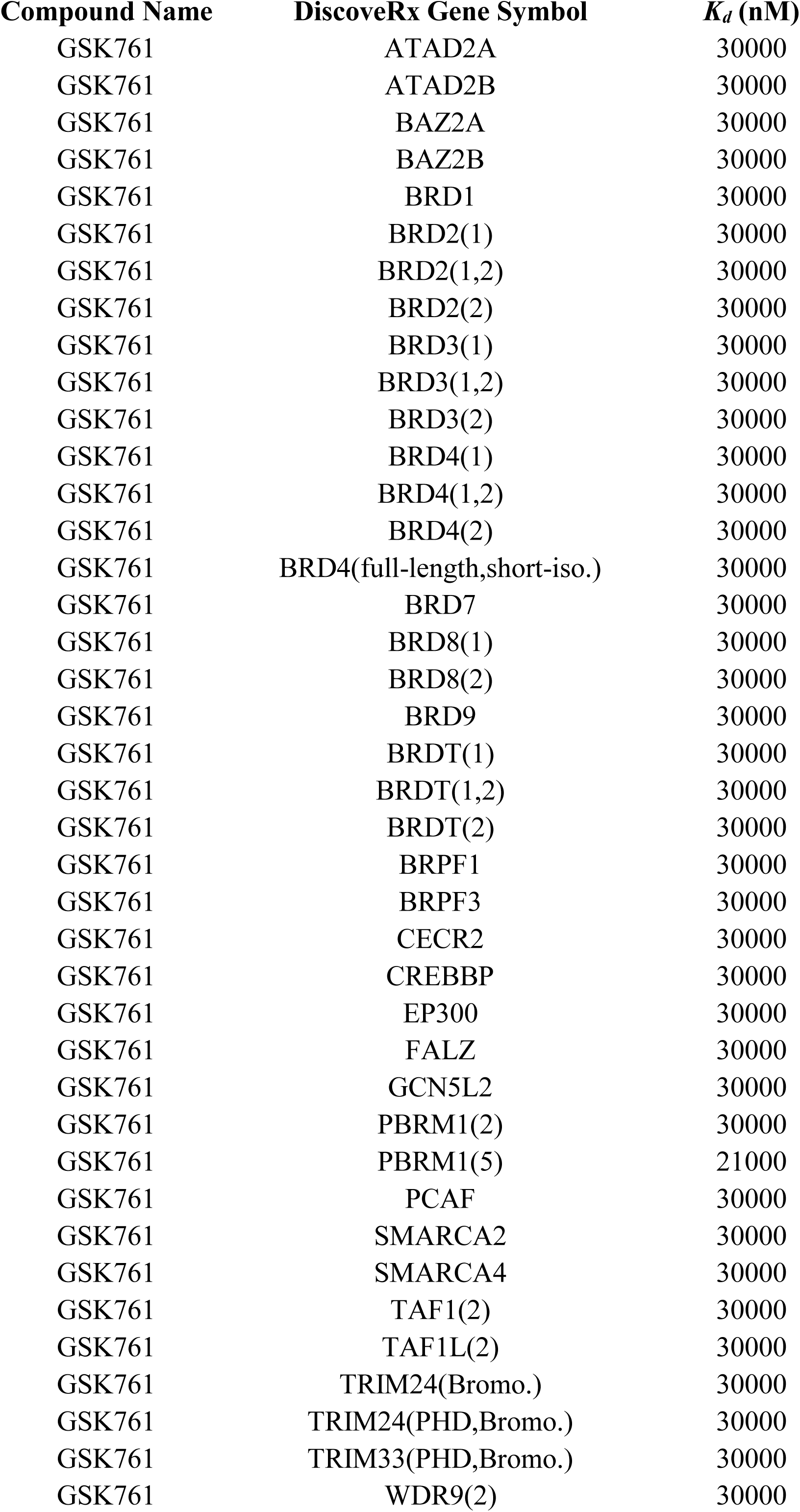
GSK761 screen against human BCPs (Bromoscan assay) reveals no binding at *Kd* of ≤ 30000 nM (for most of tested BCPs) and at *Kd* of ≤ 21000 nM for PBRM1(5), indicating a high degree of specificity of GSK761 for SP140

| Compound Name | DiscoverX Gene Symbol | $K_d$ (nM) |
| --- | --- | --- |
| GSK761 | ATAD2A | 30000 |
| GSK761 | ATAD2B | 30000 |
| GSK761 | BAZ2A | 30000 |
| GSK761 | BAZ2B | 30000 |
| GSK761 | BRD1 | 30000 |
| GSK761 | BRD2(1) | 30000 |
| GSK761 | BRD2(1,2) | 30000 |
| GSK761 | BRD2(2) | 30000 |
| GSK761 | BRD3(1) | 30000 |
| GSK761 | BRD3(1,2) | 30000 |
| GSK761 | BRD3(2) | 30000 |
| GSK761 | BRD4(1) | 30000 |
| GSK761 | BRD4(1,2) | 30000 |
| GSK761 | BRD4(2) | 30000 |
| GSK761 | BRD4(full-length,short-iso.) | 30000 |
| GSK761 | BRD7 | 30000 |
| GSK761 | BRD8(1) | 30000 |
| GSK761 | BRD8(2) | 30000 |
| GSK761 | BRD9 | 30000 |
| GSK761 | BRDT(1) | 30000 |
| GSK761 | BRDT(1,2) | 30000 |
| GSK761 | BRDT(2) | 30000 |
| GSK761 | BRPF1 | 30000 |
| GSK761 | BRPF3 | 30000 |
| GSK761 | CECR2 | 30000 |
| GSK761 | CREBBP | 30000 |
| GSK761 | EP300 | 30000 |
| GSK761 | FALZ | 30000 |
| GSK761 | GCN5L2 | 30000 |
| GSK761 | PBRM1(2) | 30000 |
| GSK761 | PBRM1(5) | 21000 |
| GSK761 | PCAF | 30000 |
| GSK761 | SMARCA2 | 30000 |
| GSK761 | SMARCA4 | 30000 |
| GSK761 | TAF1(2) | 30000 |
| GSK761 | TAF1L(2) | 30000 |
| GSK761 | TRIM24(Bromo.) | 30000 |
| GSK761 | TRIM24(PHD,Bromo.) | 30000 |
| GSK761 | TRIM33(PHD,Bromo.) | 30000 |
| GSK761 | WDR9(2) | 30000 |

**Supplemental table 3:** Summary statistics for microarray data (*SP140* siRNA M1 macrophages vs scrambled siRNA-M1 macrophages)

**Supplemental table 4:** Summary statistics for microarray data (*SP140* siRNA-M1 macrophages stimulated with 4h LPS (100 ng/mL) vs scrambled siRNA-M1 with 4h LPS (100 ng/mL))

**Supplemental table 5:** Counts and summary statistics for RNA-seq data (GSK761-pretreated M1 macrophages vs DMSO-pretreated M1 macrophages)

**Supplemental table 6:** Counts and summary statistics for RNA-seq data (GSK761-pretreated M1 macrophages stimulated with 4h LPS (100 ng/mL) vs DMSO-pretreated M1 macrophages stimulated with 4h LPS (100 ng/mL))

**Supplemental table 7:** Counts and summary statistics for RNA-seq data (GSK761-pretreated M1 macrophages stimulated with 8h LPS (100 ng/mL) vs DMSO-pretreated M1 macrophages stimulated with 8h LPS (100 ng/mL))

**Supplemental table 8:** The normalized read count for ChIP-seq (DMSO-pretreated M1 macrophages)

**Supplemental table 9:** The normalized read count for ChIP-seq (DMSO-pretreated M1 macrophages stimulated with 1h LPS (100 ng/mL))

**Supplemental table 10:** The normalized read count for ChIP-seq (DMSO-pretreated M1 macrophages stimulated with 4h LPS (100 ng/mL))

**Supplemental table 11:** The normalized read count for ChIP-seq (GSK761-pretreated M1 macrophages)

**Supplemental table 12:** The normalized read count for ChIP-seq (GSK761-pretreated M1 macrophages stimulated with 1h LPS (100 ng/mL))

**Supplemental table 13:** The normalized read count for ChIP-seq (GSK761-pretreated M1 macrophages stimulated with 4h LPS (100 ng/mL))

## Supplementary information (compound discovery and synthesis)

### Screening of DNA encoded libraries (GSK761 discovery)

Affinity screening of DNA Encoded Library (DEL) was done using AntiFlag matrix tips (Phynexus). Before use, the AntiFlag matrix tips were washed four times with 1x selection buffer and stored at 4° C. The SP140 protein (24 kDa) used for the affinity selection was: 6His-Flag-Tev-SP140 (687-867). During the screening process, each AntiFlag matrix tip was loaded with 5 μg of protein and the No target tips was treated with buffer only. The following 1x selection buffer was used in the affinity selection: 50 mM HEPES (7.5), 150 mM NaCl, 0.1% Tween 20, 1 mM BME, 1 mg/mL sheared salmon sperm DNA (sssDNA, Ambion). Three rounds of affinity selection were performed for each screening condition. No target controls (buffer only) were run in parallel as a control.

Selection Round 1: Prior to initiating target selections, 5 μg of SP140 (aa 687-867) protein was immobilized on prepared Anti-Flag resin tip (Phynexus), tips were washed four times with 100 μL of 1 x selection buffer. For each selection, 5 nM of DEL molecules (pool DEL 34-79) in 60 μL of 1 x selection buffer was incubated with the immobilized SP140 (aa 687-867) by pipetting up and down for 1h (RT). Tips were then washed eight times with 100 μL of 1x Selection buffer, and another 2 times with 100 μL of DNA free 1x Selection buffer. In order to release the bound DEL molecules off the tip, a heat elution was performed by treating the tip in 60 μL of 1 x selection buffer (minus sssDNA) for 12 minutes at 80 °C. Eluted samples were post cleared to remove denatured protein by passing over a fresh IMAC resin tip to remove any denatured protein for 10 minutes at RT. This step was repeated once. 1 μL of round 1 elution was retained to be used for qPCR. sssDNA and buffer were added to bring the total volume of the eluted material to 60 uL to be used for next round of selection.

Selection Round 2: The 2^nd^ round selection was performed by binding 5 μg of fresh SP140 (aa 687-867) protein to a fresh prepared Anti-Flag tip. The above selection procedure was repeated using the eluted material from round 1. At the end of round 2, 5 μL of the elution was retained. The eluted material was post cleared twice to remove denatured protein, as described above. sssDNA and buffer were added to bring the volume to 60 μL in order to begin round 3 of the selection.

Selection Round 3: The above selection procedure was repeated with eluted material from round 2. However, no post-clear step was performed for round 3 selection. At the end of round 3 selection, a quantitative PCR was run to assess yield from each round of selection. Target and No target samples from Round 3 elution were sequenced on an Illumina sequencer.

### Cloning, expression and purification

DNA spanning the SP140 region encoding the BRD and PHD domains (encoding SP140 amino acids 687-867) was prepared in a construct, which was subsequently inserted into a construct that comprised His_6_ and FLAG (DYKDDDDK) affinity chromatography peptides and a TEV protease cleavage site (ENLYFQ\S, “\” denotes the cleaved peptide bond). This DNA sequence was then cloned into a ET11c vector (Bioduro) and then subsequently used to transform *E. coli* BL21 CodonPlus (DE3) RIPL (Stratgene) in media containing 100 μg/mL ampicillin and 34 μg/mL chloramphenicol as selection antibiotics. A culture in Luria Bertani (LB) medium was grown at 30°C, 240 rpm, overnight in Erlenmeyer shake flasks to provide a seed for a larger expression culture in auto-inducible Overnight Express medium (supplemented with 1% Glycerol and 0.1 mM ZnCl_2_), in shake flasks, at 37 °C and 200 rpm. Incubation temperature was reduced to 25°C when O.D.600nm was 1.2. The culture was pelleted by centrifugation after a further 20h incubation and stored at −80° C. For protein purification, a 40 g cell pellet was processed by mixing with 200 mL of PBS, 50 mM Imidazole, 10% Glycerol, PIC III, 1 mg/mL lysozyme, Benzonase (Merck) pH7.4 and left to stir for 30 minutes at 4°C. The resulting suspension was lysed by sonication on ice for 15 minutes (10 seconds on /10 seconds off) and then harvested by centrifugation for 90 minutes at 30000 rpm. The lysate supernatant was collected and then passed through a 20 mL HisTrap HP column (GE Healthcare) and then washed back with buffer A. (Buffer A: PBS, 2 mM DTT, 10% Glycerol, pH 7.4). Bound SP140 was eluted using stepped protocol with buffer B (PBS, 2 mM DTT, 10% Glycerol, 500 mM Imidazole, and pH 7.4) yielding 434.4 mg of total protein. This pool was diluted down to 5 mScm using 20 mM Hepes, 2 mM DTT, 10% Glycerol, and pH 7.5 and further purified using 100 mL Source15Q anion exchange. Elution of the SP140 was carried out using a segmented linear NaCl gradient (0-50% buffer B over 20 column volumes (Cvs), 50-100% over 1 Cv). Three peaks were obtained from the post anion exchange material (Peak 1, 2 and 3) and pooled separately. The pooled fractions from absorbance peak 2 were further purified using gel filtration (Superdex 75 320 mL Cv), which lead to a final yield of purified SP140 protein of 7.87 mg, in a buffer comprising 10 mM Potassium phosphate, 100 mM NaCl, 0.5 mM TCEP, 5% Glycerol, pH7.4.

### Compound synthesis

Unless otherwise stated, all reactions were carried using anhydrous solvents. Solvents and reagents were purchased from commercial suppliers and used as received. Reactions were monitored by LCMS. Silica flash chromatography was carried out using SP4 apparatus using RediSep® pre-packed silica cartridges. Ion exchange chromatography was carried out using Biotage Isolute cartridges and extracted organic mixtures were dried using Biotage PTFE hydrophobic phase separator frits unless otherwise stated. NMR spectra were recorded at RT (unless otherwise stated) using standard pulse methods on a Bruker AV-400 spectrometer (^1^H = 400 MHz, ^13^C = 101 MHz). Chemical shifts are referenced to trimethylsilane (TMS) or the residual solvent peak, and are reported in ppm. Coupling constants are reported in Hz and refer to ^3^J_H-H_ couplings, unless otherwise stated. Coupling constants are quoted to the nearest 0.1 Hz and multiplicities are given by the following abbreviations and combinations thereof: s (singlet), d (doublet), t (triplet), q (quartet), m (multiplet), br. (broad). LCMS analysis was carried out on a Waters Acquity UPLC instrument equipped with a BEH or CSH column (50 mm x 2.1 mm, 1.7 μm packing diameter) and Waters micromass ZQ MS using alternate-scan positive and negative electrospray. Analytes were detected as a summed UV wavelength of 210 – 350 nm. Two liquid phase methods were used: **Formic**: 40 °C, 1 mL/min flow rate. Gradient elution with the mobile phases as (A) water containing 0.1% volume/volume (v/v) formic acid and (B) acetonitrile containing 0.1% (v/v) formic acid. Gradient conditions were initially 1% B, increasing linearly to 97% B over 1.5 min, remaining at 97% B for 0.4 min then increasing to 100% B over 0.1 min. **High pH**: 40 °C, 1 mL/min flow rate. Gradient elution with the mobile phases as (A) 10 mM aqueous ammonium bicarbonate solution, adjusted to pH 10 with 0.88 M aqueous ammonia and (B) acetonitrile. Gradient conditions were initially 1% B, increasing linearly to 97% B over 1.5 min, remaining at 97% B for 0.4 min then increasing to 100% B over 0.1 min. Mass directed automatic purification (MDAP): **High pH MDAP:** The HPLC analysis was conducted on an Xselect CSH C18 column (150 mm x 30 mm i.d. 5 μm packing diameter) at ambient temperature, eluting with 10 mM ammonium bicarbonate in water adjusted to pH 10 with ammonia solution (solvent A) and acetonitrile (solvent B) using an elution gradient of between 0 and 100% solvent B over 15 or 25 min. The UV detection was an averaged signal from wavelength of 210 nm to 350 nm. The mass spectra were recorded on a Waters ZQ Mass Spectrometer using alternate-scan positive and negative electrospray. Ionisation data was rounded to the nearest integer.

### GSK761

#### N-(3-(2-(*tert*-Butoxy)ethyl)phenyl)-4-formylbenzamide

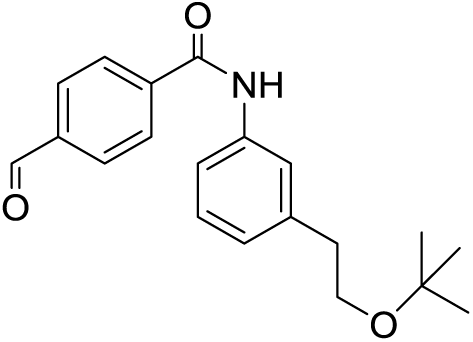

4-Carboxybenzaldehyde (2.3g, 15.32 mmol) was dissolved in Dichloromethane (DCM) (25 mL), then oxalyl chloride (5.83 g, 46.0 mmol) and N,N’-dimethylformamide (DMF) (0.024 mL, 0.306 mmol) were added and the mixture stirred till a colorless solution was obtained. This was evaporated *in vacuo* and the residue redissolved in DCM (25 mL) and cooled in an ice bath. Pyridine (3.72 mL, 46.0 mmol) was added, followed by 3-(2-tertbutoxyethyl)aniline (2.96 g, 15.32 mmol) and the resulting suspension stirred at 0°C to room temperature for 1h.

The reaction mixture was diluted with DCM (50 mL), then washed with water (50 mL) and brine (50 mL), dried and evaporated *in vacuo* to a dark brown gum. This was dissolved in DCM (10 mL) and loaded onto a 100 g silica column, then eluted with 0-50% EtOAc/cyclohexane to give *N*-(3-(2-(*tert*-butoxy)ethyl)phenyl)-4-formylbenzamide (2.75 g, 8.45 mmol, 55.2 % yield) as an amber gum.

^1^H NMR (CHLOROFORM-d, 400 MHz) δ 10.13 (s, 1H), 8.0-8.1 (m, 4H), 7.83 (br s, 1H), 7.5-7.6 (m, 2H), 7.33 (t, 1H, *J*=7.6 Hz), 7.0-7.1 (m, 1H), 3.60 (t, 2H, *J*=7.6 Hz), 2.87 (t, 2H, *J*=7.3 Hz), 1.45 (s, 9H) LCMS (2 min Formic): Rt = 1.12 min, [M -H]^+^ = 324

#### Methyl 2-(4-((3-(2-(*tert*-butoxy)ethyl)phenyl)carbamoyl)phenyl)-1-methyl-1H-benzo[*d*]imidazole-5-carboxylate

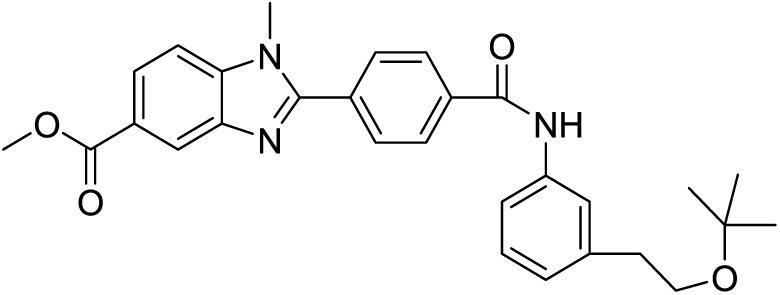

*N*-(3-(2-(*tert*-butoxy)ethyl)phenyl)-4-formylbenzamide (2.6 g, 7.99 mmol) and methyl 4-(methylamino)-3-nitrobenzoate (1.679 g, 7.99 mmol) were combined in ethanol (50 mL), then sodium dithionite (2.78 g, 15.98 mmol) in water (20 mL) was added and the mixture was stirred at 70°C overnight. The mixture was cooled and evaporated *in vacuo* to half volume, then extracted with EtOAc (2 ⨯50 mL) and the organic layer dried and evaporated *in vacuo* to give a pale yellow solid. The solid was dissolved in DCM (10 mL) and loaded onto a 100 g silica column, then eluted with 0-80% EtOAc/cyclohexane and product-containing fractions evaporated *in vacuo* to give methyl 2-(4-((3-(2-(*tert*-butoxy)ethyl)phenyl)carbamoyl)phenyl)-1-methyl-1H-benzo[*d*]imidazole-5-carboxylate (2.35 g, 4.84 mmol, 60.6 % yield)

^1^H NMR (CHLOROFORM-d, 400 MHz) δ 8.55 (s, 1H), 8.18 (s, 1H), 8.11 (dd, 1H, *J*=1.0, 8.6 Hz), 8.04 (d, 2H, *J*=8.1 Hz), 7.89 (d, 2H, *J*=8.1 Hz), 7.5-7.6 (m, 2H), 7.46 (d, 1H, *J*=8.6 Hz), 7.33 (t, 1H, *J*=7.6 Hz), 7.09 (d, 1H, *J*=7.6 Hz), 3.98 (s, 3H), 3.93 (s, 3H), 3.60 (t, 2H, *J*=7.6 Hz), 2.87 (t, 2H, *J*=7.3 Hz), 1.20 (s, 9H)

LCMS (2 min Formic): Rt = 1.14 min, [MH]^+^ = 486

#### 2-(4-((3-(2-(*tert*-Butoxy)ethyl)phenyl)carbamoyl)phenyl)-1-methyl-1H-benzo[*d*]imidazole-5-carboxylic acid

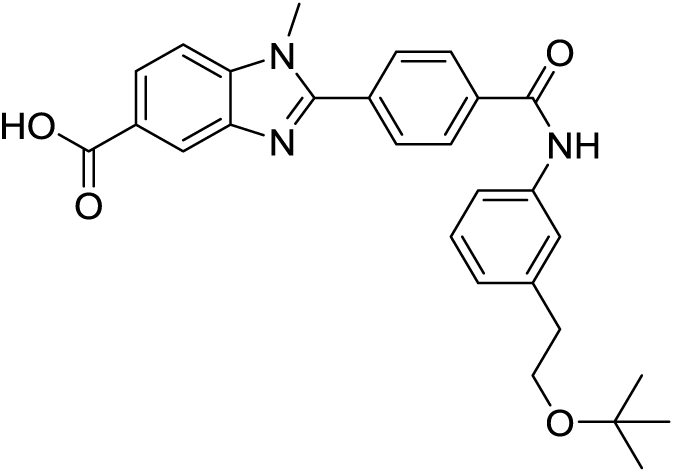

Methyl 2-(4-((3-(2-(*tert*-butoxy)ethyl)phenyl)carbamoyl)phenyl)-1-methyl-1H-benzo[*d*]imidazole-5-carboxylate (2.3 g, 4.74 mmol) was dissolved in THF (10 mL), then LiOH (0.340 g, 14.21 mmol) in water (5 mL) was added and the solution was heated at 60°Cfor 2h, then cooled to room temperature and evaporated *in vacuo*. The residue was dissolved in water (20 mL) and acidified with 2M HCl to pH 4, then the resulting beige solid collected by filtration and the solid dried in the vacuum oven overnight at 60°C to give 2-(4-((3-(2-(*tert*-butoxy)ethyl)phenyl)carbamoyl)phenyl)-1-methyl-1H-benzo[*d*]imidazole-5-carboxylic acid (2.2 g, 4.67 mmol, 98 % yield).

^1^H NMR (DMSO-d_6_, 400 MHz) δ 10.40 (s, 1H), 8.33 (d, 1H, *J*=1.0 Hz), 8.20 (d, 2H, *J*=8.6 Hz), 8.08 (d, 2H, *J*=8.6 Hz), 8.02 (dd, 1H, *J*=1.0, 8.6 Hz), 7.86 (d, 1H, *J*=8.6 Hz), 7.6-7.7 (m, 2H), 7.28 (t, 1H, *J*=7.8 Hz), 7.02 (d, 1H, *J*=7.6 Hz), 4.01 (s, 4H), 3.54 (t, 2H, *J*=7.3 Hz), 2.75 (t, 2H, *J*=7.1 Hz), 1.14 (s, 9H)

LCMS (2 min Formic): Rt = 1.02 min, [MH]^+^ = 472

#### *N*-(3-(2-(*tert*-Butoxy)ethyl)phenyl)-2-(4-((3-(2-(*tert*-butoxy)ethyl)phenyl)carbamoyl)phenyl)-1-methyl-1H-benzo[*d*]imidazole-5-carboxamide

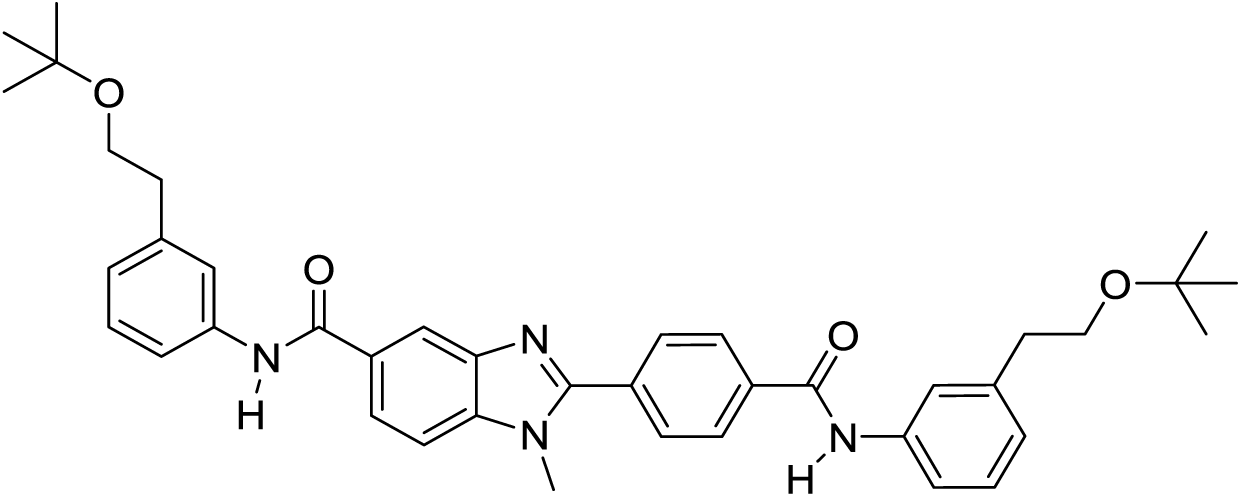

To a mixture of 2-(4-((3-(2-(*tert*-butoxy)ethyl)phenyl)carbamoyl)phenyl)-1-methyl-1H-benzo[*d*]imidazole-5-carboxylic acid (100 mg, 0.212 mmol) and 3-(2-(tert-butoxy)ethyl)aniline (62 mg, 0.321 mmol) in DMF (1 mL) were added HATU (121 mg, 0.318 mmol) and DIPEA (0.056 mL, 0.318 mmol) and the reaction mixture was stirred at room temperature for 3 hours. LCMS showed complete consumption of starting material. The solution was then partitioned between EtOAc and water. The organic layer was washed with water (3x 10 mL), dried over magnesium sulphate and evaporated under vacuum. The sample was loaded in DCM and purified on a 10 g silica cartridge eluting with 0-100% EtOAc-cyclohexane. The appropriate fractions were combined and evaporated *in vacuo* to give the required product *N*-(3-(2-(*tert*-butoxy)ethyl)phenyl)-2-(4-((3-(2-(*tert*-butoxy)ethyl)phenyl)carbamoyl)phenyl)-1-methyl-1H-benzo[d]imidazole-5-carboxamide (118 mg, 0.182 mmol, 86 % yield), as a colorless glass.

^1^HNMR (CHLOROFORM-d, 400MHz): δ (ppm) 8.28 (d, *J*=1.3 Hz, 1H), 8.01 - 8.06 (m, 3H), 7.94 - 7.99 (m, 2H), 7.89 (d, *J*=8.3 Hz, 2H), 7.56 (s, 2H), 7.52 - 7.56 (m, 2H), 7.50 (d, *J*=8.6 Hz, 1H), 7.27 - 7.36 (m, 2H), 7.01 - 7.10 (m, 2H), 3.94 (s, 3H), 3.58 (t, *J*=7.5 Hz, 4H), 2.86 (t, *J*=7.5 Hz, 4H), 1.19 (s, 18H)

LCMS (2 min Formic): Rt = 1.30 min, [MH]^+^ = 647

### GSK064

#### 4-((2,2-Dimethyl-4-oxo-3,8,11-trioxa-5-azatridecan-13-yl)amino)-3-nitrobenzoic acid

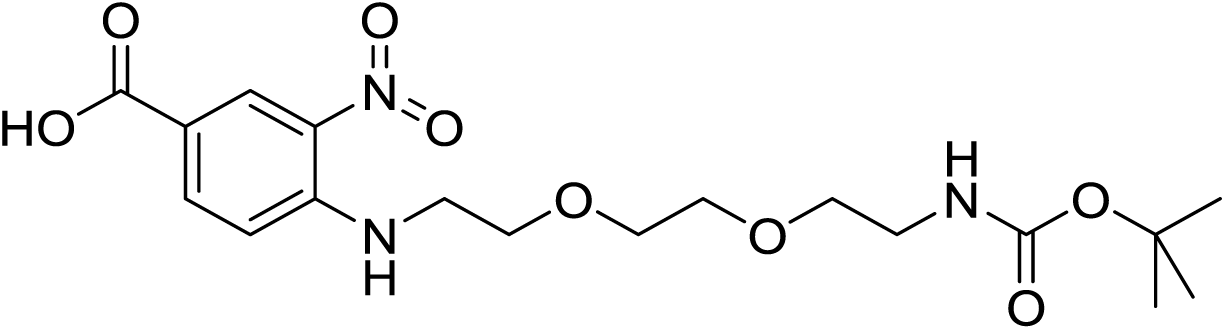

*N*-Boc-2,2’-(ethylenedioxy)diethylamine (1.300 mL, 5.48 mmol) and diisopropylethylamine (2.781 mL, 16.25 mmol) were added to a stirred solution of 4-fluoro-3-nitrobenzoic acid (1.0033 g, 5.42 mmol) in ethanol (15 mL) at ambient temperature. The resulting mixture was stirred under N_2_ atmosphere at 80 °C for 3.5 hr. The solvent was evaporated and dried *in vacuo* to give the crude product, 4-((2,2-dimethyl-4-oxo-3,8,11-trioxa-5-azatridecan-13-yl)amino)-3-nitrobenzoic acid (3.5849 g, 8.67 mmol, 160 % yield) as an orange/yellow oil.

The crude product was used in the next step without further purification.

^1^H NMR (CHLOROFORM-d, 400 MHz) δ 8.92 (d, 1H, *J*=2.0 Hz), 8.43 (br t, 1H, *J*=4.8 Hz), 8.16 (dd, 1H, *J*=1.8, 8.8 Hz), 6.84 (d, 1H, *J*=8.6 Hz), 5.1 (br s, 1H), 3.81 (t, 2H, *J*=5.3 Hz), 3.6-3.7 (m, 4H), 3.5-3.6 (m, 4H), 3.3-3.4 (m, 2H), 1.40 (s, 9H)

LCMS (2 min Formic): Rt = 0.92 min, [MH+]= 414.

#### 3-Amino-4-((2,2-dimethyl-4-oxo-3,8,11-trioxa-5-azatridecan-13-yl)amino)benzoic acid

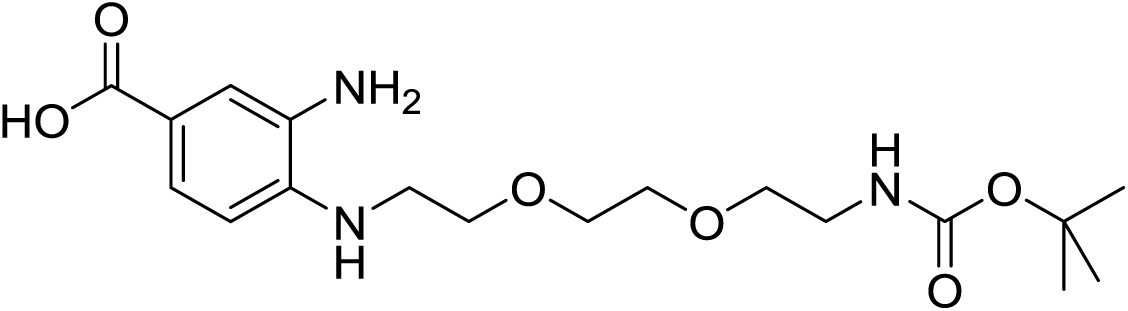

Under N_2_ atmosphere ethanol (10 mL) was to added palladium, 10 wt. % (dry basis) on activated carbon, wet, Degussa type E101 NE/W (0.3776 g, 3.55 mmol) in a hydrogenation flask. A solution of the crude starting material 4-((2,2-dimethyl-4-oxo-3,8,11-trioxa-5-azatridecan-13-yl)amino)-3-nitrobenzoic acid (3.548 g equivalent to 2.241 g, 5.42 mmol) dissolved in ethanol (15 mL) was then added to the catalyst mixture under vacuum. The resulting mixture was placed under an atmosphere of hydrogen gas at room temperature and pressure and stirred vigorously. Once hydrogen uptake had ceased, the reaction mixture was filtered through Celite™ under nitrogen. The solvent was evaporated from the filtrate *in vacuo* to leave a residue that was dried to give the crude product, 3-amino-4-((2,2-dimethyl-4-oxo-3,8,11-trioxa-5-azatridecan-13-yl)amino)benzoic acid (3.141 g, 8.19 mmol, 151 % yield), as a brown oil, which was used without purification in the next step.

LCMS (2 min Formic): Rt = 0.73 min, [MH+]= 384.

#### 2-(4-Carboxyphenyl)-1-(2,2-dimethyl-4-oxo-3,8,11-trioxa-5-azatridecan-13-yl)-1H-benzo[I]imidazole-5-carboxylic acid

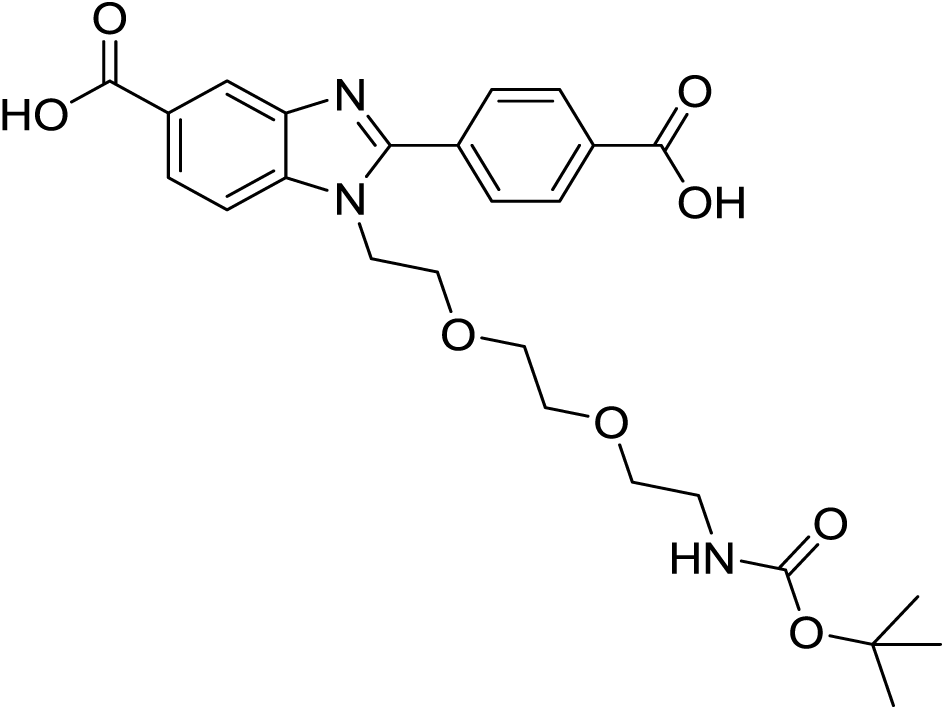

Acetic acid (14 mL) was added to the crude 3-amino-4-((2,2-dimethyl-4-oxo-3,8,11-trioxa-5-azatridecan-13-yl)amino)benzoic acid (2.350 g crude, equivalent to 1.557 g, 4.06 mmol) and 4-carboxybenzaldehyde (0.6122 g, 4.08 mmol). The resulting mixture was stirred under nitrogen at room temperature for 66 hr. The solvent was evaporated *in vacuo* to give a dark reddish-brown oil. The oil was loaded onto a 100 g silica cartridge as a partial suspension in dichloromethane and then purified by silica flash chromatography, eluting with 0-100 % of (1:10:89 acetic acid:MeOH:DCM)/DCM. The required fractions were combined and the solvent evaporated *in vacuo* to give the final product, 2-(4-carboxyphenyl)-1-(2,2-dimethyl-4-oxo-3,8,11-trioxa-5-azatridecan-13-yl)-1H-benzo[I]imidazole-5-carboxylic acid (799.2 mg, 1.323 mmol, 32.6 % yield), as a brown crunchy foam.

^1^H NMR (400 MHz, METHANOL-d_4_) δ 8.62 (s, 1H), 8.43 – 8.37 (m, 2H), 8.26 (dd, *J* = 8.6, 1.5 Hz, 1H), 8.18 – 8.14 (m, 2H), 7.96 – 7.92 (m, 1H), 4.76 – 4.71 (m, 2H), 4.07 – 4.02 (m, 2H), 3.70 – 3.56 (m, 2H), 3.53 – 3.46 (m, 4H), 3.28 – 3.25 (m, 2H), 1.59 (s, 9H)

LCMS (2 min Formic): Rt = 0.78 min, [MH+]= 514

#### *tert*-Butyl (2-(2-(2-(5-((3-(2-(tert-butoxy)ethyl)phenyl)carbamoyl)-2-(4-((3-(2-(*tert*-butoxy)ethyl)phenyl)carbamoyl)phenyl)-1H-benzo[*d*]imidazol-1-yl)ethoxy)ethoxy)ethyl)carbamate

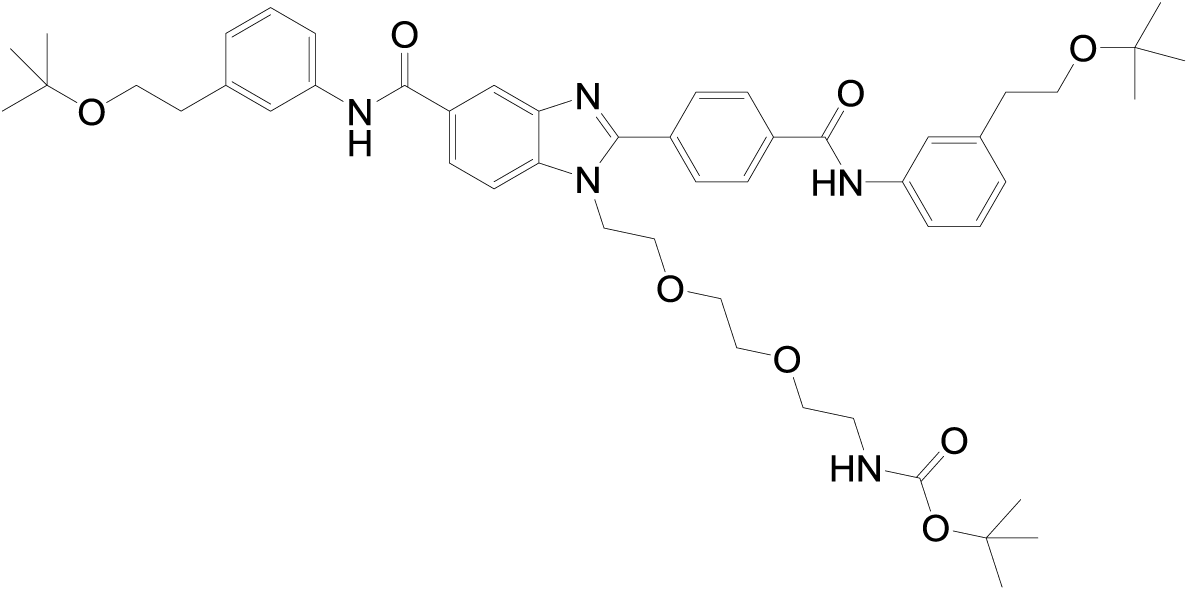

Diisopropylethylamine (0.400 mL, 2.337 mmol) was added to a solution of 3-(2-(*tert*-butoxy)ethyl)aniline (181.3 mg, 0.938 mmol), 2-(4-carboxyphenyl)-1-(2,2-dimethyl-4-oxo-3,8,11-trioxa-5-azatridecan-13-yl)-1H-benzo[*d*]imidazole-5-carboxylic acid (399.4 mg, 0.778 mmol) and HATU (356.7 mg, 0.938 mmol) in DMF (5 mL) and the resulting mixture stirred at room temperature under nitrogen for 150 min. The solvent was evaporated under a stream of nitrogen, citric acid (10 % aq) (50 mL) was added and the aqueous phase extracted with EtOAc (3 x 50 mL). The combined organic phases were filtered through a hydrophobic frit and the solvent evaporated *in vacuo* to give a brown residue. The residue was dissolved in MeOH and loaded onto a 50 g silica cartridge, which was then dried *in vacuo* and eluted with 30-100 % (1 % acetic acid in ethyl acetate)/cyclohexane. The desired product and mono-coupled by-products co-eluted and fractions containing the desired products were combined and the solvent evaporated *in vacuo* to give a pale yellow residue. The resulting residue was dissolved in dichloromethane and loaded on to a 50g silica cartridge, then eluted with 0-10 % MeOH/DCM to give *tert*-butyl (2-(2-(2-(5-((3-(2-(*tert*-butoxy)ethyl)phenyl)carbamoyl)-2-(4-((3-(2-(tert-butoxy)ethyl)phenyl)carbamoyl)phenyl)-1H-benzo[*d*]imidazol-1-yl)ethoxy)ethoxy)ethyl)carbamate (125.7 mg, 0.131 mmol, 16.83 % yield as a pale yellow solid.

^1^H NMR (CHLOROFORM-d, 400 MHz) δ 8.3-8.4 (m, 1H), 7.9-8.2 (m, 6H), 7.6-7.7 (m, 5H), 7.3-7.4 (m, 2H), 7.07 (t, 2H, *J*=7.1 Hz), 4.8-5.0 (m, 1H), 4.4-4.6 (m, 2H), 3.92 (br s, 2H), 3.60 (t, 4H, *J*=7.6 Hz), 3.49 (br s, 4H), 3.3-3.4 (m, 2H), 3.1-3.2 (m, 2H), 2.8-2.9 (m, 4H), 1.43 (s, 9H), 1.21 (s, 18H)

LCMS (2 min Formic): Rt = 1.40 min, [MH+] = 864.

#### 1-(2-(2-(2-Aminoethoxy)ethoxy)ethyl)-*N*-(3-(2-(*tert*-butoxy)ethyl)phenyl)-2-(4-((3-(2-(*tert*-butoxy)ethyl)phenyl)carbamoyl)phenyl)-1H-benzo[*d*]imidazole-5-carboxamide

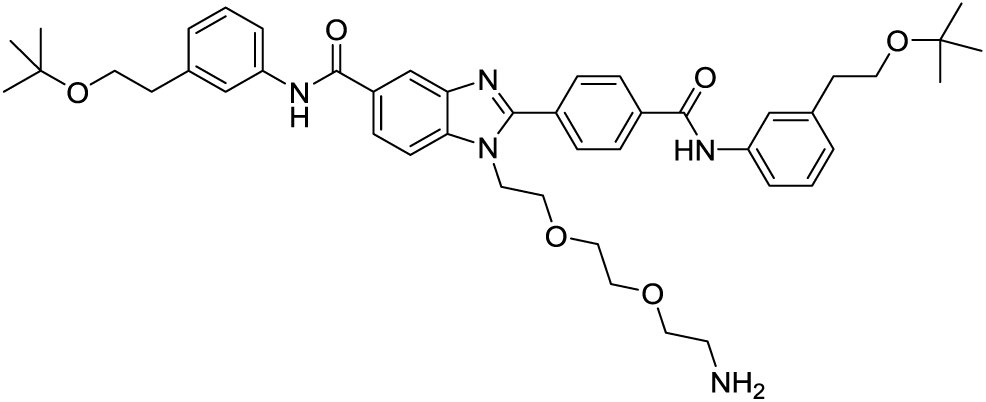

Hydrogen chloride solution (4.0 M in dioxane, 2 mL, 8.00 mmol) was added to *tert*-butyl (2-(2-(2-(5-((3-(2-(*tert*-butoxy)ethyl)phenyl)carbamoyl)-2-(4-((3-(2-(*tert*-butoxy)ethyl)phenyl)carbamoyl)phenyl)-1H-benzo[*d*]imidazol-1-yl)ethoxy)ethoxy)ethyl)carbamate (112.1 mg, 0.130 mmol) and the resulting mixture stirred for 2 min at room temperature. The reaction was quenched with saturated sodium carbonate solution (2 mL), then water (3 mL) and EtOAc (5 mL) were added and the phases separated. The aqueous phase was extracted with further ethyl acetate (2 x 5 mL) and the combined organic phases were filtered through a hydrophobic frit and the solvent evaporated. The resulting residue was dissolved in methanol (1 mL) and purified by high pH MDAP to give 1-(2-(2-(2-aminoethoxy)ethoxy)ethyl)-N-(3-(2-(*tert*-butoxy)ethyl)phenyl)-2-(4-((3-(2-(*tert*-butoxy)ethyl)phenyl)carbamoyl)phenyl)-1H-benzo[*d*]imidazole-5-carboxamide (46.1 mg, 0.057 mmol, 44.2 % yield), was obtained as a very pale yellow glass.

^1^H NMR (CHLOROFORM-d, 400 MHz) δ 8.72 (s, 1H), 8.3-8.4 (m, 2H), 8.0-8.1 (m, 4H), 7.95 (dd, 1H, *J*=1.8, 8.3 Hz), 7.5-7.6 (m, 4H), 7.32 (dt, 2H, *J*=3.5, 7.8 Hz), 7.07 (t, 2H, *J*=7.8 Hz), 4.51 (t, 2H, *J*=5.3 Hz), 3.91 (t, 3H, *J*=5.1 Hz), 3.60 (t, 6H, *J*=7.3 Hz), 3.5-3.6 (m, 4H), 3.39 (t, 2H, *J*=5.3 Hz), 2.88 (t, 4H, *J*=7.6 Hz), 2.80 (t, 2H, *J*=5.3 Hz), 1.21 (s, 18H)

LCMS (2 min high pH): Rt = 1.27 min, [MH+]= 764

#### 3-(2-((1E,3E,5Z)-5-(3-(6-((2-(2-(2-(5-((3-(2-(*tert*-Butoxy)ethyl)phenyl)carbamoyl)-2-(4-((3-(2(*tert*-butoxy)ethyl)phenyl)carbamoyl)phenyl)-1H-benzo[*d*]imidazol-1yl)ethoxy)ethoxy)ethyl)amino)6-oxohexyl)-3-methyl-5-sulfo-1-(3-sulfopropyl)indolin-2-ylidene)penta-1,3-dien-1-yl)-3,3-dimethyl-5-sulfo-3H-indol-1-ium-1-yl)propane-1-sulfonate, 3 Ammonia salt

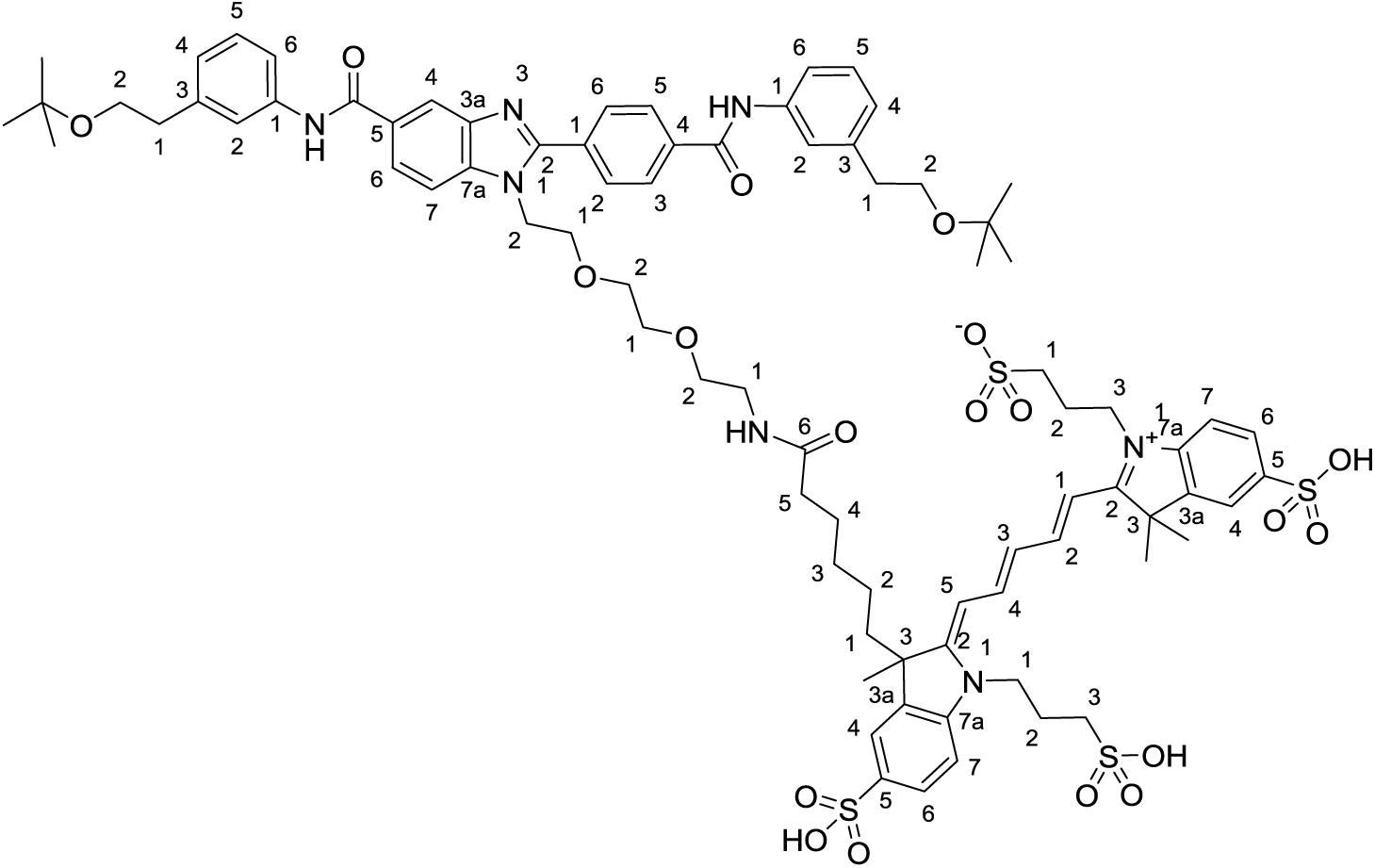

Alexa Fluor 647 carboxylic acid succinimidyl ester (5mg) (Invitrogen) on 15th July 2011; Lot 890799; cat# A20106; mf not given; Exact Mass: not given; Molecular Weight: not given but cited as “∼1250”.

Structure of AlexaFluor 647 determined from WO02/26891 Compound 9. This structure contains two pendant propyl sulfonic acids contrary to some reports suggesting that the molecule contains butyl sulfonic acids. Salt form from patent example of tris(triethylamine) has MWt. 1259.67 which approximates to the vendor’s citation of “∼1250”.

3-(2-((1E,3E,5Z)-5-(3-(6-((2,5-dioxopyrrolidin-1-yl)oxy)-6-oxohexyl)-3-methyl-5-sulfo-1-(3-sulfopropyl)indolin-2-ylidene)penta-1,3-dien-1-yl)-3,3-dimethyl-5-sulfo-3H-indol-1-ium-1-yl)propane-1-sulfonate, 3 N,N-diethylethanamine salt (5 mg, 3.97 µmol) in anhydrous DMF (200 µL) was added to 1-(2-(2-(2-aminoethoxy)ethoxy)ethyl)-N-(3-(2-(tert-butoxy)ethyl)phenyl)-2-(4-((3-(2-(tert-butoxy)ethyl)phenyl)carbamoyl)phenyl)-1H-benzo[d]imidazole-5-carboxamide (5.4 mg, 7.07 µmol). Diisopropylethylamine (1.4 µL, 8.02 µmol) was added and the resulting mixture stirred at room temperature in a vial wrapped in foil for 16 hr. Acetonitrile was added to increase the sample volume to 1 mL. The sample was then purified by high pH MDAP to give 3-(2-((1E,3E,5Z)-5-(3-(6-((2-(2-(2-(5-((3-(2-(*tert*-butoxy)ethyl)phenyl)carbamoyl)-2-(4-((3-(2-(*tert*-butoxy)ethyl)phenyl)carbamoyl)phenyl)-1H-benzo[*d*]imidazol-1-yl)ethoxy)ethoxy)ethyl)amino)-6-oxohexyl)-3-methyl-5-sulfo-1-(3-sulfopropyl)indolin-2-ylidene)penta-1,3-dien-1-yl)-3,3-dimethyl-5-sulfo-3H-indol-1-ium-1-yl)propane-1-sulfonate, 3 Ammonia salt (6.7 mg, 3.84 µmol, 97 % yield), as a blue solid.

^1^H NMR (METHANOL-d_4_, 400 MHz) δ 8.2-8.5 (m, 3H), 8.17 (d, 2H, *J*=8.6 Hz), 8.0-8.1 (m, 3H), 7.9-7.9 (m, 3H), 7.8-7.9 (m, 2H), 7.6-7.7 (m, 4H), 7.43 (dd, 2H, *J*=8.3, 10.9 Hz), 7.31 (t, 2H, *J*=7.8 Hz), 7.07 (dd, 2H, *J*=3.8, 7.3 Hz), 6.6-6.8 (m, 1H), 6.45 (br dd, 2H, *J*=5.3, 13.4 Hz), 4.57 (t, 2H, *J*=4.8 Hz), 4.3-4.4 (m, 4H), 3.88 (t, 2H, *J*=5.1 Hz), 3.65 (dt, 6H, *J*=2.5, 7.1 Hz), 3.42 (br d, 6H, *J*=3.5 Hz), 3.1-3.2 (m, 2H), 2.9-3.1 (m, 5H), 2.8-2.9 (m, 5H), 2.2-2.3 (m, 5H), 1.92 (s, 2H), 1.73 (s, 4H), 1.70 (s, 7H), 1.4-1.5 (m, 2H), 1.17 (s, 18H), 1.1-1.2 (m, 1H), 0.83 (br d, 1H, *J*=3.0 Hz), 0.5-0.7 (m, 1H)

LCMS (2 min high pH): Rt = 0.88 min, [M+2H]2+= 804

### GSK675

#### *N*-(3-(2-(tert-Butoxy)ethyl)phenyl)-4-chloro-3-nitrobenzamide

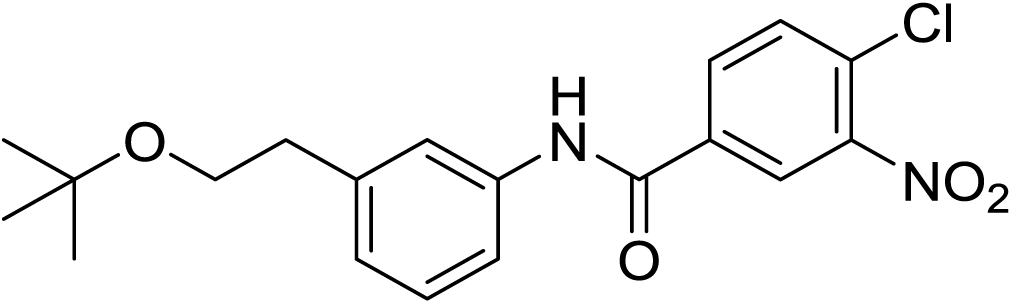

4-Chloro-3-nitrobenzoic acid (4194.8 mg, 20.81 mmol) was taken up into DCM (100 mL) in an ice-bath under nitrogen. Oxalyl dichloride (5.45 mL, 62.4 mmol) was slowly added to the mixture followed by DMF (2 drops), at which point the mixture started to slightly bubble. The reaction was then allowed to stir at RT for 2.5h then the volatiles removed *in vacuo*.

The crude acid chloride was taken up into DCM (100 mL) in an ice-bath under nitrogen. 3-(2-(*tert*-butoxy)ethyl)aniline (4023 mg, 20.81 mmol) was slowly added followed by pyridine (3.36 mL, 41.6 mmol). The reaction was allowed to stir at RT for 2h, then evaporated *in vacuo* to give a brown thick oil. The crude was dissolved in DCM (40 mL) and washed with saturated aq sodium bicarbonate solution (2 x 40 mL) then with hydrochloric acid (1M aq, 40 mL), dried with magnesium sulfate and filtered under vacuum. The solvent was evaporated under vacuum to give *N*-(3-(2-(*tert*-butoxy)ethyl)phenyl)-4-chloro-3-nitrobenzamide (8132.3 mg, 20.50 mmol, 99 % yield) as a brown oil.

^1^H NMR (DMSO-d_6_, 400 MHz) δ 10.46 (s, 1H), 8.64 (d, 1H, *J*=2.3 Hz), 8.27 (dd, 1H, *J*=2.1, 8.5 Hz), 7.97 (d, 1H, *J*=8.6 Hz), 7.6-7.7 (m, 2H), 7.2-7.4 (m, 1H), 7.03 (d, 1H, *J*=7.6 Hz), 3.52 (t, 2H, *J*=7.1 Hz), 2.74 (t, 2H, *J*=7.1 Hz), 1.12 (s, 9H)

LCMS (2 min Formic): Rt = 1.28 min, [M - H]^+^ = 375, 377 (1 Cl)

#### 4-((2-((2-Aminoethyl)(methyl)amino)ethyl)amino)-N-(3-(2-(tert-butoxy)ethyl)phenyl)-3-nitrobenzamide

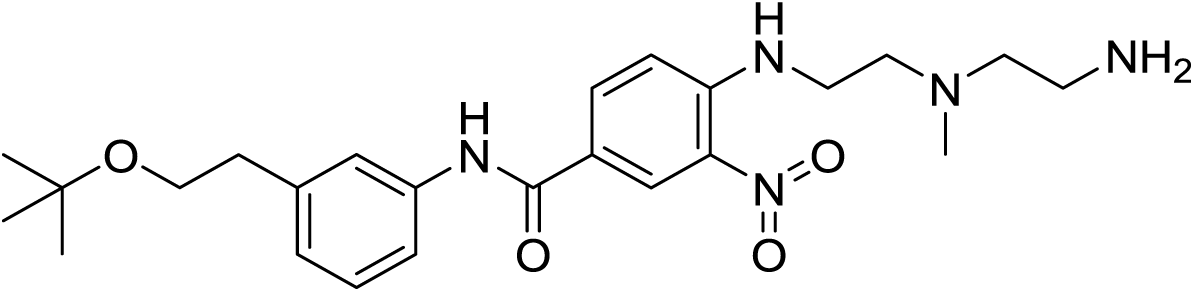

N-(3-(2-(*tert*-Butoxy)ethyl)phenyl)-4-chloro-3-nitrobenzamide (1.03g, 2.73 mmol) and N1-(2-aminoethyl)-*N*1-methylethane-1,2-diamine (0.480 g, 4.10 mmol) were stirred in DMF (10 mL) and triethylamine (0.381 mL, 2.73 mmol) for 3h at 50°C, then the solution was cooled and diluted with water (50ml), extracted with EtOAc (2 x 50 mL) and the organic layer washed with water (2 x 50 mL), dried and evaporated *in vacuo*. The residue was loaded onto a 50 g silica column and eluted with 0-20% 2 M methanolic ammonia/DCM to give 4-((2-((2-aminoethyl)(methyl)amino)ethyl)amino)-*N*-(3-(2-(*tert*-butoxy)ethyl)phenyl)-3-nitrobenzamide (0.85g, 1.858 mmol, 68.0 % yield) as a bright yellow gum

^1^H NMR (CHLOROFORM-d, 400 MHz) δ 8.7-8.8 (m, 1H), 8.67 (d, 1H, *J*=2.5 Hz), 8.52 (s, 1H), 8.0-8.1 (m, 1H), 7.5-7.6 (m, 2H), 7.2-7.3 (m, 1H), 7.00 (d, 1H, *J*=7.6 Hz), 6.85 (d, 1H, *J*=9.1 Hz), 3.56 (t, 2H, *J*=7.3 Hz), 3.38 (q, 2H, *J*=5.6 Hz), 2.8-2.9 (m, 4H), 2.7-2.8 (m, 2H), 2.4-2.6 (m, 2H), 2.28 (s, 3H), 1.17 (s, 9H)

LCMS (2 min high pH): Rt = 1.10 min, [MH]^+^ = 458

#### *tert*-Butyl (2-((2-((4-((3-(2-(tert-butoxy)ethyl)phenyl)carbamoyl)-2-nitrophenyl)amino)ethyl)(methyl)amino)ethyl)carbamate

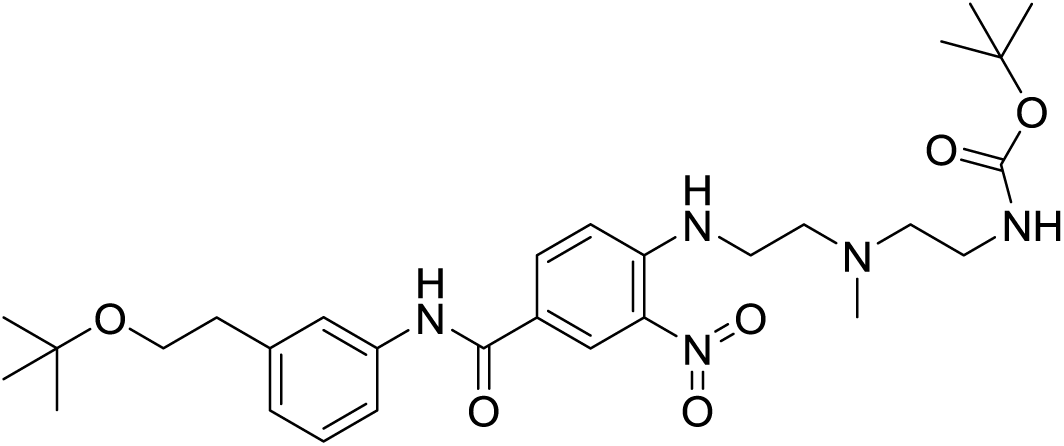

4-((2-((2-Aminoethyl)(methyl)amino)ethyl)amino)-*N*-(3-(2-(tert-butoxy)ethyl)phenyl)-3-nitrobenzamide (0.84g, 1.836 mmol) and Boc_2_O (0.426 mL, 1.836 mmol) were dissolved in DCM (20 mL) and allowed to stand for 2h, then the solvent was evaporated *in vacuo* and the residue was purified by chromatography on a 50 g silica column eluting with 0-10% 2 M methanolic ammonia/DCM to give *tert*-butyl (2-((2-((4-((3-(2-(tert-butoxy)ethyl)phenyl)carbamoyl)-2-nitrophenyl)amino)ethyl)(methyl)amino)ethyl)carbamate (0.85g, 1.524 mmol, 83 % yield) as a bright yellow gum.

^1^H NMR (CHLOROFORM-d, 400 MHz) δ 8.7-8.9 (m, 1H), 8.69 (br s, 1H), 7.9-8.2 (m, 2H), 7.5-7.6 (m, 2H), 7.2-7.4 (m, 1H), 7.03 (d, 1H, *J*=7.6 Hz), 6.90 (d, 1H, *J*=9.1 Hz), 5.10 (br s, 1H), 3.58 (t, 2H, *J*=7.3 Hz), 3.40 (q, 2H, *J*=5.6 Hz), 3.27 (q, 2H, *J*=5.6 Hz), 2.85 (t, 2H, *J*=7.6 Hz), 2.77 (t, 2H, *J*=5.8 Hz), 2.58 (t, 2H, *J*=5.8 Hz), 2.34 (s, 3H), 1.40 (s, 9H), 1.19 (s, 9H)

LCMS (2 min high pH): Rt = 1.41 min, [MH]^+^ = 558

#### 1-(2-((2-Aminoethyl)(methyl)amino)ethyl)-*N*-(3-(2-(*tert*-butoxy)ethyl)phenyl)-2-(4-((3-(2-(*tert*-butoxy)ethyl)phenyl)carbamoyl)phenyl)-1H-benzo[*d*]imidazole-5-carboxamide

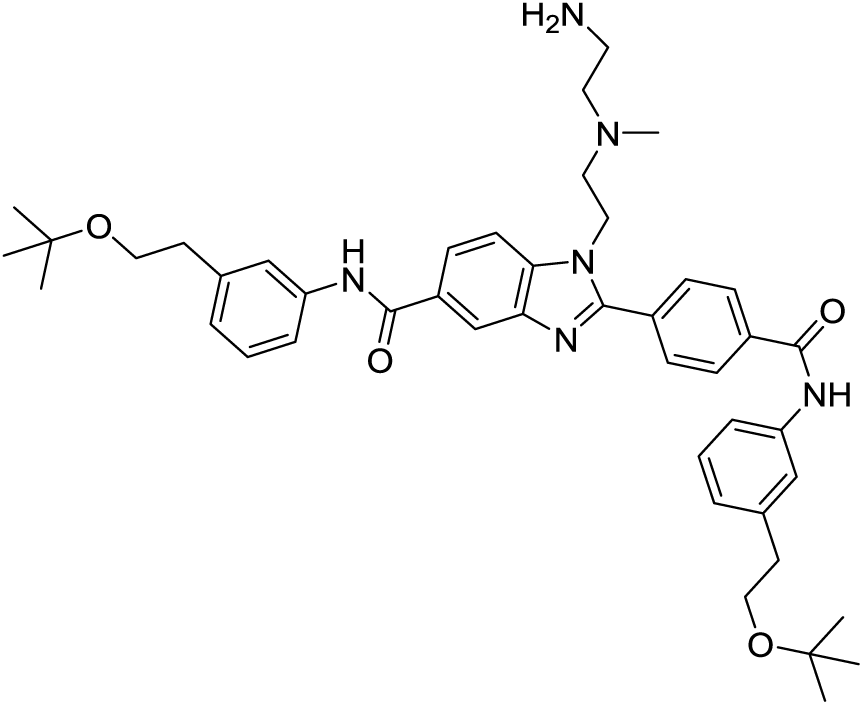

*tert*-Butyl (2-((2-((4-((3-(2-(*tert*-butoxy)ethyl)phenyl)carbamoyl)-2-nitrophenyl)amino)ethyl)(methyl)amino)ethyl)carbamate (0.21g, 0.377 mmol) and *N*-(3-(2-(*tert*-butoxy)ethyl)phenyl)-4-formylbenzamide (0.123 g, 0.377 mmol) were dissolved in ethanol (10 mL), then a solution of sodium dithionite (0.197 g, 1.130 mmol) in water (3 mL) was added and the mixture was heated at reflux for 3h. The solvent was evaporated *in vacuo* and the residue was partitioned between EtOAc (10 mL) and water (5 mL), the organic layer dried and evaporated to give a pale yellow gum.

The crude product was dissolved in DCM (3 mL) and HCl (1M in ether, 1 mL, 1.000 mmol) was added, the mixture stirred for 2h, then evaporated *in vacuo*. A sample (20mg) of the crude product was purified by high pH MDAP to give 1-(2-((2-aminoethyl)(methyl)amino)ethyl)-N-(3-(2-(*tert*-butoxy)ethyl)phenyl)-2-(4-((3-(2-(*tert*-butoxy)ethyl)phenyl)carbamoyl)phenyl)-1H-benzo[*d*]imidazole-5-carboxamide (10mg, 0.014 mmol, 3.62 % yield) as a pale yellow gum.

The remaining crude amine was purified by chromatography on silica (25 g column) eluting with 0-20% 2 M methanolic ammonia/DCM to give 1-(2-((2-aminoethyl)(methyl)amino)ethyl)-N-(3-(2-(*tert*-butoxy)ethyl)phenyl)-2-(4-((3-(2-(*tert*-butoxy)ethyl)phenyl)carbamoyl)phenyl)-1H-benzo[*d*]imidazole-5-carboxamide (85 mg, 0.116 mmol, 30.8 % yield) as a colorless foam.

^1^H NMR (CHLOROFORM-d, 400 MHz) δ 8.48 (s, 1H), 8.27 (s, 1H), 8.14 (s, 1H), 8.03 (d, 3H, *J*=8.1 Hz), 7.96 (dd, 1H, *J*=1.5, 8.6 Hz), 7.87 (d, 3H, *J*=8.6 Hz), 7.5-7.6 (m, 7H), 7.3-7.4 (m, 3H), 7.07 (t, 2H, *J*=8.8 Hz), 4.40 (br t, 2H, *J*=6.6 Hz), 3.60 (dt, 5H, *J*=1.8, 7.5 Hz), 2.87 (t, 5H, *J*=7.3 Hz), 2.71 (t, 2H, *J*=6.3 Hz), 2.4-2.5 (m, 2H), 2.3-2.4 (m, 2H), 2.09 (s, 3H), 1.20 (s, 18H)

LCMS (2 min high pH): Rt = 1.27 min, [MH]^+^ = 733

#### *N*-(3-(2-(*tert*-Butoxy)ethyl)phenyl)-2-(4-((3-(2-(*tert*-butoxy)ethyl)phenyl)carbamoyl)phenyl)-1-(2-(methyl(2-(6-(5-((3aS,4S,6aR)-2-oxohexahydro-1H-thieno[3,4-d]imidazol-4-yl)pentanamido)hexanamido)ethyl)amino)ethyl)-1H-benzo[*d*]imidazole-5-carboxamide

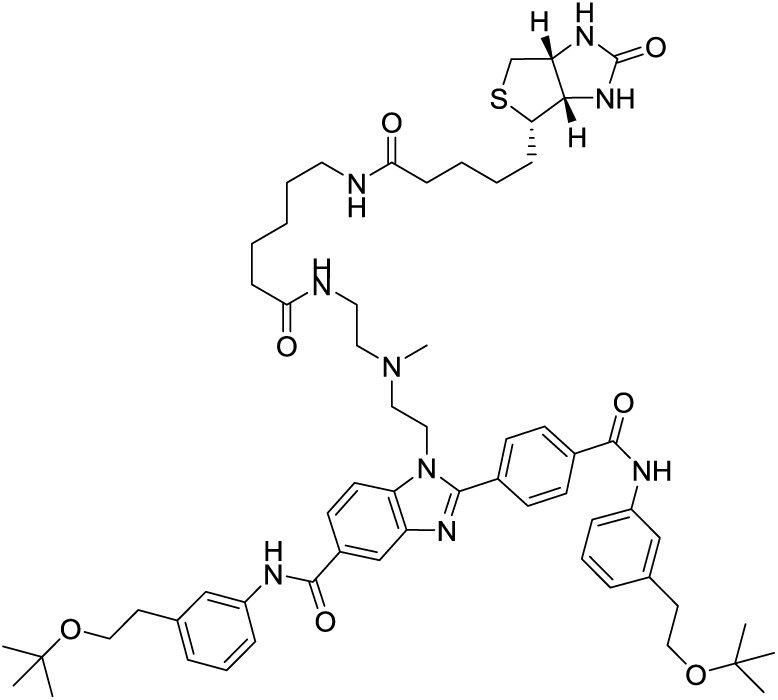

1-(2-((2-Aminoethyl)(methyl)amino)ethyl)-N-(3-(2-(*tert*-butoxy)ethyl)phenyl)-2-(4-((3-(2-(*tert*-butoxy)ethyl)phenyl)carbamoyl)phenyl)-1H-benzo[*d*]imidazole-5-carboxamide (15 mg, 0.020 mmol) and 2,5-dioxopyrrolidin-1-yl 6-(5-((3aS,4S,6aR)-2-oxohexahydro-1H-thieno[3,4-d]imidazol-4-yl)pentanamido)hexanoate (9.30 mg, 0.020 mmol) were dissolved in DMF (0.9 mL) and triethylamine (8.56 µl, 0.061 mmol) and the solution was allowed to stand overnight. LCMS (2 min high pH): Rt = 1.21 min, [MH]^+^ = 1072. The solution was purified by high pH MDAP to give *N*-(3-(2-(*tert*-butoxy)ethyl)phenyl)-2-(4-((3-(2-(*tert*-butoxy)ethyl)phenyl)carbamoyl)phenyl)-1-(2-(methyl(2-(6-(5-((3aS,4S,6aR)-2-oxohexahydro-1H-thieno[3,4-d]imidazol-4-yl)pentanamido)hexanamido)ethyl)amino)ethyl)-1H-benzo[*d*]imidazole-5-carboxamide (8.5 mg, 7.93 µmol, 38.7 % yield) as a colorless solid.

^1^H NMR (METHANOL-d_4_, 400 MHz) δ 8.39 (d, 1H, *J*=1.5 Hz), 8.19 (d, 2H, *J*=8.6 Hz), 8.04 (dd, 1H, *J*=1.5, 8.6 Hz), 8.0-8.0 (m, 2H), 7.83 (d, 1H, *J*=8.6 Hz), 7.5-7.7 (m, 4H), 7.32 (dt, 2H, *J*=3.0, 7.8 Hz), 7.08 (t, 2H, *J*=6.8 Hz), 4.54 (t, 2H, *J*=6.1 Hz), 4.47 (dd, 1H, *J*=5.1, 8.1 Hz), 4.2-4.3 (m, 1H), 3.66 (t, 5H, *J*=7.3 Hz), 3.1-3.2 (m, 1H), 3.00 (t, 2H, *J*=6.6 Hz), 2.8-2.9 (m, 5H), 2.8-2.8 (m, 2H), 2.6-2.7 (m, 1H), 2.37 (t, 2H, *J*=6.3 Hz), 2.17 (t, 2H, *J*=7.3 Hz), 2.06 (t, 2H, *J*=7.6 Hz), 1.4-1.8 (m, 11H), 1.3-1.3 (m, 3H), 1.21 (s, 18H)

LCMS (2 min high pH): Rt = 1.04 min, M/2+H observed 536

### GSK306

#### *N*-(3-(2-(*tert*-Butoxy)ethyl)phenyl)-2-(4-((3-(2-(*tert*-butoxy)ethyl)phenyl)carbamoyl)phenyl)-1-(2-(2-(2-(3’,6’-dihydroxy-3-oxo-3H-spiro[isobenzofuran-1,9’-xanthen]-5-ylcarboxamido)ethoxy)ethoxy)ethyl)-1H-benzo[*d*]imidazole-5-carboxamide

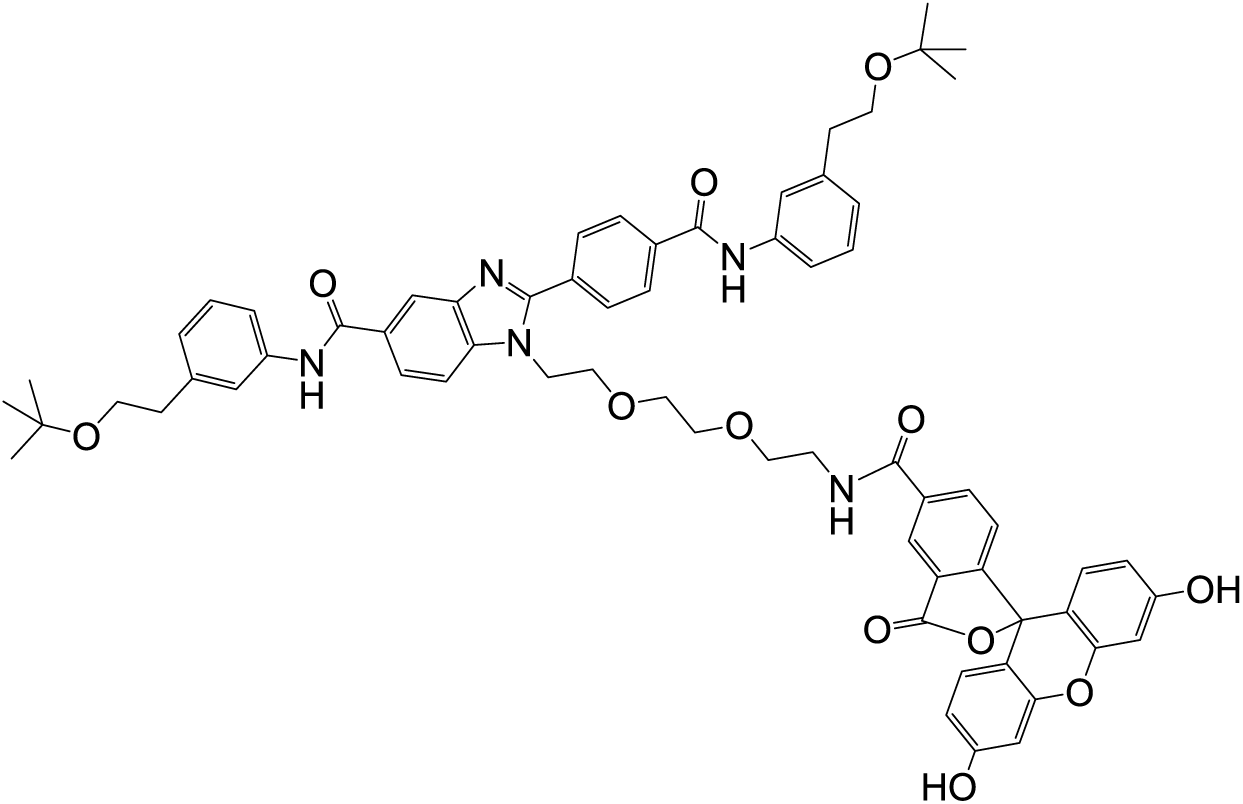

DMF (0.4 mL) and triethylamine (4.42 µL, 0.032 mmol) were added to 5-carboxyfluorescein N-succinimidyl ester (5.1 mg, 10.77 µmol) and 1-(2-(2-(2-aminoethoxy)ethoxy)ethyl)-N-(3-(2-(tert-butoxy)ethyl)phenyl)-2-(4-((3-(2-(tert-butoxy)ethyl)phenyl)carbamoyl)phenyl)-1H-benzo[d]imidazole-5-carboxamide (13.5 mg, 0.018 mmol) and the resulting mixture stirred at room temperature in a vial wrapped in foil for 17.5 hr. The mixture was left to stand overnight, then diluted with methanol to 1 mL and then the sample purified by MDAP (high pH) to give *N*-(3-(2-(*tert*-butoxy)ethyl)phenyl)-2-(4-((3-(2-(tert-butoxy)ethyl)phenyl)carbamoyl)phenyl)-1-(2-(2-(2-(3’,6’-dihydroxy-3-oxo-3H-spiro[isobenzofuran-1,9’-xanthen]-5-ylcarboxamido)ethoxy)ethoxy)ethyl)-1H-benzo[*d*]imidazole-5-carboxamide (6.8 mg, 5.76 µmol, 53.4 % yield), as an orange solid.

^1^H NMR (METHANOL-d_4_, 400 MHz) δ 8.37 (dd, 2H, *J*=1.3, 11.9 Hz), 8.1-8.2 (m, 2H), 8.0-8.1 (m, 3H), 7.98 (dd, 1H, *J*=1.5, 8.6 Hz), 7.80 (d, 1H, *J*=8.6 Hz), 7.6-7.7 (m, 2H), 7.58 (br d, 2H, *J*=7.6 Hz), 7.2-7.3 (m, 2H), 7.21 (d, 1H, *J*=8.1 Hz), 7.0-7.1 (m, 2H), 6.6-6.8 (m, 4H), 6.5-6.6 (m, 2H), 4.57 (br t, 2H, *J*=4.8 Hz), 3.94 (br t, 2H, *J*=5.1 Hz), 3.63 (dt, 4H, *J*=2.0, 7.1 Hz), 3.5-3.6 (m, 8H), 2.81 (br t, 4H, *J*=6.6 Hz), 1.18 (m, 18H) (5 exchangeable protons not seen).

LCMS (2 min high pH): Rt = 0.98 min, [MH+]= 1122

